# Spatial Trophic Dynamics Shape and are Shaped by Desertification Transitions

**DOI:** 10.64898/2026.02.04.703762

**Authors:** Koustav Halder, Juan A. Bonachela

## Abstract

Drylands, which sustain billions of people, face desertification driven by climate change and grazing pressures. From the bottom up, desertification is affected by water availability, with vegetation often self-organising into spatial patterns that vary with aridity levels. However, how these patterns and, ultimately, desertification transitions are affected by the spatial dynamics of higher trophic levels remains poorly understood. Here, we introduce a spatially explicit tri-trophic model that links vegetation pattern formation to consumer-resource interactions and foraging behaviour. We find that the nature of vegetation spatial distribution and desertification transition strongly influence consumer spatial dynamics, which in turn influences desertification. Vegetation organised regularly in space relates to “boom—bust” synchronised metapopulations, whereas scale-free vegetation organisation is associated with scale-free consumer clustering and low synchrony. Our results reveal feedbacks between spatial trophic dynamics and ecosystem resilience, underscoring the need to integrate trophic interactions and behaviour into predictions informing management strategies for dryland ecosystems.

## 1 Introduction

Desertification – the process by which arable, semi-arid landscapes collapse to barren desert – affects 500 million people worldwide, the vast majority based in the drylands of the Global South (Caretta et al. 2022). Drylands account for over two-thirds of the area of seven biodiversity hotspots worldwide (The State of the World’s Forests 2020). Their inhabitants are especially vulnerable to desertification partly due to their socio-ecological dependence on grazing: drylands comprise 78% rangelands and pastures worldwide, with grazers delivering fundamental ecosystem services such as soil fertility and plant community stability (Reynolds et al. 2007a; Maestre et al. 2022). Thus, with climate change accelerating the expansion of desertification hotspots into such fertile rangelands and a projected increase in food insecurity for an estimated three billion people by 2050 (Intergovernmental Panel on Climate Change 2022a; Rosenzweig et al. 2014), understanding the mechanisms that influence desertification is of utmost importance.

Dryland landscapes are often characterised by spatial vegetation patterns, commonly appearing as ‘scale-free’ (i.e. forming clusters whose size shows a power-law distribution) (Scanlon et al. 2007; Kéfi et al. 2014) or ‘regular’ (i.e. forming motifs that are repeated regularly in space) (Rietkerk and van de Koppel 2008) (Fig. 1A-B). These patterns are thought to emerge from different mechanisms: the relative scales of local facilitative interactions and demographic processes favour the emergence of scale-free patterns, whereas longer-range competition for e.g., water availability favours regular patterns (Kéfi et al. 2007; Manor and Shnerb 2008). Because they result from vegetation response to bottom-up and top-down pressures that change in space and time, these patterns capture the current state of the ecosystem more comprehensively than non-spatial features (e.g. total or average vegetation biomass). Importantly, vegetation patterns have been linked to ecosystem resilience, that is, the ability for vegetation to survive and therefore for the ecosystem to avoid desertification for the given environmental conditions (Holling 1973). For example, changes in regular patterns from stripes to clumps have been proposed as *‘Early Warning Signals* (EWS) indicating ecosystem health deterioration, aiding to anticipate the desertification transition (Rietkerk and van de Koppel 2008; Kéfi et al. 2014). Due to the scales at which vegetation in arid ecosystems changes in space and time, however, quantifying and monitoring changes in ecosystem health and resilience requires very long time series that are typically not available empirically (but see (Guttal and Jayaprakash 2008)). Theoretical models are thus an excellent tool to help understand and predict desertification.

**Figure 1:**
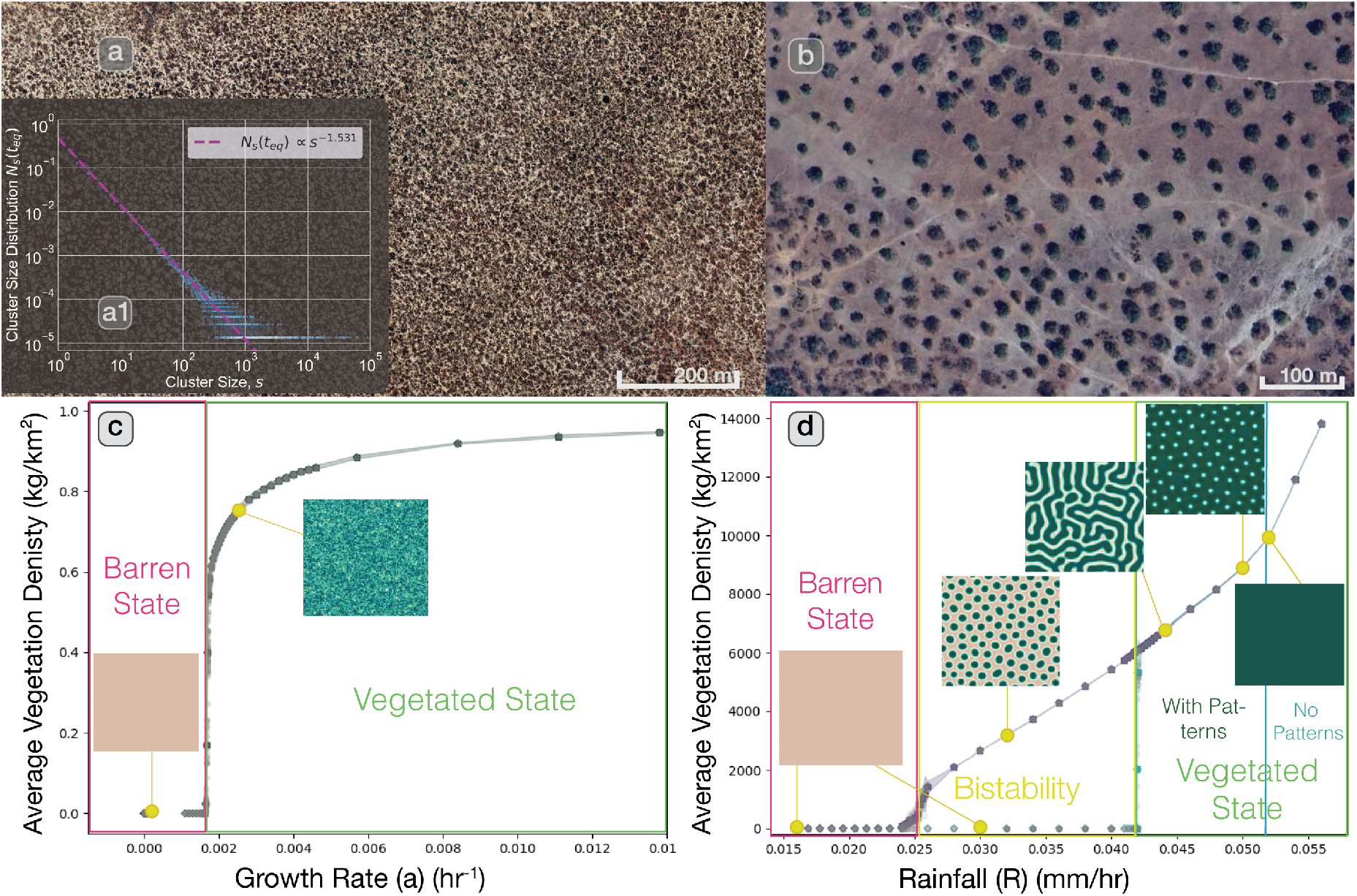
Example of patterned vegetation in drylands, and desertification transition predicted by standard theoretical models used to replicate these patterns. **a** Satellite imagery of scale-free vegetation in the Hwange National Park, Zimbabwe (19.021S, 26.464E) *(image credits: Google Earth)*, characterised by clusters whose sizes follow a power-law probability distribution with an exponent *τ* = 1.531 (inset, **a1**). **b** Satellite imagery of regular vegetation near Marahoué National Park, Côte d’Ivoire (7.248N, 6.108W) *(image credits: Google Earth)*, characterised by a clump-like distribution (Lejeune, Tlidi, and Couteron 2002). **c** Phase diagram corresponding to a scale-free vegetation model (see Eq. (1a)); scale-free vegetation is generally predicted to follow a continuous desertification transition, with desertification occurring at a critical value at which scale-free (i.e. power-law) behaviour emerges for various vegetation-related observables such as the cluster size distribution. **d** Discontinuous transition predicted by a regular-vegetation model (Eqs. (1b)–(1d)), with the barren and vegetated states sharing a region of bi-stability where random fluctuations in environmental drivers may trigger a shift from one state to another. Insets show the spatial vegetation patterns that emerge at various levels of aridity (related to growth rates in panel c and to rainfall levels in panel d).

Until recently, theoretical models focused on the role of bottom-up drivers of desertification, investigating how vegetation patterns and transitions depend on aridity (i.e. water availability, estimated indirectly from vegetation growth rates or directly from rainfall levels). Scale-free patterns are commonly described as emerging from the stochasticity inherent to vegetation birth-death processes, that is, to random fluctuations that are proportional in magnitude to the population density (Villa Martín et al. 2015; Scanlon et al. 2007; Manor and Shnerb 2008). Thus, scale-free patterns result when this demographic stochasticity plays a dominant role in vegetation dynamics. Regular vegetation patterns, on the other hand, are commonly described as emerging from *‘scale-dependent feedbacks’* (SDF) (Rietkerk and van de Koppel 2008): in short, high vegetation density at a location leads to increased soil-water accumulation from rainfall and thus more local vegetation growth, which generates a short-range positive feedback; simultaneously, vegetation at neighbouring locations competes for water, which leads to a long-range negative feedback.

Top-down pressures such as grazing are also important for drylands (see above) (Maestre et al. 2022; Pichon et al. 2025). However, whether the effect of grazing on these ecosystems is positive (e.g. reducing fuel loads, enhancing primary production) or negative (contributing to degradation and desertification) is still unclear. The latter has recently motivated the inclusion of grazing in vegetation models by utilising various approaches that range from a spatially homogeneous constant grazing pressure (Rietkerk and van de Koppel 1997) to more realistic spatially heterogenous grazing (Schneider and Kéfi 2016; Ge and Liu 2022). These models have advanced knowledge on the interaction between grazers and the underlying vegetation pattern-forming mechanisms, and the resulting impact on land-scape resilience (Tarnita et al. 2017; Pichon et al. 2025). One such model has found, for instance, that the interaction between keystone herbivores and heterogenous nutrient patterns influences vegetation spatio-temporal dynamics and boosts overall ecosystem productivity (Vardaro et al. 2021). Another example found that herbivory generally countered the effects of aridity on vegetation spatial structure, leading to a coarsening of vegetation patterns (e.g. overall increase of clump size) (Pichon et al. 2025). Including grazers in a more realistic way thus leads to counterintuitive results since their consumption of vegetation does not necessarily translate into ecosystem degradation or decreased resilience (i.e. higher risk of desertification).

Key gaps remain, however, that prevent the development of reliable predictive theories for desertification. For example, grazer behaviour and movement is typically neglected (or included in a simplistic way (Ge and Liu 2022; Singha, Uecker, and Meron 2025)). Also neglected are predators, which impact pastoralist and wild grazers and often shape their behaviour and foraging dynamics (Gaynor et al. 2019). This is especially true in drylands such as in the East African savannah, where dominant ungulate species modify foraging behaviour and movement to reduce predation risk (Riginos 2015). Whether and how these behavioural changes affect the underlying vegetation pattern or potential trophic cascades is still unknown.

These factors can affect and be affected by vegetation patterns and the desertification transition. Vegetation patterns and dynamics can, for example, affect temporal synchrony (that is, the spatial correlation between populations at different times) arising from foraging dispersal effects (Paradis et al. 1999), or affect the interaction of synchronous populations with other synchronous populations (Lieb-hold, Koenig, and Bjørnstad 2004). Changes in the spatio-temporal distribution of vegetation can also influence how consumer dynamics depend on a synchronising external factor such as rainfall (i.e. “Moran effect”) (Ranta et al. 1997). These correlations, important for biodiversity, ecosystem functioning, and conservation efforts (Guill, Hülsemann, and Klauschies 2021; Grünzweig et al. 2022), are of particular relevance in the desertification context, as inter-trophic synchrony plays a key role in stabilising trophic communities (Visser and Gienapp 2019). Thus, while different pattern-forming mechanisms are expected to impact predator and grazer spatial dynamics and foraging behaviours in different ways (Noonan et al. 2023), these impacts have not been systemically studied.

Here, we attempt to unravel the role of bottom-up and top-down factors shaping dryland communities and the desertification transition. To that end, we introduce a spatially explicit multi-scale model that considers the dynamics of vegetation, grazers, and predators, and that accounts for consumer movement via novel data-informed allometric relationships. With this model, we tackle the following questions: *i)* How do vegetation pattern-formation mechanisms and the desertification transition affect consumer spatial dynamics, foraging behaviour, and spatio-temporal correlations (i.e. synchrony)?, and *ii)* How do such spatial trophic dynamics and behaviour affect the emergent vegetation patterns and the underlying transition? We find that demographic stochasticity induces scale-free vegetation and consumer spatial organisation, but consumers somewhat decrease the resilience of the ecosystem (i.e. desertification occurs for aridity levels for which vegetation by itself would have survived). On the other hand, scale-dependent feedbacks induce regular vegetation and consumer organisation, with emergent vegetation patterns that are coarser and observed for a wider range of environmental conditions than in the absence of consumers. Furthermore, the spatial trophic dynamics show boom-bust cycles that exhibit oscillatory synchrony, and their timing provides EWSs that encode temporal, spatial, and behavioural information.

All together, our results highlight that dryland trophic communities are shaped by intertwined bottom-up and top-down pressures, underlining the need to incorporate trophic dynamics into current understanding of dryland ecosystems and their resilience to desertification.

## 2 Materials and Methods

### 2.1 Models

To represent the spatial dynamics of tri-trophic communities (vegetation, grazer, predator) in a typical semi-arid ecosystem setup, we linked standard dryland vegetation models to an existing consumer-resource model (Brose et al. 2008; Kéfi et al. 2012) that we extended by introducing consumer behaviour and movement. We summarise the main aspects of the models below, but see Supplementary Information and Tables S.1 and S.2 for details.

#### 2.1.1 Vegetation Equations

Our models track vegetation (*V* (*x, t*)), grazer (*G*(*x, t*)) and predator (*Pr*(*x, t*)) biomass densities (in *kg km*^−2^). To represent vegetation dynamics, we considered two models that produce different spatial patterns. The use of two models is justified from an ecological perspective, as the two types of vegetation patterns occur in different types of semi-arid ecosystems, under different conditions, due to different mechanisms, and are associated to different types of desertification transitions (see above). Moreover, our aim is the study of mutual interaction between consumers and these two different mechanisms for pattern formation, rather than contrast these mechanisms against each other in a unifying framework. Finally, from a technical perspective, to the best of our knowledge there is no single model able to produce all three regular patterns observed in semi-arid ecosystems (clumps, labyrinths, and gaps) as well as scale-free patterns.

##### i) Scale-free vegetation (SFV)

the typical model used to describe the dynamics of vegetation that shows a scale-free distribution of cluster sizes (Scanlon et al. 2007; Manor and Shnerb 2008; Kéfi et al. 2007; Villa Martín et al. 2015) does not consider consumers and is based on the so-called Directed Percolation model from physics (Hinrichsen 2000). In the scale-free vegetation (SFV) model, vegetation growth rate is typically the parameter tweaked to study desertification due to its correlation with environmental conditions (e.g. aridity), and patterns and power laws emerge only at the (smooth) transition (critical growth rate *a* = *a*_*c*_, Fig. 1C), with scaling exponents that depend on the macroscopic properties of the system. Here, we modified the original equation to include grazing:

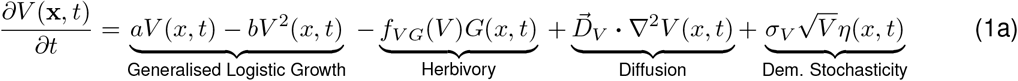

Vegetation grows (first term) limited by intraspecific competition (second term), declines due to grazing (third term), spreads due to dispersal or cloning (fourth term) and shows demographic stochasticity (last term, white noise *η* with variance proportional to vegetation density).

##### ii) Regular vegetation (RV)

we chose a classic model that produces regular, Turing-like vegetation patterns, which emerge due to SDFs (Rietkerk et al. 2004) (Fig. 1D) before the (abrupt) desertification transition. We modified this regular vegetation (RV) model by adding a grazing term:

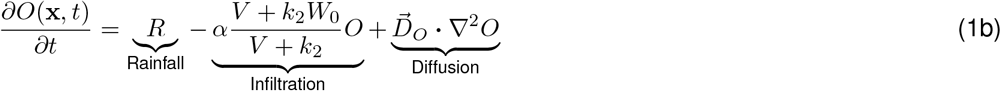

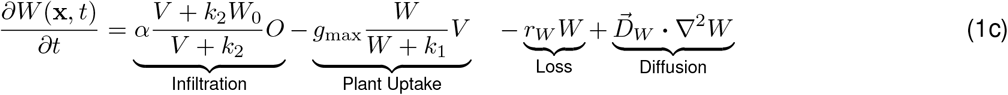

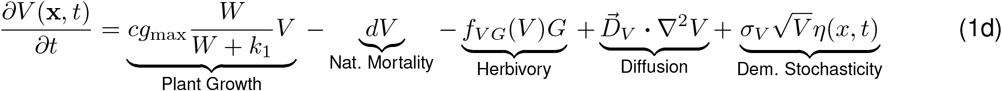

Surface water *O* accumulates due to rainfall (first term, first equation) and infiltrates into soil facilitated by vegetation (second term), and moves non-directionally on the surface (third term). Infiltration increases underground water *W* (first term, second equation), which is reduced via uptake (second term) or evaporation (third term), and that can spread non-directionally on the water table (last term). Vegetation *V* grows thanks to uptake (first term, third equation), is reduced by natural mortality or grazing (second and third term, respectively), spreads non-directionally via plant dispersal or cloning (fourth term), and is subject to demographic stochasticity (last term). See Table S.1 for parameter definition, units, and values.

#### 2.1.2 Consumer Equations

For both models, we added the dynamics for grazer biomass density (*G*(**x**, *t*)):

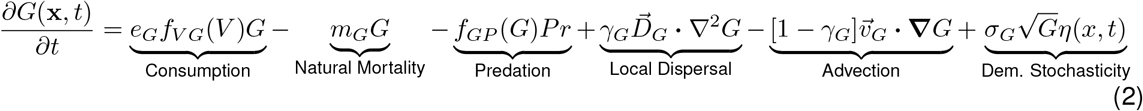

Grazer density increases through herbivory (first term), and decreases via natural mortality (second term) and predation (third term). The fourth and fifth terms represent foraging (see next section), respectively; the last term represents demographic stochasticity.

We added a similar equation for predator density *Pr*(**x**, *t*):

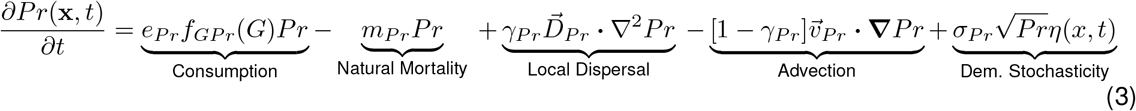

We assumed that consumption functions saturate (Holling 1959):

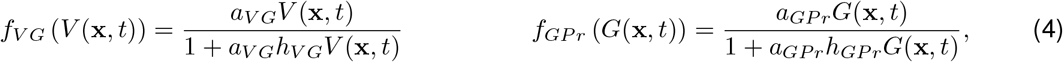

with *a*_*V G*_ and *a*_*GP r*_ representing maximum consumer-resource encounter rates and *h*_*V G*_ and *h*_*GP r*_, handling times (Pawar, Dell, and Van M. Savage 2012).

#### 2.1.3 Consumer Movement and Decision-Making

We considered that consumers at a given site 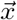 and time *t* may either disperse in a non-directional (i.e. unbiased) random walk or advect in response to cues (away from high consumer densities to reduce intra-specific competition and, for grazers, away from high predator densities; Fig. 2A) (Skalski and Gilliam 2003; Tyson, Wilson, and Lane 2011). Following standard approaches, dispersal is represented using the second spatial derivative (∇^2^) with the area tiled per unit time given by *D* (in *km*^2^ *hr*^−1^), which translates into local random movement; advection at a velocity 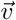 (in *km hr*^−1^) is captured with the first spatial derivative, which translates into decisive movement away from the current location. Since consumers base the decision to relocate on perception radius (see below), we assume that directionality for advection is random as they do not have information beyond their perception area.

**Figure 2:**
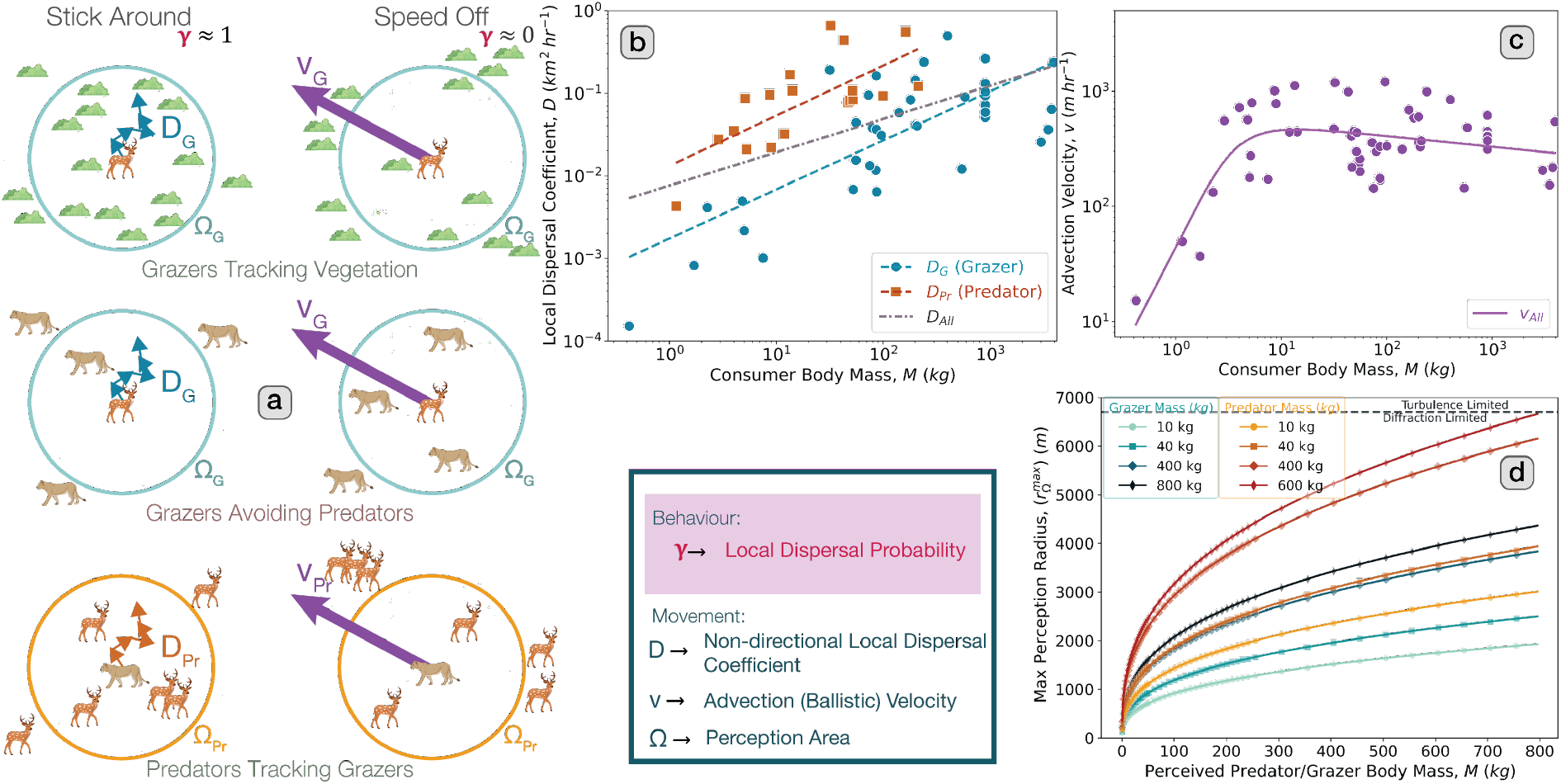
Schematic representation of consumer movement and decision-making, and novel data-informed allometries developed here to inform it. **a** Illustration of consumer foraging behaviour, considered here through the non-directional dispersal probability, *γ*, representing their decision to persist in favourable environments or alternatively advect out of less favourable ones. Consumers track resources within their perception area Ω (top and bottom panels), with grazers also avoiding predators (middle panel). The stronger the cues to leave (e.g. low resource and, for the grazer, high predator density), the lower the value of their dispersal probability *γ* and the higher their affinity for advection. Colours of coefficients *D*_*G*_, *D*_*Pr*_, *v*_*G*_ and *v*_*Pr*_ match colours of their respective allometries in panels b and c. **b**-**c** Allometries deduced here for the non-directional dispersal (diffusion) coefficient 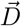 and the advection velocity 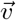 using geo-located movement data from (Noonan et al. 2023). **d** Allometries for the maximum perception radius 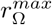 for grazers (cooler palette) and predators (warmer palette) which represents the distance over which an organism of a given bodymass can visually detect the other trophic group (predator and grazer, respectively), plotted as a function of bodymasses of the other trophic group; see SIA for details.

Although empirical work supports advection-dispersal movement (Johnson et al. 2008; Turchin 1998), research relating the underlying decision-making to local cues remains limited (Ge and Liu 2022). Nevertheless, a recent analysis suggests strong links between terrestrial mammal movement strategy and local resource abundance (Noonan et al. 2023). Thus, we incorporated behaviour as decision-making relating movement strategy (stay vs leave) to local resource abundance for both consumers; for grazers, we also added predation pressure which, in addition to being ecologically reasonable, has been documented in semi-arid environments (‘ecology of fear’ (Riginos 2015; Gaynor et al. 2019)). Specifically, we defined the dispersal probability of a consumer, *γ* (Fig. 2A). For the predator, this probability is given by the average proportion of grazer density within the predator’s perception radius Ω_*Pr*_, that is, the ratio “average grazer density within perception radius to global average grazer density ⟨*G*⟩”:

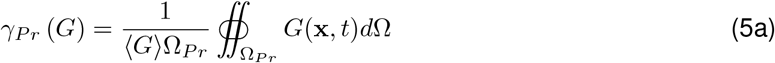

Grazer behaviour is influenced by vegetation density (first term in integral below), but also contemplates predator avoidance by considering the relative densities of predators within their perception area Ω_*G*_:

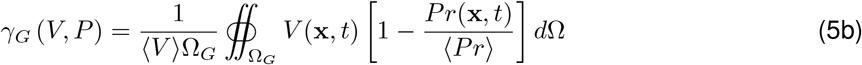

Consumers are thus more likely to disperse locally in a resource-rich region, and advect out of a resource-poor one. Additionally, grazers are more likely to advect out of predator-rich areas and stay in areas with low predatory pressures (Fig. 2A).

Finally, the perception area Ω is defined by its radius *r*_Ω_, in turn limited by vegetation cover: perception is more challenging in systems that have more vegetation cover 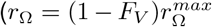, where 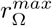 is maximum perception radius and *F*_*V*_, the proportion of sites with vegetation).

### 2.2 Empirical Allometric Parametrisation

Although there are well-accepted parameterisations for vegetation in semi-arid ecosystems (e.g.(Rietkerk et al. 2002; Bonachela et al. 2015)), no comprehensive parameterisation exists for consumers. Thus, we used allometries to parameterise the consumer equations (Eqs (2)-(5b)), approach ecologically justified as trophic relationships and the energetics of foraging scale with consumer size (e.g. (Pawar, Dell, and Van M. Savage 2012; Kéfi et al. 2012)). Empirical allometries exist for most consumer parameters in our model (Table S.2) (Yodzis and Innes 1992; Pawar, Dell, and Van M. Savage 2012) except for the movement-related ones, i.e. dispersal coefficient, 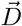, advection velocity 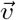, and maximum perception area Ω_*max*_.

For 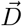 and 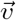, we used existing data on median spatial variance, advection length, and auto-correlation time-scales (Noonan et al. 2023) to obtain allometries: for 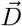, by fitting a linear curve to the data; for 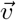, by using the *nls* function in R, following (Hirt et al. 2017). We derived an allometry for Ω_*max*_ by linking existing allometric relationships for visual acuity and axial diameters (Howland, Merola, and Basarab 2004; Veilleux and Kirk 2014; West, Brown, and Enquist 1999; Silva 1998), obtaining expressions for maximum perception radius, 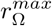 (in *m*). See SIA for more details.

### 2.3 Numerical simulations

To study desertification, we performed the analyses below for a wide range of aridity conditions, which in the SFV model was achieved by changing vegetation growth rate and, in the RV model, by changing rainfall levels.

As described in SIB, our simulations used two-dimensional grids with periodic boundary conditions. Our chosen grid spacing *dx* = 100*m* and timestep *dt* = 0.11*hr* captured movement scales and guaranteed computational convergence (Eq (B.2c)). We chose a grazer bodymass *M*_*G*_ = 20*kg* and predator bodymass *M*_*Pr*_ = 100*kg* that represent examples typical of semi-arid ecosystems (e.g. several types of gazelle for grazers, lions and hyenas for predators).

We initialised simulations either homogenously (constant density of all species at all grid sites) or heterogenously (vegetation clumps of 400 *m* spaced either 1, 2 or 3 *km* apart, removing 5%, 10% or 50% clumps randomly to represent different initial landscape degradation). To study the effects of consumer pressure, we tried four different combinations of scalings for the allometric grazer and predator attack rates (i.e. *a*_*V G*_ and *a*_*GP*_, respectively, taking values 1x, 25x the value in Table S.2). Unless otherwise noted, all results for the SFV model were obtained using homogenous initial conditions, as it is customary for this model. For the RV model, because a standard parameterisation leads to a high average vegetation density, we chose as baseline a 25x scaling of *a*_*V G*_ and 1*x* scaling of *a*_*GP*_, with vegetation clumps 2*km* initially apart, and with 10% initial clump removal. See SI B.1.1 for other alternative simulations.

### 2.4 Spatial Analyses

To analyse snapshots of the simulated system, we used either density thresholds or Gaussian Mixture Models (GMMs) (Biernacki, Celeux, and Govaert 2000) to classify pixels into high- or low-density sites. This step facilitated analysis of images by binarising them (i.e. mapping to black and white).

We then analysed separately binarised snapshots of vegetation, grazer, and predator, for each model, replicate and aridity level explored. To search for scale invariance in these snapshots, we used an algorithm to identify clusters (Hoshen and Kopelman 1976), then measured the probability distribution of cluster sizes and its Complementary Cumulative Distribution Function (CCDF). In the vegetation-only SFV model, these distributions behave as power-laws only at the desertification threshold, *a*_*c*_. To identify regular spatial patterns, we calculated the Fast Fourier Transform of binarised snapshots and their associated radially averaged power spectra; peaks in this metric indicate the presence of a characteristic spatial scale in the snapshot and, therefore, regularity (Kéfi et al. 2014; Chen, Halder, and Bonachela 2024).

To measure spatial similarity/correlation (i.e. synchrony) among snapshots of densities and/or consumer behaviour, we used three different metrics: maximum normalised 2D cross-correlation, *max*(*C*^Δ*T*^) (Zhao, Sang, and Duan 2019); Pearson 2D correlation coefficient; and adjusted mutual information (Vinh, Epps, and Bailey 2010). We constructed a synchrony timeseries by measuring these spatial similarity metrics (generically denoted here as *ℳ*^ΔT^) between a reference snapshot (ℳ _0_) and another snapshot separated in time by Δ*T*, and tried different values for Δ*T* (Fig. 3C-D). In addition to identifying e.g. oscillations in these timeseries, we also constructed the probability distribution of the ℳ^ΔT^ values observed over time and replicates (synchrony strength hereon), and explored whether distributions change with aridity.

**Figure 3:**
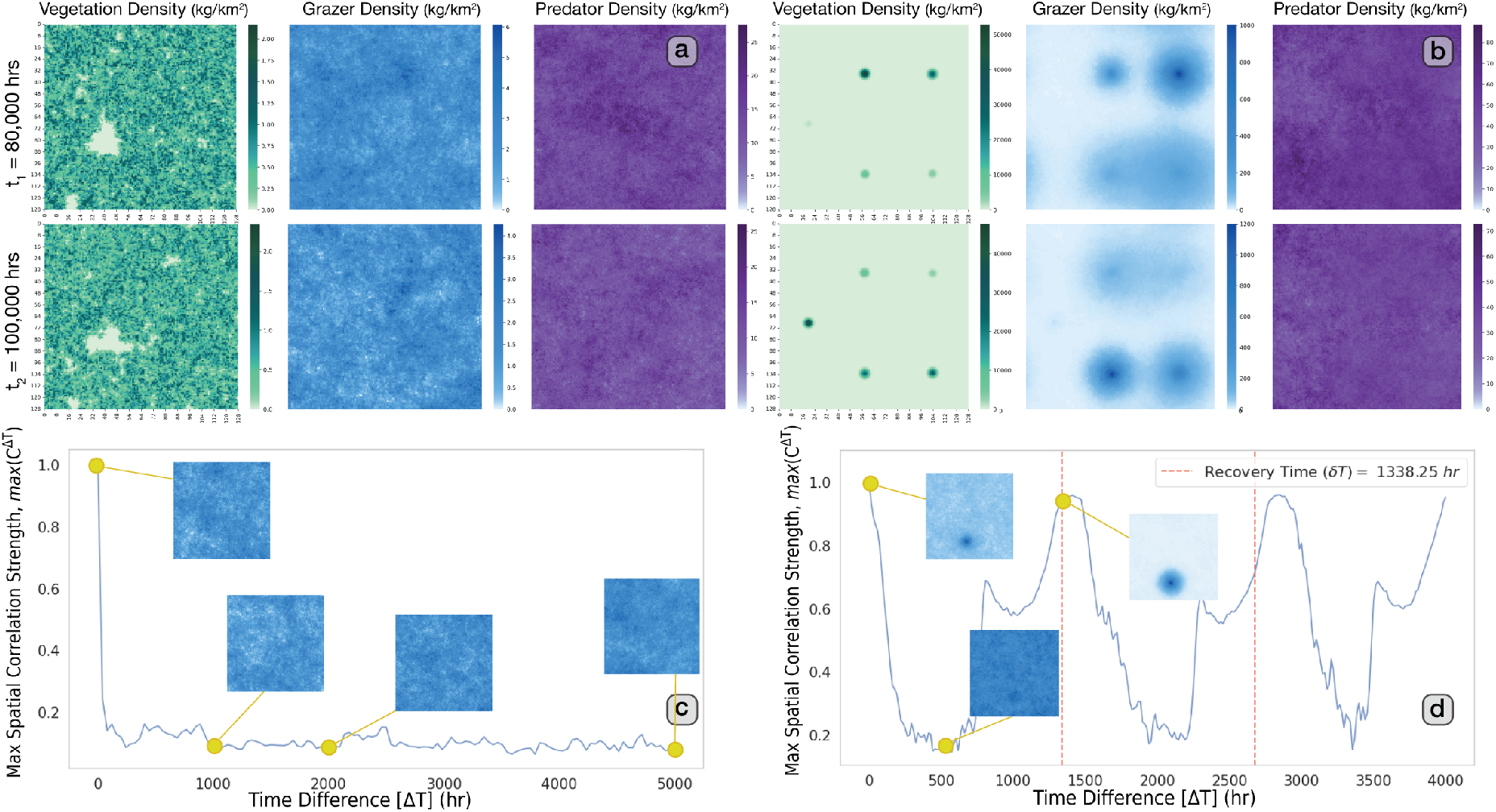
Spatio-temporal consequences of the interaction between structured vegetation and higher trophic dynamics. **a**-**b** Representative snapshots of vegetation, grazer, and predator biomass density (left, middle, and right panels, respectively, with darker colours representing higher density) at *t* = 8*·*10^4^ hr (top row) and *t* = 10^5^ hr (bottom row) for the scale-free case (panel a) and regular vegetation case (panel b). Snapshots correspond to *L* = 128*×* 128 systems (i.e. 12.8 *km×* 12.8 *km*), with vegetation parameters informed by Table S.1. Consumer parameters were informed by Table S.2, with grazer bodymass *M*_*G*_ = 20*kg*, and predator bodymass *M*_*Pr*_ = 100*kg*. In the regular case, we further used a 25x scaling of the allometric grazer attack rate (*a*_*V G*_) to induce boom-bust cycles (see SIB and Fig. S.11). **c**-**d** Spatial auto-correlations (measured as *max*(*C*^Δ*T*^), see MM) for grazers in scale-free (panel c) and regular vegetation settings (panel d). Grazers in the scale-free case quickly become asynchronous (correlation becomes negligible), while in the regular vegetation case they show sustained and strongly oscillatory temporal synchrony as spatial patterns show recurrence stemming from boom-bust cycles observed in the spatial vegetation pattern; the period of these correlation oscillations (in this case, Δ*T≈*1338 hr) represents the recovery timescale (*δt*) of the pattern.

See SIC and D for details on pattern and synchrony analyses, respectively.

## 3 Results

Without consumers, vegetation followed expectations (Fig. 1): the SFV model Eq.(1a) resulted in a smooth desertification transition as aridity increased (i.e. as growth rate, *a*, decreased) with scale-free vegetation patterns that emerged only at the transition point *a*_*c*_ (Scanlon et al. 2007); the regular vegetation model, Eqs.(1c)-(1d), resulted in an abrupt desertification transition preceded by a sequence of patterns (homogeneous vegetation, gaps, labyrinths, and clumps) as rainfall decreased (Rietkerk et al. 2002). SI B.1.1 summarizes results obtained with other simplified versions of the model.

### 3.1 Novel Consumer Movement and Perception Allometries

Following the steps detailed in Materials and Methods (MM) and SI 2.2, we obtained the following allometries for the non-directional dispersal coefficient 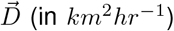 for grazers and predators, respectively (Fig. 2B):

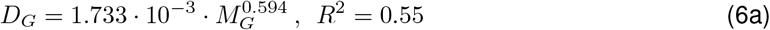

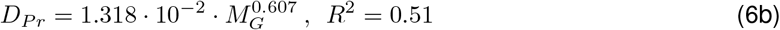

An allometry encompassing both consumers (*D*_*all*_, table S.2) did not provide a strong correlation (*R*^2^ = 0.28) because predators cover a larger area per unit time than grazers with identical bodymasses (black line in Fig. 2B).

For the advection velocity 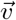 (in *m hr*^−1^, Fig. 2C), we obtained:

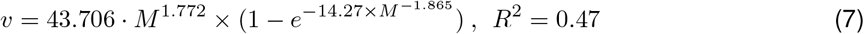

Finally, our allometry for the maximum perception radius 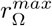 (in *m*) also required separated expressions, as predators detecting prey show substantially higher perception radii than similar-bodymass grazers detecting predators (Fig. 2D). The latter, in turn, is more limiting than detecting vegetation, and thus grazer perception is characterized only by 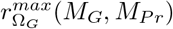 (SI 2.2):

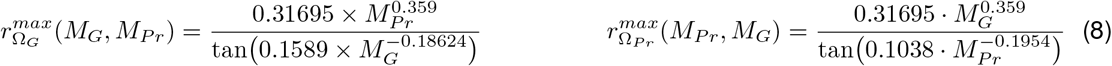

### 3.2 Grazer-only case

The addition of grazers (Eq.(2)) to either vegetation model, parameterising behaviour and movement with the allometries above, resulted in an eventual system collapse for any grazing rates *a*_*V G*_ capable of affecting vegetation density (SI B.1.1). This result thus highlights the need of including predators when a realistic spatio-temporal description of grazing is considered.

### 3.3 Bottom-Up Effects of Vegetation on Consumers

#### 3.3.1 Effects on Consumer Dynamics and Spatial Structure

The SFV model coupled with both grazers and predators (i.e. Eqs. (1a) and (2)-(3)) showed an emergent scale-free spatial organisation of vegetation and consumers (Fig. 3A). Although vegetation –like in the vegetation-only SFV model– only showed a power-law cluster size distribution at the desertification transition, the size distribution for consumer clusters showed a scale-free behaviour across a wide range of *a* values similar for both consumers (Fig. S.8A). The exponent of the power-law shown by grazers was similar to that of vegetation, but significantly different from the one from predators. These spatial patterns showed low synchrony, i.e. spatial auto-correlations quickly decayed over time, with snapshots a few hundred hours apart already showing negligible correlation (Fig. 3C).

Conversely, the RV model coupled with consumers (i.e. Eqs. (1b)-(1d) and (2)-(3)) showed a regular spatial organisation of vegetation and consumers. However, the emergent vegetation patterns differed with respect to those observed in the vegetation-only case, as the landscape was affected by grazer metapopulations that exploited vegetation patches before moving onto regenerated vegetation patches in dynamical ‘boom-bust’ cycles, and by predators always in pursuit (Fig. 3B, SI Video S.1). Despite the dynamic nature of these consumer spatial patterns, the radially averaged power spectra showed sharp peaks (represented as yellow bands in Fig. S.8B), indicating regular spatial organisation of these consumer metapopulations. Boom-bust cycles created strong, oscillatory synchrony (i.e. spatial auto- and cross-correlations for consumers showed large oscillations sustained over time, Fig. 3D) as a consequence of patterns that changed with time but were recurrently regenerated (see insets, and SI video S1). These recurrent patterns occurred across a range of rainfall levels wider than the range for which patterns occurred in the vegetation-only model. The period of these oscillations (hereafter referred to as ‘recovery time’) thus reflects the time period of the boom-bust cycles.

#### 3.3.2 Effects on Consumer Behaviour and Synchrony

Differences across vegetation models were further reflected in consumer emergent decision-making behaviour (given by the local dispersal probability, *γ*, see MM). Grazers in the SFV model showed a higher dispersal probability than in the RV model (blue distributions in Fig. 4A-B). Predators in the SFV model were also more prone to dispersion (higher *γ* value), but showed in the RV case a longer tail towards advection, particularly at intermediate rainfall levels (Fig. 4A-B, pink).

**Figure 4:**
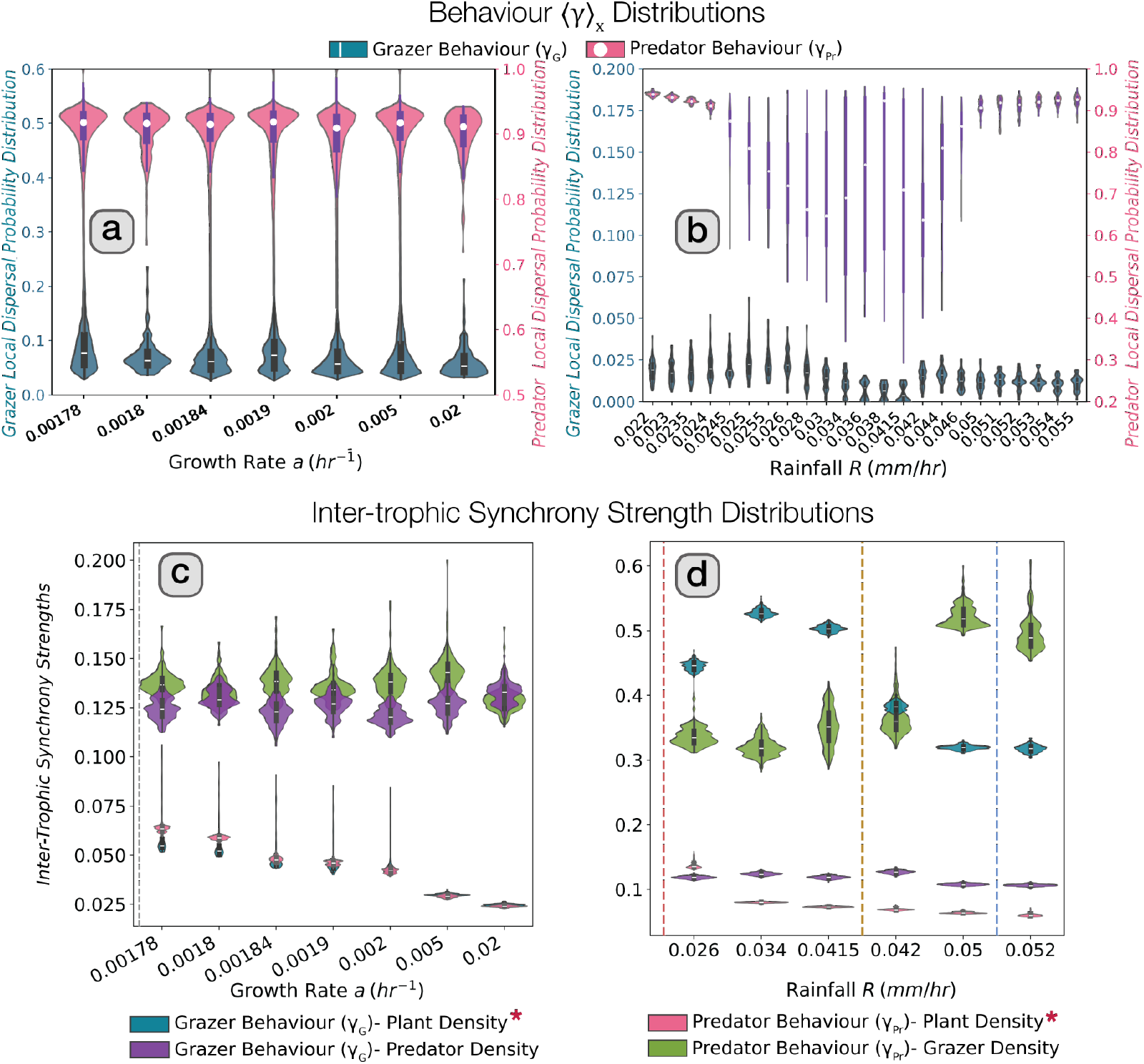
Effect of interactions between structured vegetation and higher trophic dynamics on behaviour and synchrony. **a**-**b** Spatially-averaged dispersal probabilities (⟨*γ*⟩ _*X*_) for grazers (in blue) and predators (in pink), for the scale-free case (panel a) and the regular vegetation case (panel b). Averages were estimated for individual frames corresponding to each surviving replicate (i.e. replicates where densities of all trophic levels are non-zero), measured between *t*∈ [1.5*·* 10^5^, 2 *·*10^5^] *hr*. In both cases, grazers mostly favour advection (low *γ*) whereas predators favour diffusive movement (high *γ*); in the regular vegetation scenario, however, the mean *γ* values are significantly lower than in the scale-free case (particularly for grazers) and predator ⟨*γ*⟩ _*X*_ distributions show long tails, indicating high behavioural variability. **c**-**d** Strength of inter-trophic synchrony (i.e. spatial correlations over time, *max*(*C*^Δ*T*^), between consumer behaviour and the density of another trophic level) in the scale-free case (panel c) and regular vegetation case (panel d). In both scenarios, inter-trophic synchrony between grazer behaviour and vegetation density (in blue), and predator behaviour and vegetation density (in pink) rise significantly with increasing aridity (Mann-Whitney U rank test, *p <* 0.001, denoted by (*)). In panel d, dashed vertical lines from left to right, mark the desertification threshold (red, ~ 0.024 *mm/hr*), the end of bistability (yellow, ~ 0.0419 *mm/hr*) and the transition to homogenous (i.e. non-patterned) vegetation (light blue, ~ 0.051 *mm/hr*) expected in the regular vegetation-only model.

In both SFV and RV cases, the probability distributions of auto-correlation strength (i.e. distributions of the spatial correlation between a metric and itself over time, e.g. built with the curves in Fig. 3C or D) for vegetation densities increased near the desertification threshold (Fig. S.2A-B), which is expected before transitions. Consumers in the SFV case showed low non-oscillatory auto- and cross-correlations across aridity levels, with small-yet-significant trends in the correlation between grazer behaviour *γ*_*G*_ and vegetation density *V*, and between predator behaviour *γ*_*Pr*_ and vegetation density *V* (Fig. 4C, S.5). These inter-trophic correlations rose as vegetation growth rates approached the desertification threshold (vertical line). The correlation of predator behaviour *γ*_*Pr*_ with grazer density *G* and viceversa (i.e. *γ*_*G*_ with *Pr*) were barely affected by growth conditions. In the RV case, inter-trophic correlations between grazer behaviour *γ*_*G*_ and vegetation density *V*, and between predator behaviour *γ*_*Pr*_ and vegetation density *V*, rose significantly with increasing aridity (i.e. decreasing rainfall). Similarly, cross-correlation between grazer behaviour *γ*_*G*_ and predator density *Pr* remained constant across growth conditions despite the decline in the correlation of predator behaviour *γ*_*Pr*_ and grazer *G* at lower rainfall levels.

We further examined whether features of the boom-bust cycle in the RV model relate to rainfall thresholds that are relevant in the vegetation-only case (e.g. distance to desertification). We observed that the recovery timescales of the boom-bust cycles (period of the oscillatory spatial auto-correlations, Fig. 5A-B) were *i)* generally shorter (i.e. faster recovery) for rainfalls for which, in the vegetation-only model, both the desertic and the vegetated states are alternative stable states (bistability region, see Fig 1D); *ii)* increased for rainfall levels for which bistability would not be expected but spatial patterns would be predicted (mid rainfall range); and *iii)* plunged for rainfall values for which homogeneous vegetation would be expected (high rainfall). In other words, increasing rainfall beyond the bistability threshold (brown dashed line) led to a sharp slow-down in recovery and longer cycles (Fig. S.9), whereas crossing the rainfall threshold for pattern formation (blue dashed line) led to a speed-up of recovery and shorter cycles. We observed similar trends in the periods of the oscillations corresponding to correlations across trophic levels: between predator behaviour *γ*_*Pr*_ and grazer density *G*, and grazer behaviour *γ*_*G*_ and vegetation density *V* (Fig. 5C); and between predator and grazer density, and grazer and vegetation density (*Pr*-*G* and *G*-*V*, Fig. 5D).

**Figure 5:**
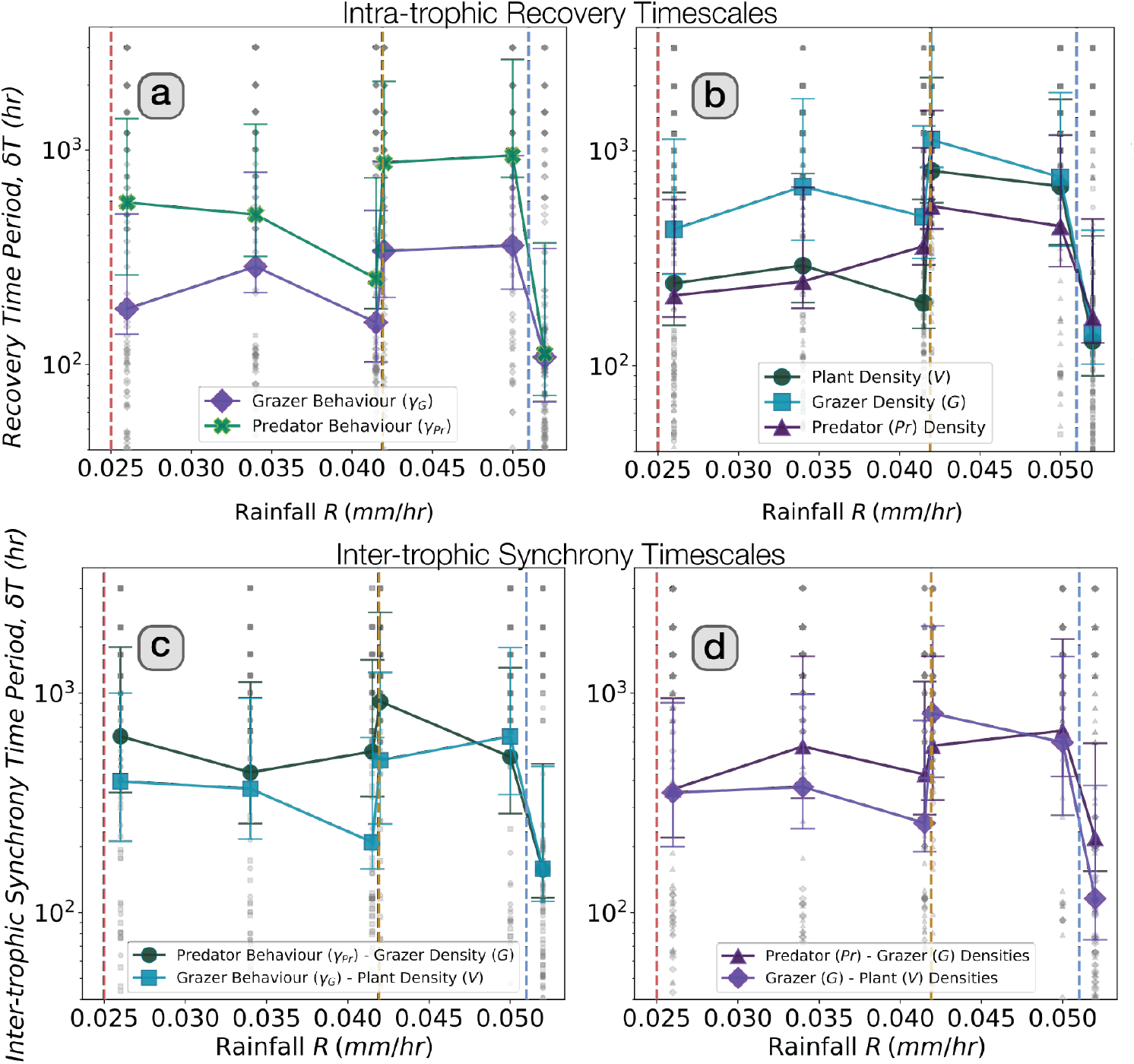
Effect of interactions between structured vegetation and higher trophic dynamics on behaviour and synchrony timescales in the regular model (RV). Dashed vertical lines from left to right indicate the desertification threshold (*red*, ≈0.024 *mm/hr*), the end of bistability (*yellow*, ≈0.0419 *mm/hr*) and the transition to homogenous (i.e. non-patterned) vegetation (*light blue*, ≈0.051 *mm/hr*) in the regular vegetation-only model. **a**-**b** Time period of the spatial intra-trophic auto-correlations for consumer behaviour and biomass densities respectively (equivalent to the recovery timescale, *δt*, of spatio-temporal boom-bust cycles). **c**-**d** Time period of the inter-trophic correlation (i.e. cross-correlation) between consumer behaviour and resource density, and between consumer and resource densities respectively, for the regular vegetation case (equivalent to tracking timescales).

The synchrony trends above were largely but not fully robust against other metrics measuring spatial similarity (Adjusted Mutual Information, 2D Pearson Correlation, Fig. S.3-S.4), and under increased predatory pressure and habitat degradation (Fig. S.6).

### 3.4 Top-Down Effects on Vegetation

#### 3.4.1 Effects on Vegetation Patterns and Desertification Transitions

For the SFV model, while consumer dynamics did not prevent the emergence of scale-free patterns, they affected the desertification threshold separating barren and vegetated states. At this critical threshold *a*_*c*_, the probability distribution of vegetation cluster sizes follows a power law (Muñoz et al. 1999; Kéfi et al. 2007). For the vegetation-only model, the threshold was *a*_*c*_ ≈1.694*·* 10^−3^ *hr*^−1^ whereas, in the presence of consumers, it shifted to *a*_*c*_ ≈1.728*·* 10^−3^ *hr*^−1^ (Fig. 6A-B).

**Figure 6:**
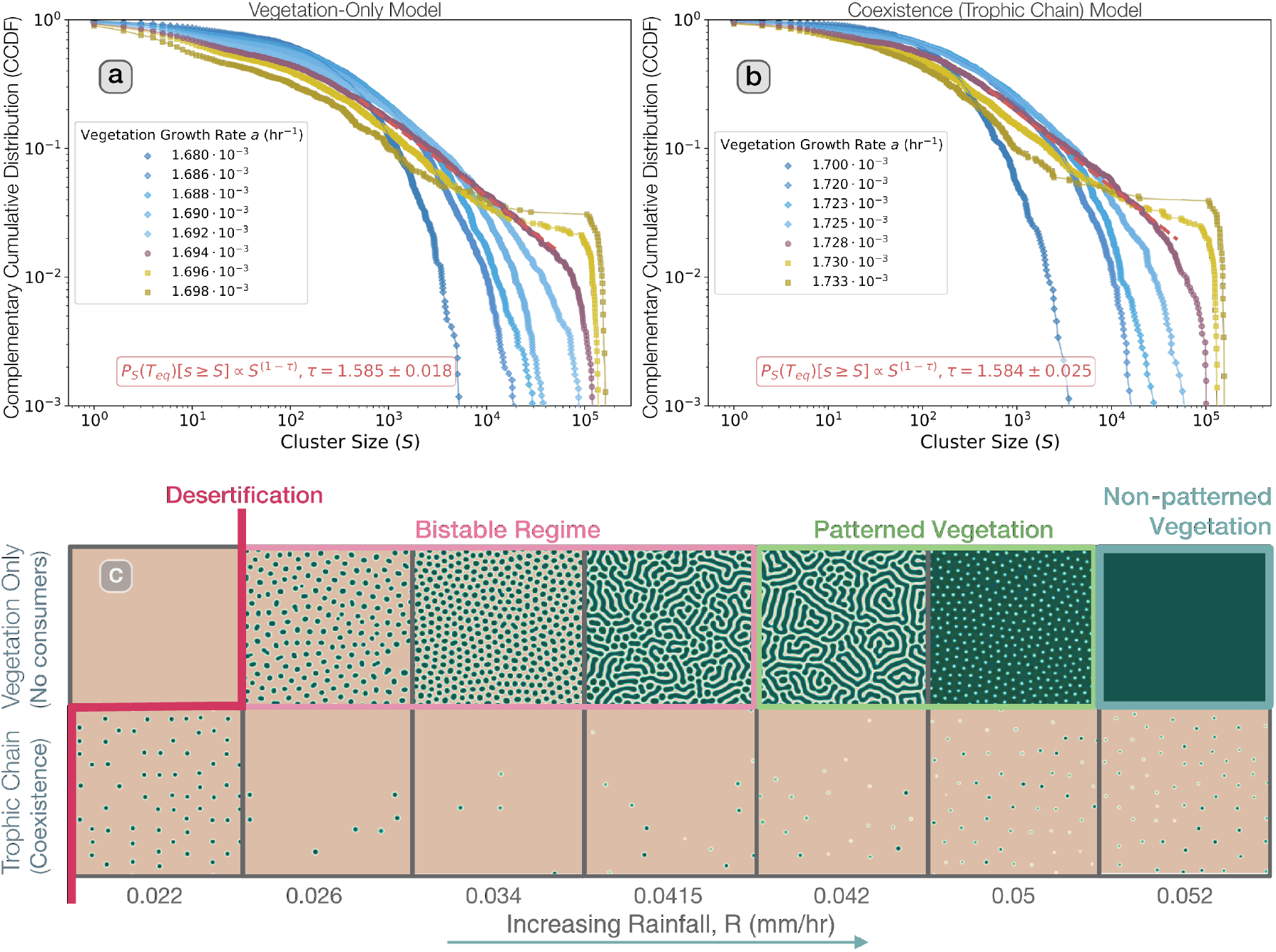
Effect of consumer spatial dynamics on desertification transition and emergent vegetation patterns. **a**-**b** Complementary cumulative probability distributions (CCDF) for vegetation cluster sizes (*P*[*s* ≥ *S*]) in the vegetation-only SFV model (panel a), and the full SFV model (panel b), for different values of vegetation growth rate. The presence of consumers increase the desertification threshold from the critical growth rate, *a*_*c*_ 1.694 *·*10^−3^ *hr*^−1^ to *a*_*c*_ ≈1.728 *·*10^−3^ *hr*^−1^, but does not affect the associated exponent *τ* ≈1.585. The distributions were obtained in a *L* = 512 *×*512 pixel (i.e. 51.2 *km ×*51.2 *km*) system at stationarity (*t* = 120226 *hr*). **c** Spatial patterns of vegetation emerging for different rainfall levels in the vegetation-only RV case (upper row), and the counterpart with consumers (lower row). In the absence of consumers, a barren state transitions into vegetation patterns that follow the classic clump-labyrinth-gap pattern progression as rainfall increases, with patterns disappearing at very high rainfall levels. The presence of consumers leads to dynamic regular clump-like vegetation patches only, with an increased range of rainfall levels for which patterns emerge: clumps first emerge from desert at lower rainfall levels for the complete model than they do for the vegetation-only model (red line), and homogeneous vegetation is observed at higher rainfall levels (bold blue box). Snapshots were taken in an *L* = 256 *×*256 system (25.6 *km×* 25.6 *km*), at the same equilibrium time as in the SFV model, i.e. *t* = 120226 *hr*. Vegetation was initialized in clumps 1*km* apart, removing 50% of the initial setup.

For the RV model, the presence of consumers altered both the emergence and shape of vegetation patterns (Fig. 6C). Consumer dynamics altered the ‘clumps-labyrinths-gaps’ progression expected in the vegetation-only system as rainfall increases; instead, the dynamic clumps of the boom-bust cycles emerged for all rainfall levels for which vegetation survived (Fig. 6C). Moreover, the presence of consumers lowered the desertification threshold from ~0.024 *mmhr*^−1^ to ~0.021 *mmhr*^−1^. Unsurprisingly (given the difference in patterns), the peaks of the radially-averaged power spectra in the vegetation-only case occurred at different locations (i.e. different pattern characteristic scales) and changed with rainfall in a different way than in the complete trophic-chain model (Fig. S.10).

## 4 Discussion

Our framework links dryland vegetation dynamics and resilience to consumer-resource dynamics, consumer foraging behaviour, and synchrony within and across trophic levels. Existing work has focused on the interaction of regular vegetation patterns and the dynamics of grazers, partially studying the feedback loop between pattern formation, landscape resilience, and consumer dynamics (Schneider and Kéfi 2016; Siero et al. 2019; Ge and Liu 2022; Maimaiti, Yang, and Wu 2022; Singha, Uecker, and Meron 2025). We provide here a more complete picture that includes predators and a more ecologically realistic implementation of behaviour and movement, and that compares regular and scale-free vegetation. The ecosystem collapse and instability we observed in the absence of predators emphasises the need for models that incorporate not only grazers, but also predators and realistic behaviour. More generally, this result would support the importance of predators in dryland conservation by promoting the resilience of semi-arid ecosystems (Morris and Letnic 2017; Farías et al. 2021).

To include behaviour, we developed novel empirical allometries for the (non-directional) consumer dispersal coefficient 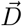 the (directional) consumer advection velocity 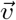, and the maximum perception area Ω_*max*_. Our deduced 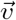 is in moderate agreement with existing maximum speed models (Hirt et al. 2017), which are typically used to inform 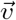 (Hirt et al. 2018). This agreement indicates that a mammal trade-off between aerobic metabolism and anaerobic capacity that applies when moving at maximum velocity also applies when moving at their foraging advection velocities. Additionally, our derivation of Ω_*max*_ shows that mammalian vision is diffraction-limited rather than turbulence-limited (Born and Wolf 2019) for the vast majority of realistic predator-prey pairs (SIA), and thus visual resolution for the consumer is limited by the perceiving eye lens rather than atmospheric effects.

In the RV model, and despite the demographic stochasticity of vegetation and consumers, the interaction of vegetation SDFs and consumer trophic behaviour led to regular vegetation landscapes that are dominated by regularly organised travelling waves of grazers, which is in line with recent theoretical work that used a simpler model with grazing but not predation (Maimaiti, Yang, and Wu 2022; Singha, Uecker, and Meron 2025). Similar to (Maimaiti, Yang, and Wu 2022), increasing grazer pressure (i.e. using higher *a*_*V G*_) shifts the dynamics from spatially homogenous coexistence to boom-bust cycles. In our framework, however, these cycles emerge as grazers interact with water-vegetation dynamics to form roving metapopulations that shape vegetation into distinct islands and consume them to near-extinction; predators serve as a keystone species, enabling coexistence by limiting grazer growth and influencing their movement, thus contributing to vegetation recovery. In these boom-bust cycles, grazers concentrate away from predator detection, prompting predator advection in a perpetual bid to catch up with their prey; thus, predator perception range influences grazer spatial spread, which in turn limits vegetation cluster size and pattern wavelength. Boom-bust cycles are particularly marked for mid-low rainfall, where vegetation recovery times are low, and are reminiscent of empirically observed dynamics in dryland communities such as boom-bust cycles in rodent populations in the Australian outback (Pavey et al. 2017). This cycle of vegetation near-depletion and regeneration creates robust spatio-temporal trophic patterns that show strong synchrony (i.e. large correlation). The oscillatory character of this trophic synchrony is indicative of travelling waves across trophic levels (Bjørnstad and Bascompte 2001) which, in our case, are the boom-bust cycles. These dynamic spatial patterns thus differ from the classic patterns predicted with vegetation-only models, in which vegetation ultimately reaches a stationary pattern that depends on rainfall levels (Rietkerk et al. 2002).

Consumers in the SFV case, on the other hand, show scale-free probability distributions of cluster sizes, with similar exponents for vegetation and grazers but significantly different exponent for predators. Our results, however, show that grazer fractality does not result from that of vegetation (SI B.1.1), and thus we hypothesise that for both vegetation and grazers the spatial distribution results from the dominance of demographic stochasticity, in the case of grazers also influenced by predation. Behaviour is key for trophic dynamics and spatial patterns: trophic cascades observed in the SFV model disappear when eliminating behavioural modulation (SI section B.1.1), which is consistent with work that observed that the “ecology of fear” can generate trophic cascades (Schmitz, Beckerman, and O’Brien 1997). The above hypotheses are validated by the increased grazer dispersal with respect to the RV case, in which grazers need advection to reach the dynamic vegetation clusters. In turn, and as a result of the scale-free grazer distribution, predators do not need to advect to find prey, and thus their associated dispersal behaviour prevents the emergence of any characteristic scale, that is, predators also show a scale-free distribution. While the vegetation cluster size distribution shows, with or without consumers, scale-free behaviour only at the desertification threshold *a*_*c*_, these consumer scale-free distributions emerge for a range of vegetation growth rates below and above the desertification threshold (i.e. generalised scale invariance). We hypothesise that this result is a consequence of trophic dynamics and consumer behaviour, potentially because grazers selectively target the higher density vegetation clusters that emerge above the critical point, and avoid the more depleted clusters emerging below the critical point, with predators precluding grazers from depleting vegetation.

Our results also show that bottom-up effects of desertification strongly influence the ‘geography of spatial synchrony’ (Walter et al. 2017). Both scale-free and regular cases show strong Moran effects, that is, a dependence of biotic synchrony with environmental drivers, in this case aridity (Liebhold, Koenig, and Bjørnstad 2004). Both grazer and predator densities and behaviour show higher synchrony with vegetation density at higher aridity (i.e. lower vegetation growth/rainfall levels). This result agrees with the match-mismatch hypothesis, which postulates that inter-trophic synchrony positively affects consumer fitness because fitness depends on synchronisation with food resources (Visser and Gienapp 2019; Rao et al. 2024). We hypothesise that increased correlation between grazer behaviour *γ*_*G*_ and vegetation density *V* under increasing stress translates into an increase correlation between predator behaviour *γ*_*Pr*_ and vegetation *V*. Increased environmental stress did not however translate into increased grazer-predator inter-trophic synchrony (i.e. correlation of predator behaviour *γ*_*Pr*_ with grazer density *G*, and grazer behaviour *γ*_*G*_ with predator density *Pr*). This lack of synchrony indicates that grazers in both scale-free and regular cases respond primarily to bottom-up pressures with increasing aridity. We speculate that grazer-predator inter-trophic synchrony is negated by grazer behavioural modulation (i.e presence of boom-bust cycles) in the RV case, and by the generalised scale invariance shown by consumers across aridity levels in the SFV case.

The jumps in synchronisation timescales observed at rainfall thresholds that are characteristic of the vegetation-only model indicate that the oscillatory patterns of the full RV model remain somewhat tied to the underlying vegetation-only transitions. We hypothesise that the slowing down of recovery and interaction timescales above the desertification and bistability thresholds relate to critical-slowing-down effects that emerge in the underlying vegetation–only dynamics.

We propose that these changes in synchrony strengths (for SFV/RV landscapes) and timescales (for RV landscapes) are Early Warning Signals (EWS) of desertification. For example, stakeholders could sample at given time-intervals the spatial density/abundance of each consumer population (using remote-sensing and/or drone footage (Marescot et al. 2025)) over a window of time that corresponds to the expected recovery or interaction time (in our simulations, a few months). Once abundance is mapped onto geospatial data, the spatio-temporal metrics measured here can be calculated. Observing periodicity in correlations/synchrony would enable the definition of a recovery and interaction timescales, for which a hump shape as aridity increases (similar to the grazer or plant curves in Fig. 5) would indicate decreasing ecosystem health and resilience, and impending desertification. Similarly for increasing inter-trophic synchrony of consumers (especially grazers) with vegetation density. Our EWSs thus add to traditional ones (also observed here) such as increases in vegetation density auto-correlation strengths close to desertification (for both the scale—free and regular cases) (Dai, Korolev, and Gore 2013; Kéfi et al. 2014). In contrast with these existing spatial EWSs, which monitor vegetation-only metrics, our synchrony-based EWSs also encode information about trophic dynamics and how they affect vegetation.

Finally, similarly to previous studies (Schneider and Kéfi 2016; Ge and Liu 2022; Siero et al. 2019), we see that spatially heterogenous top-down pressures interact with the pattern-formation mechanisms. In our case, regular vegetation is shaped by grazers to show clump-like patterns across a wide range of rainfall levels, in contrast to the diverse patterns observed in the vegetation-only case. Thus, in line with recent empirical evidence (Pichon et al. 2025), consumers in the RV case increase the circularity of patches and the connectivity of bare soil. Crucially, consumers increase the resilience of vegetation to desertification, as they allow vegetation to persist in more arid (lower rainfall) environments, and the only vegetation pattern that emerges (clumps) persists for rainfall levels that are higher than those for which the vegetation-only model predicts homogeneous vegetation. Our findings thus add nuance to recent observations that grazer dynamics positively impact the resilience of regular dryland landscapes (Bonachela et al. 2015; Tarnita et al. 2017; Pichon et al. 2025; Singha, Uecker, and Meron 2025). In contrast, the desertification threshold for the SFV case shows a shift towards lower aridity (higher growth rate) while conserving the cluster-size distribution (similar to that for the Kalahari landscape in Fig. 1a), indicating that consumers make scale-free landscapes more vulnerable to desertification without changing their spatial distribution.

Our findings, however, have important limitations. For example, we did not consider other important drivers of desertification such as land use (Reynolds et al. 2007b; Intergovernmental Panel on Climate Change 2022b) or rising temperatures (Intergovernmental Panel on Climate Change 2022a). In addition, we considered only tri-trophic communities instead of more realistic trophic networks. Finally, our focal spatial scale (*dx* = 100*m*) is 5-10 times larger than the scale of empirical vegetation patterns observed arising from SDFs (Rietkerk and van de Koppel 2008), which is a consequence of the separation between the spatial scale for these patterns and that of consumer behaviour, and of our focus on the latter as a key novelty in consumer description. Although large-scale SDF-like vegetation patterns exist (Fig. S.12), this spatial separation in itself calls for caution when attempting to use our framework for specific real-world systems. Nonetheless, our results more generally highlight the intricate interplay of bottom-up and top-down pressures in shaping dryland communities, the role of inter-trophic synchrony as a buffer to aridity, and the need to consider trophic dynamics and synchrony when analysing dryland resilience and desertification transitions.

## Acknowledgements

The authors would like to thank Sonia Kéfi, members of the Bonachela lab, Ricardo Martínez García and Miguel A. Muñoz for helpful discussions. This research was partially funded by NSF grant DMS-2052616 to J.A.B.

## Supplementary Information

### A Derivation of Allometries for Max Visual Perception Radius

We define the maximum perception radius 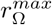 as the maximum focal distance at which an organism can reliably visually detect and respond to a resource or target. This radius depends on both physical detection limits (influenced by the size of the eye-lens for visual perception and atmospheric effects) as well as complex ecological and phylogenetic factors, such as phyla, diet (herbivory vs carnivory), and diurnal or nocturnal activity patterns (Veilleux and Kirk 2014).

Since we focus on visual perception, we assume that perception is more likely to be visually based (and limited) in diurnal mammals, and thus focus subsequently on diurnal taxa.

### A.1 Existing Allometries

We started by noting existing allometries that link the axial diameter (*AD*, in *mm*) of vertebrate eyes to their body-masses (*M*, in *kg*) (Howland, Merola, and Basarab 2004):

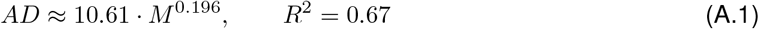

and another allometry that links the length of mammals (*L*_*A*_ in *m*) to their body-masses (Silva 1998):

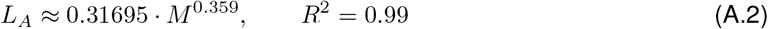

Next by analysing Table S.1 from ref. (Veilleux and Kirk 2014) (see SI Fig. S.1B), we arrived at the following relationships linking behavioural visual acuity (*θ*^*′*^, in cycles-per-degree *cpd*) for diurnal herbivores and carnivores as follows:

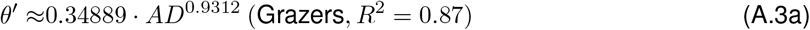

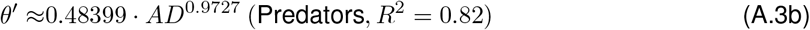

Then we converted this behavioural visual acuity to behavioural visual resolution (*θ* in °) with *θ* = 1*/*(2*θ*^*′*^) and, utilising (A.1), we arrived at:

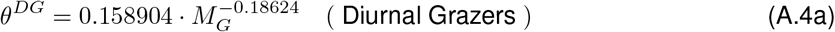

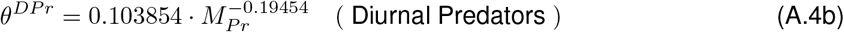

Following the same steps but considering all data together, we further derived a common visual resolution across all diurnal mammals as follows:

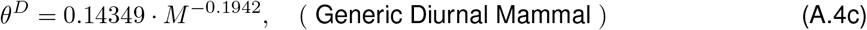

Note that the allometric scaling exponent for *θ*^*DP r*^ is much smaller than *θ*^*DG*^, indicating that predators have better (i.e. smaller) visual resolution and thus higher perception radii than equivalently sized grazers for a wide range of bodymasses (Fig. 2D). This result makes evolutionary sense in many predator-prey interactions, where predators would presumably have higher fitness benefits in detecting prey at a longer perception threshold with respect to the perception threshold of the prey for the predators.

### A.2 Atmospheric Effects and the Diffraction Limit

Eqs. (A.4a)-(A.4c) provide relationships for visual resolution as a function of the numerical aperture of the eye-lens. Nonetheless, resolving power is still subject to atmospheric effects. With increasing perception distance, atmospheric distortions grow larger and the numerical aperture of the lens no longer effectively determines the effective resolution. We refer to this scenario as ‘turbulence-limited’ perception, where local turbulence effects (influenced by temperature gradients, wind speeds, diurnal heating, etc.) dominate the determination of visual resolution. This limit is in contrast to ‘diffraction-limited’ perception, where the numerical aperture of the lens primarily determine visual resolution (Born and Wolf 2019; Yoder and Vukobratovich 2011). We define the perception radius separating diffraction-limited visual acuity from turbulence-limited acuity as 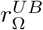(in *m*). In what follows, we examine the diffraction-limited case, and provide an estimate for 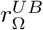.

In the diffraction limit, i.e. when the maximum perception radius 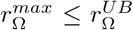, we can simply relate visual resolution *θ* to the angle subtended by the length of the perceived object (i.e. predator for grazers, and grazer for predators) at the maximum distance such that it is still visible – i.e. 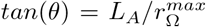 (see Fig. S.1B). Thus by subsituting Eqs. (A.4a)-(A.4b), we arrive at Eqs. (8) in the main text, which we reproduce here:

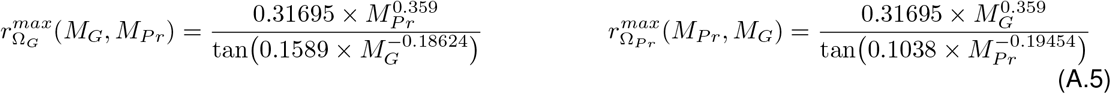

A corresponding allometric scaling relationship for the perception radius of grazers for vegetation can be reached by noting that the length of vascular vegetation (*L*_*V*_ in *m*) scales with plant mass as (West, Brown, and Enquist 1999) *L*_*V*_ = *l*_0_*/*(1− *n*^−1*/*3^) ∝ *M* ^0.25^*/*(1− *n*^−1*/*3^), where *l*_0_ is the length of the trunk (which has the allometric form: *l*_0_ = *c*_*l*_ *M* ^0.25^) and *n* is the branching ratio (i.e. the number of daughter branches borne out of any parent branch). This expression allows us to follow the steps above to reach the following form for the perception threshold of grazers for vegetation, 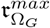:

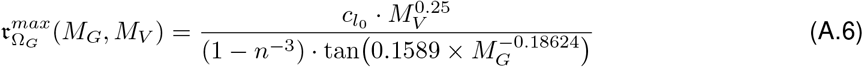

Due to the variability of 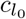 and *n* across plants, however, this allometry cannot be used to generically estimate vegetation length *L*_*V*_ (West, Brown, and Enquist 1999) and thus, by extension, 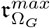. Nonetheless, across a range of potential 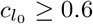 (i.e. a vascular plant of mass 1*kg* would have a trunk length of 60 *cm* or more), and *n* ≥ 2 (i.e. branching ratios of 2 or more), we found 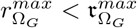 for any given *M*_*G*_ and where *M*_*Pr*_≤ *M*_*V*_ (assuming reasonable terrestrial predator bodymasses up to ≈400 *kg*). This establishes 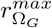 (Eq. (8)) as the most limiting maximum perception radius for grazers for a broad range of parameters and thus allows us to bypass the unreliability of *L*_*V*_.

Regarding the upper bound for diffraction-limited vision 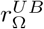, the ratio *D*_*EP*_ */r*_0_ determines the effect of atmospheric turbulence on vision, where *D*_*EP*_ is the pupil entrance length and *r*_0_ is the Fried parameter (the coherence length of the atmosphere, where the air is well-mixed and non-turbulent) (Fried 1966). In particular, when *D*_*EP*_ */r*_0_ *<* 3.7, atmospheric turbulence primarily produces image motion or shifting, whereas *D*_*EP*_ */r*_0_ ≥3.7 leads to image blur and visual resolution becomes independent of numerical eye-lens aperture (Yoder and Vukobratovich 2011).

When imaging along a horizontal path of length *r*_Ω_, the Fried parameter *r*_0_ varies with distance and is combined with another atmospheric turbulence parameter, the index of refraction structure parameter or atmospheric turbulence strength 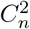 (expressed in units of *m*^−2*/*3^) (Hardy 1998) through the relationship 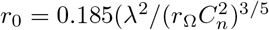. By assuming typical values of 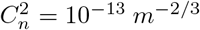 (medium turbulence effects common around mid-day) (Yoder and Vukobratovich 2011), assuming *λ* = 550 *nm* (green light), and further assuming *D*_*EP*_ = (1*/*6) *×AD* (which is the proportionality factor for human eyes), we can arrive at the following heuristic estimate for 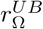:

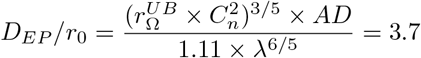

Thus, using Eq. (A.1) and simplifying:

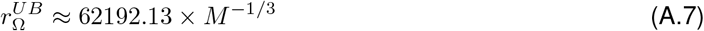

Larger animals have larger eye lens, which in turn are more likely to be constrained by the Fried parameter, and thus have smaller perception thresholds beyond which they become turbulence-limited. Nevertheless, our results show that both grazers and predators are almost always in the diffraction limit, even for very large grazer and predator sizes (Fig. 2D).

## B Numerical Integration of the Model Equations

Integrating stochastic partial differential equations (SDEs) with demographic noise can be challenging as, when the main stochastic variable is close to zero, stochasticity may yield negative values, which are not only biologically unreasonable in our case but also mathematically problematic due to the square root noise. An elegant solution to the problem is to use an existing split-step integration scheme that prevents negative values for the variable (Dornic, Chaté, and Muñoz 2005), which we adapted to our specific model.

At each time step we first calculated, at each local site (*i, j*), the advection velocity direction 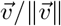 and the local dispersal probability 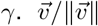 was sampled as a random 2*D* unit vector, based on our assumption that consumers choose their advection direction at random as consumers cannot perceive beyond their perception radius in factoring their advective relocation behaviour, and by extension the optimal direction for relocation. Further note that *γ* integrands (Eqs. (5a)-(5b)) were evaluated as their equivalent convolution kernels:

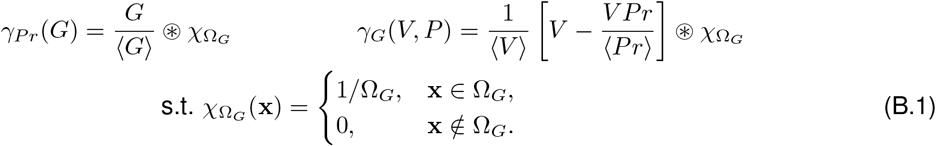

where ⊛ represents 2*D* convolution. This reduced the 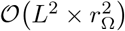 integrand evaluations in Eqs. (5a)-(5b) to quicker *O(L*^2^ *×* log *r*_Ω_) evaluations. We further imposed *γ* ∈ [0, 1] bounds after all such evaluations, to avoid unrealistic values for a probability.

Next, at each site, we split our SDEs into linear and non-linear terms, grouping the linear terms together with the noise to compose linear Langevin equations of the form 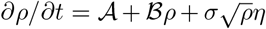, with *ρ* representing the stochastic variable of the focal SDE (e.g. *G* in Eq. (2)) and *A* ≥0. For our consumer equations, using a standard Upwind Finite Difference Method (UFDM) scheme to discretise the spatial derivatives (LeVeque 2002) leads to the following linear Langevin equation (see details in section B.2 below):

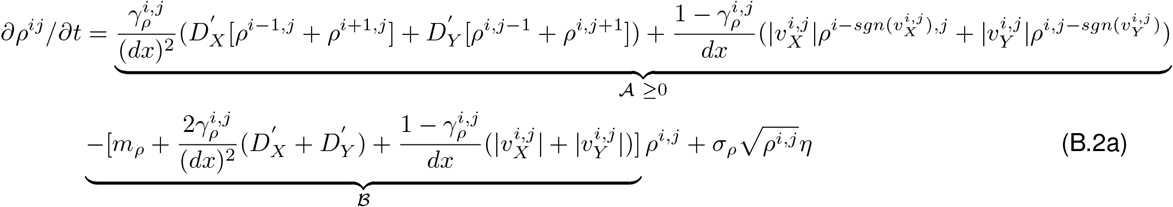

where *sgn*(*·*) is the sign function (1 if the argument is non-negative, −1 otherwise) and *dx* is the grid spacing. Note that, as the UFDM and other finite-difference methods introduce excess (extranumerary) diffusion (E. Ewing and Wang 2001), we corrected for this excess using the corrected diffusion coefficient 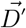 (see details in section B.3 below):

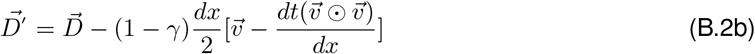

where ⊙ denotes the Hadamard product and *dt* is the time step. Note that the UFDM scheme has the following Courant–Friedrichs–Lewy condition for ensuring convergence (LeVeque 2002), which constrains *dt* for a given *dx*:

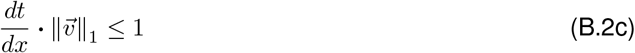

where 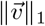 represents the *l*^1^ norm of 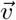.

For the vegetation population *V*, which always spreads by local dispersal (i.e. through cloning or seed dispersal), *γ*_*V*_ = 1 necessarily. Thus, for the scale-free case, the linear Langevin Eq. (B.2a) (i.e. Eq. (1a) without the quadratic term) reduces to:

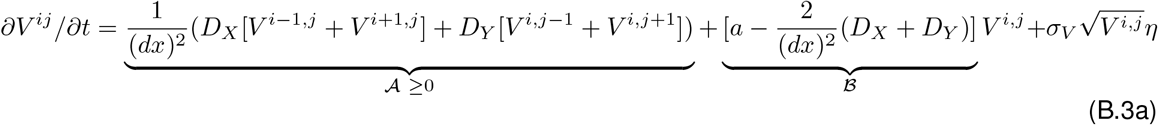

and for the regular case (Eq. (1d) without the first term):

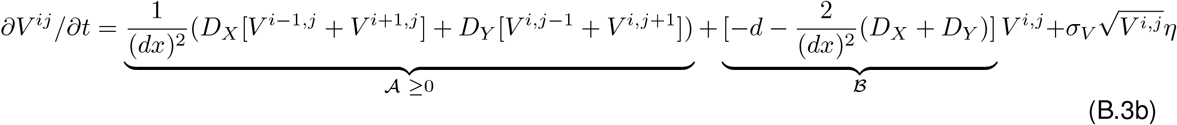

Re-writing Eqs.(B.2a), (B.3a)-(B.3b) in the simplified form 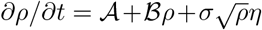 allowed us to write and solve its equivalent Fokker-Planck equations (FPE), which has the following probability distribution as a known analytical solution (Dornic, Chaté, and Muñoz 2005):

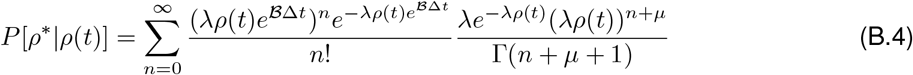

where *λ* = 2 ℬ*/*(*σ*^2^(*e*^*ℬt*^ −1)) and *µ* = −1 + 2 *A /σ*^2^. Sampling this probability distribution at each site provided the intermediary solution *ρ*^*^, which was then used as the input to integrate the remaining (i.e. non-linear) terms of the original equations. We integrated numerically these non-linear terms using a standard Runge-Kutta 4 method.

### B.1 Numerical Simulations and Sensitivity Analysis

We implemented the above steps in C++23 using a 2*D* system with grid sizes ranging from 128*×* 128 to 512*×* 512, with periodic boundary conditions. The estimation of *γ* via the 2*D* convolutions (see Eq. (B.1)) was implemented using FFTW3 (Frigo and Johnson 2005). We chose grid spacing *dx* = 100*m* and time step *dt* = 0.11*hr* to guarantee convergence (see Eq (B.2c)). For these simulations, and unless otherwise noted, we chose a grazer bodymass *M*_*G*_ = 20*kg* and predator bodymass *M*_*Pr*_ = 100*kg* that represent typical grazers and predators found in semi-arid ecosystems (e.g. several types of gazelle for grazers, lions and hyenas for predators). We ran simulations until a maximum time *t* = 2 *·*10^5^ hr or until vegetation extinction, whichever occurred first.

We initialised simulations either homogenously (that is, with a constant density of all species at all grid sites) or heterogenously (vegetation clumps of 400 *m* spaced either 1, 2 or 3 *km* apart, with further 5%, 10% or 50% of these clumps randomly removed, indicating increased initial landscape degradation). This led to a total of 9 distinct, heterogenous initialisations, in addition to the homogenous initialisation. For each of these initialisations, we explored four distinct scalings consumer attack rates (i.e. *a*_*V G*_ and *a*_*GP r*_ respectively, taking value 1x, 25x the value in Table S.2), alongside various aridity levels (growth rates for SFV model, rainfall levels for RV model) leading to vegetated and desertic states of the ecosystem, thus capturing the desertification transition. In most cases involving elevated grazer attack rates (25x), we saw the emergence of boom-bust cycles in the RV model, this is summarised across 16 distinct initial conditions in Figure S.7. We further measured the synchrony characteristics of these boom-bust cycles across 4 distinct initialisations, following the analysis protocol in SID. This sensitivity analysis is presented in Figure S.6.

All simulation and analysis scripts, written in C++ and Python respectively with optional CUDA acceleration, are available on Github.

#### 5.1.1 Sensitivity analysis

To understand better how the different components of the model influence our results, we ran a series of simulations in which we removed some of those components, or assumed different values for key parameters.

##### Without consumers

As explained in the main text, when consumers are not present our models revert to classic vegetation models (Fig. 1c and d). The SVF model leads to a smooth desertification transition that occurs at a particular aridity level, characterised by a vegetation growth rate *a* = *a*_*c*_; at exactly that growth rate, the spatial distribution of vegetation shows scale-free properties, which for example translate into a power-law behaviour for the distribution of vegetation cluster size. The RV model, in turn, shows a sequence of different vegetation spatial patterns as aridity increases, in this case represented by a decrease in rainfall levels; thus, as rainfall decreases, homogeneous vegetation self-organizes into gaps, then labyrinths, then clumps and, ultimately, desertification occurs and all vegetation disappears. Before the latter occurs, however, the system shows bistability (i.e. both vegetated and desertic states are stable for the same rainfall level).

##### Without predators

When predators are not present, for both models grazers grow too fast and drive vegetation to extinction, in turn causing their own demise (Fig. S.13). This result confirms that predators play a key role in our tri-trophic ecosystem by exerting an essential top-down regulation on grazers.

##### Without vegetation patterns

When forcing the condition of a homogeneous vegetation distribution (i.e. ∂*V/*∂*t* = 0 and *V* = 1 *kg/km*^2^ across the whole system), consumers in both the SFV and the RV models showed a spatial distribution with scale-free properties (see cluster size distribution in Fig. S.14). This result indicates that the scale-invariant distribution for consumers observed in the SFV model is not imposed by the underlying vegetation pattern but rather emerges from the consumer dynamics. Moreover, this result evidences that the scale-dependent feedbacks present in the RV model are essential for the emergence and maintenance of the boom-bust cycles, and non-trivially affect the spatial distribution of consumers which abandon a scale-free distribution to adopt a more regular and dynamic one.

##### Without behavioural modulation

We ran simulations in which we forced the behavioural parameter *γ* for both consumers to either *γ* = 0 (purely advective behaviour) or *γ* = 1 (purely diffusive behaviour). Forcing behaviour to exclusively advection or exclusively diffusion leads in general to a reduced survivorship for the ecosystem due to predator extinction and the consequent lack of top-down controls for grazers (see above). For the SFV model, surviving replicates still show scale-invariant spatial distribution of vegetation (for *a* = *a*_*c*_) and consumers (for a range of *a* that includes *a*_*c*_), as when behavioural modulation is allowed; differently from the complete case, however, the predator generic scale invariance seems more limited (i.e. happening for a narrower range of *a*) and, importantly, trophic cascades disappear (Fig. S.15). For the RV model, the lack of behavioural modulation also leads to a reduced survivorship, with advection proving to be much more important to that end than diffusion; specifically, although in both cases boom-bust cycles are observed, when *γ* = 1 (purely diffusive case) ecosystem survival only happens when trophic dynamics are very slow, i.e. close to the desertification transition (see main text).

##### Arbitrary perception ranges

We also ran simulations by forcing a specific value for the perception radii, that is, by replacing Eqs.(8) with a fixed number that is half the values used in our main simulations, or twice the value. As a result, we still obtained scale-free cluster size distributions with the SFV model, and boom-bust cycles with the RV model. In the latter, larger perception radii led to larger areas being simultaneously grazed or predated upon, and thus more noticeable and more coordinated trophic waves.

##### Different consumption rates

To study the effects of consumer pressure, we tried four different combinations of scalings for the allometric grazer and predator attack rates (i.e. *a*_*V G*_ and *a*_*GP*_, respectively, taking values 1x, 25x the value in Table S.2). The SFV model was barely affected. For the RV model, we obtained broadly similar dynamics (i.e. the emergence of dynamic boom-bust cycles with high, oscillatory synchrony, see Fig. S.6) for most of the 36 possible combinations of the above rates and heterogenous initial conditions, except for the 1x scaling of both *a*_*V G*_ and *a*_*GP*_, which resembled the vegetation-only case, see SI Fig. S.11. Thus, and as expected, decreasing sufficiently such a maximum consumption rate led to negligible consumption and therefore the loss of e.g. the boom-bust cycles for the RV model, with vegetation qualitatively reverting to the non-consumer case (Fig. S.11).

##### Different initial conditions

For the RV model, we used a heterogeneous initialisation in which an initial group of clumps is separated by a certain distance. For this model, we used the initial configuration to study the effect of different initial characteristic scales and different degrees of initial ecosystem degradation. Thus, we tried configurations in which vegetation clumps of 400 *m* were spaced either 1, 2 or 3 *km* apart, and in which we randomly removed 5%, 10% or 50% the initial clumps. Figure S.7 summarizes the combined effect of rainfall and different initial conditions, for two choices of predator attack rates. We observed coexistence with boom-bust dynamics across most of these configurations (see Fig. S.6 for a comparison of synchrony results across some of these configurations). In short, sparser initial vegetation layouts, which correspond to increasing initial difficulty of successful grazer movement, led to increased coexistence, as did increased predator attack rates and higher rainfall levels. For the SFV model, we initialised simulations homogenously, that is, with constant density of all species at all grid sites. This initialization is customary in the vegetation-only SFV model, and facilitates the study of vegetation density decay over time. Nonetheless, when using the heterogenous initialisations discussed above for the RV model, we still observed the emergence of fractality and scale-free structure.

### B.2 Derivation of Linear Langevin Equations

For an arbitrary, non-negative 2D density variable *ρ* (*ρ* ≥ 0 everywhere) the first-order discretised diffusion stencil 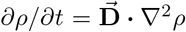 can be split into a constant positive term 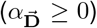 and a linear term 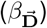 at each spatial site (*i, j*) as follows:

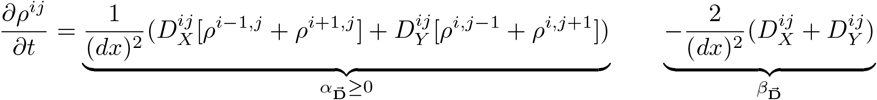

where 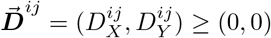 necessarily.

Similarly, we can decompose any arbitrary advection process for the variable, 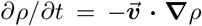 (where 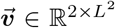), through the Upwind Finite Difference Method (UFDM) scheme to generically yield a constant positive term 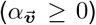 and a linear term 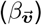. Thus, at each spatial site (*i, j*) and for any arbitrary direction of 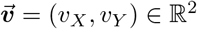, we can construct the following first-order stencil:

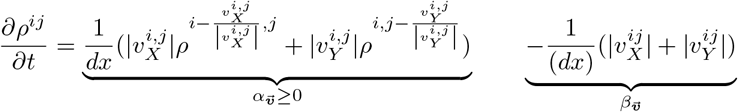

As the UFDM is one-sided (LeVeque 2002), the generic decomposition above ensures convergence 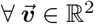, subject to the CFL condition in Eq. (B.2c).

The vegetation equations only involve the diffusion stencils, which leads to Eqs. (B.3a)-(B.3b). On the other hand, if we generically describe our consumer SDEs (Eqs. (2)–(3)) as follows (i.e. splitting them into linear and non-linear components):

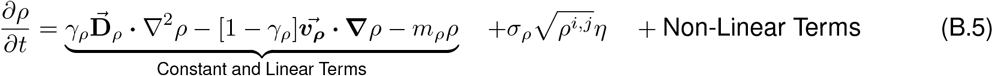

we can use the stencils above to arrive at the discretised Langevin corresponding to 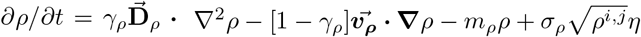:

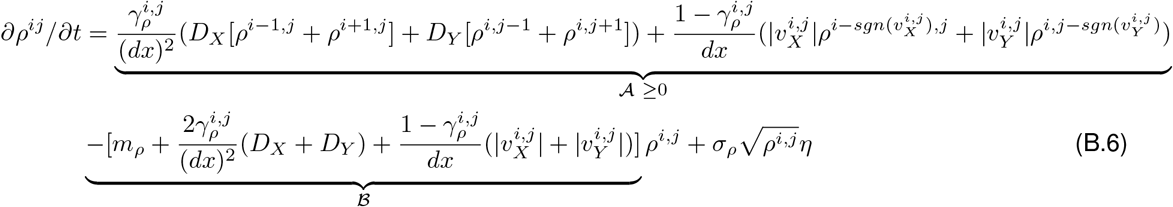

Note that this equation differs from (B.2a) in the correction factor for the excess diffusion that is introduced by the UFDM scheme, which we derive in the following sub-section.

### B.3 Derivation of Correction Factor for Excess Diffusion

Consider a 2*D* variable *q* subject to an advective velocity 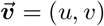, where *u* and *v* are constant scalars over some arbitrary time-interval [*t, t* + Δ*t*). We can write the classic advection equation as follows:

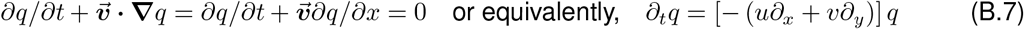

Using that 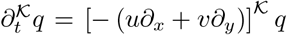 for any *K* ⩾ 1 (noting the commutativity of the partial derivatives), for *K* = 2:

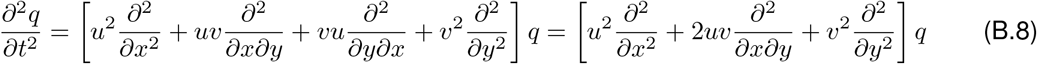

Then, using the notation 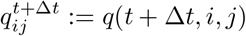, a simple Taylor expansion around *q*(*t, i, j*) yields:

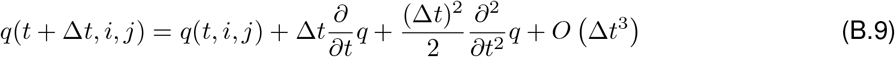

or, using the suprascript notation:

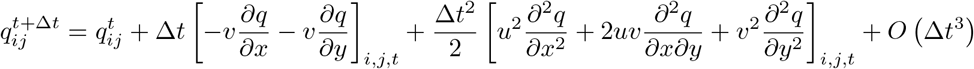

Using Eqs. (B.7) and (B.8):

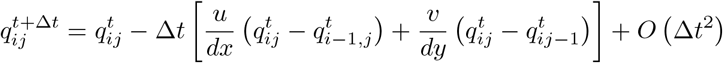

Discretising 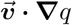 using UFDM and taking *dx* = *dy*:

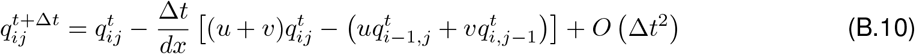

Further Taylor expansions around 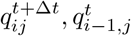 and 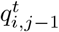 then yields:

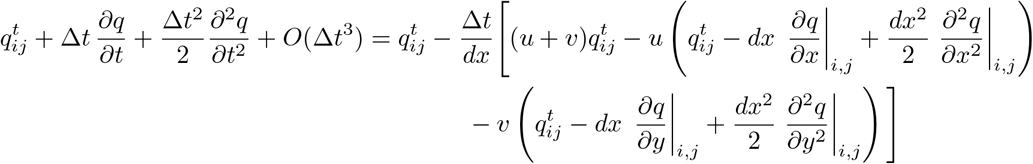

Using Eq. (B.8) and ignoring 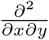 and higher order terms:

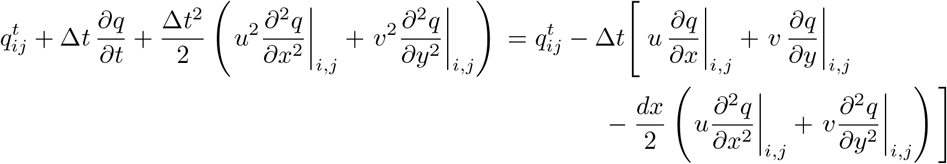

This expression provides the excess (extranumerary) diffusion coefficient factor coefficient 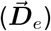:

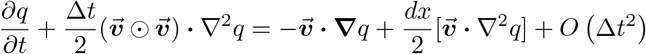

and rearranging:

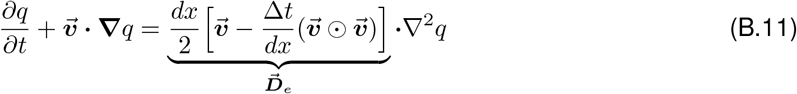

where ⊙ denotes the Hadamard product.

Thus, solving Eq. (B.7) by a UFDM discretisation of 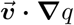 actually solves the advection-diffusion equation Eq. (B.11) with the extranumerary diffusion coefficient 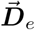. It is for this reason that we used the correction factor, scaled by the advection probability 1 − *γ*, as noted in Eq. (B.2b).

## C Spatial Pattern Analyses

Detecting the presence of spatial patterns, particularly scale-free patterns in consumers, required a classification algorithm able to reliably distinguish ‘high-density’ regions (‘active’ clusters) from ‘low-density’ regions. Thus, we used Gaussian Mixture Models (GMM) with 2 components, implemented using *sklearn*.*mixture*.*GaussianMixture* in Python. This method uses an Expectation-Maximisation (EM) algorithm to fit a mixture of two Gaussians to the snapshots to be analysed. We chose GMMs over popular alternatives such as KMeans because GMMs do not assume spherical cluster covariances in the underlying data, and therefore produce a better classification of irregularly shaped components in the frames (Biernacki, Celeux, and Govaert 2000).

To measure cluster sizes *s* from these classified snapshots, we used the two-pass Hoshen-–Kopelman algorithm, implemented using *scipy*.*ndimage*.*label* in Python, a standard union-finding method (Hoshen and Kopelman 1976). We then measured the Complementary Cumulative Distribution Function (CCDF) (*P*[*S* ≥ *s*]), which facilitates the identification of whether the distribution is scale-free because it avoids binning artefacts common to simpler cluster size probability distribution plots (*P*[*s*]) (Clauset, Shalizi, and Newman 2009). The CCDF allowed for the estimate of the desertification threshold *a*_*c*_ in the DP model, where the probability distribution of active (i.e. non-zero) vegetation cluster sizes follows a power-law (i.e. *P*[*S*] ∝*S*^−*τ*^ or, equivalently, *P*[*s* ≥*S*] ∝*S*^1−*τ*^). In other words, for the critical threshold of vegetation growth rate *a*_*c*_ separating the barren and vegetated states, this distribution shows a power law behaviour that saturates for *a > a*_*c*_ and declines exponentially to zero for *a < a*_*c*_ (Muñoz et al. 1999)

We measured regularity in classified snapshots by first converting them to frequency space using discrete 2*D* Fast Fourier Transforms (FFT). We implemented the latter using assorted *numpy/cupy* functions in Python. We then radially averaged the resulting frequency space 2D peridograms over concentric circular bands of increasing radii, to arrive at the radially averaged power spectrum (Kéfi et al. 2014; Chen, Halder, and Bonachela 2024). Peaks in this power spectrum indicate regularity in the frames. For more details about the methodology, refer to ref. (Chen, Halder, and Bonachela 2024).

## D Synchrony Analyses

We broadly measured spatial similarity among snapshots of the stationary state of the system using three different metrics: the maximum normalised cross-correlation, the Pearson correlation coefficient, and the adjusted mutual information. The first two metrics required the calculation of the time-lagged Zero-mean Normalised Cross Correlation (ZNCC) matrix, *C* ^Δ*T*^, while the latter represents an entropic measure (Vinh, Epps, and Bailey 2010).

### D.1 Zero-Normalised Cross Correlation (ZNCC)

For a reference snapshot *ℳ*_0_ and another snapshot *ℳ*_Δ*T*_ taken after time Δ*T* (both of sizes *L ×L*), the time-lagged ZNCC matrix *C*^Δ*T*^ is a real matrix of size (2*L* −1) *×* (2*L* −1), with its elements given by: (Zhao, Sang, and Duan 2019)

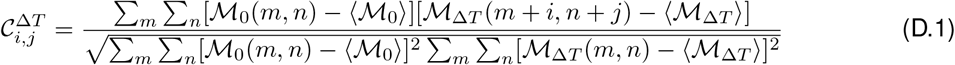

*where i < L* − 1 & *j < L* − 1.

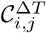 represents the cross-correlation when the origins of these two frames are spatially shifted by *I* and *j* grid-spacing units respectively. The first element 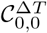 (representing spatial overlap) is equivalent to the 2D Pearson correlation coefficient, whereas 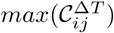 represents the maximum cross-correlation across all potential overlaps, thereby identifying shifted patterns (Zhao, Sang, and Duan 2019).

Grayscaling the respective snapshots (i.e. normalising them to snapshots with zero mean and unit variance as *M*:= (−*ℳ* ⟨*ℳ*⟩)*/σ*_*ℳ*_) allows for comparisons between frames with different ‘brightness’ or ‘intensities’ (specifically in our case, it allowed for comparisons between different species with different densities). Additionally, it simplifies the above relation and allows for the direct application of the circular convolution theorem:

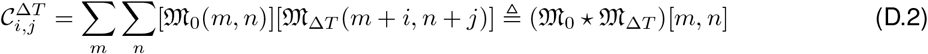

where ⋆ represents 2*D* cross-correlation.

We calculated this time-lagged matrix *C*^Δ*T*^ using 2*D* convolution, implemented using *scipy/ cupyx*.*signal*.*fftconvolve* in Python.

### D.2 Adjusted Mutual Information (AMI)

We further measured the Adjusted Mutual Information (AMI), which is an entropic metric (Vinh, Epps, and Bailey 2009; Vinh, Epps, and Bailey 2010). Unlike correlation-based measures such as *C* ^Δ*T*^, which quantify linear relationships between pixel intensities, the AMI captures the general statistical dependence between two spatial distributions without assuming any particular functional form. This makes it particularly advantageous for detecting non-linear spatial correspondences and complex pattern similarities that may not be adequately captured by correlation metrics alone.

The calculation of the AMI requires the calculation of mutual information (MI). MI quantifies the amount of information about one random variable obtained by observing another. For two grayscaled snapshots *M*_0_ and *M*_Δ*T*_ (normalised to zero mean and unit variance as described in Section D.1), we first discretised the continuous pixel intensity values into histogram bins to obtain discrete partitions. The mutual information between these partitions is then defined as:

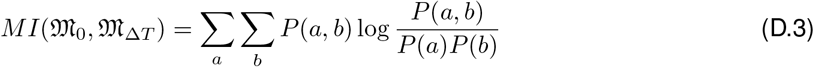

where *P*(*a*) and *P*(*b*) are the marginal probability distributions of the binned pixel intensities in *M*_0_ and *M*_Δ*T*_ respectively (obtained from their individual histograms), and *P*(*a, b*) is their joint probability distribution (constructed from the two-dimensional histogram of co-occurring intensity values). Intuitively, MI measures how much knowing the intensity distribution in one frame reduces uncertainty about the intensity distribution in the other frame. When the two frames are independent, *P*(*a, b*) = *P*(*a*)*P*(*b*), and thus *MI* = 0.

MI has, however, a significant limitation: its magnitude is not bounded and depends on both the number of bins used to construct the histograms and the size of the dataset (Vinh, Epps, and Bailey 2010). This dependence makes it difficult to meaningfully compare MI values across different discretisation schemes, different frame sizes, or even between different pairs of snapshots if the effective information content varies. Consequently, raw MI values lack a natural interpretation of what constitutes ‘high’ or ‘low’ similarity (Vinh, Epps, and Bailey 2009). The Adjusted Mutual Information addresses these limitations by normalising MI and correcting for chance agreement between partitions (Vinh, Epps, and Bailey 2009; Vinh, Epps, and Bailey 2010).

The AMI (*a*) is defined as:

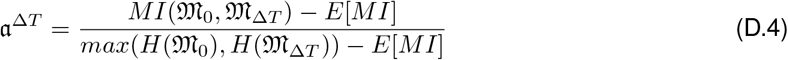

where *E*[*MI*] is the expected mutual information under a null model that assumes random permutation of the partitions (i.e., the baseline MI expected by chance alone), and *H*(*·*) denotes the Shannon entropy of a partition. The denominator normalises the adjusted information by the maximum possible entropy, ensuring that AMI is bounded. This normalisation guarantees that AMI equals 1 for perfect agreement between the two frames, 0 for chance-level agreement (no more similar than random permutations would produce), and can take negative values for agreements worse than random chance (Vinh, Epps, and Bailey 2010). The adjustment for chance is particularly important when comparing snapshots with varying levels of intrinsic structure or entropy.

To compute the AMI between time-separated snapshots, we first normalised both *ℳ*_0_ and *ℳ*_Δ*T*_ to grayscaled frames *M*_0_ and *M*_Δ*T*_ with zero mean and unit variance, following the same procedure described in Section D.1. We then constructed histogram partitions for both frames to discretise the continuous pixel intensity distributions. The number of bins for these histograms was determined using Scott’s rule (Scott 1979), which estimates the optimal bin width based on the data’s standard deviation and sample size to balance between under-smoothing and over-smoothing the underlying distribution. We capped the maximum number of bins at 2*L* to prevent excessive discretisation that could lead to sparse histograms with unreliable probability estimates. Finally, we calculated the AMI using *sklearn*.*metrics*.*adjusted_mutual_info_score* in Python, which implements the formulation given in Eq. (D.4) and handles the computation of both the expected MI under random permutation and the normalisation factor.

### D.3 Synchrony Time-Series Analysis

Using our spatial similarity metrics 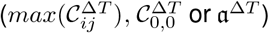, generically denoted here as *ℳ*^Δ*T*^, we constructed the corresponding synchrony time series by measuring *ℳ*^ΔT^ between a reference grayscaled snapshot *ℳ*_0_ (measured at *t* = 8 *·*10^4^ hr) and another snapshot *ℳ*_Δ*T*_ separated in time by Δ*T*. We sampled Δ*T* at 20 hr intervals between *t* and *t* + 6000 hr (*t* + 5000 hr for scale-free simulations). For each initialisation condition, and for each choice of aridity level (growth rate/rainfall level) we generated 100-120 independent *ℳ*^ΔT^ timeseries replicates. We measured periodicity in the resulting time series by performing, using *scipy*.*signal*.*find_peaks* in Python, a 1*D* FFT and selecting the primary peak (if it exists) as long as it is at least 10% the maximum signal (FFT) value detected by the algorithm.

Additionally, from each time series, we measured the probability distributions obtained with the values of *ℳ*^ΔT^, from which we excluded the initial Δ*T* = 200 *hr* transients to ensure that initial correlations do not bias the results. Thus, we tested whether these synchrony time series distributions (i.e. synchrony strengths) increased with rising aridity (i.e. lower vegetation growth rates/rain-fall levels) using the non-parametric Mann–Whitney U / Wilcoxon rank-sum test, implemented using *scipy*.*stats*.*mannwhitneyu* (Mann and Whitney 1947). If 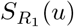 and 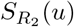 are the survival functions of the distributions underlying the lowest aridity level (*R*_1_) and the highest aridity level (*R*_2_), we tested the null hypothesis 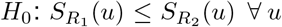, against the alternative hypothesis that the distribution at the lowest aridity level is higher than that at the highest aridity level, i.e.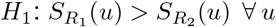.

## Supplementary Figures and Tables

**Figure S.1:**
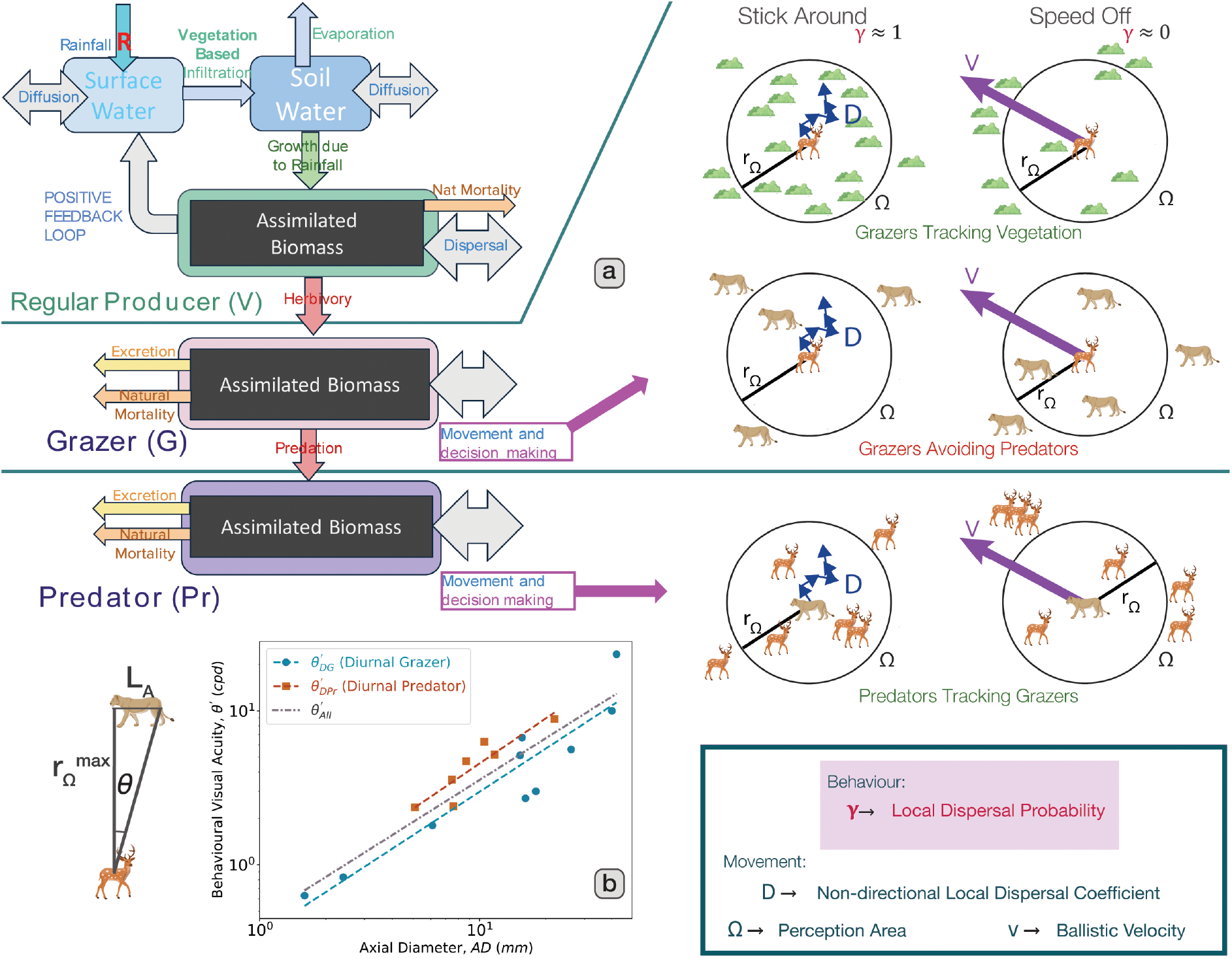
Schematic representation of the complete model. **a** Energy flow in the consumer-resource model developed here for the regular case (see MM for details). Rainfall (*mm/hr*) controls plant growth as it seeps into the soil and is then taken up by existing plant growth. More plant density at a site leads to a short-scale positive feedback, while root systems compete for water creating a long-scale negative feedback. Plants can disperse slowly, with plant density removed by grazing or natural mortality. Consumers grow through consumption, die of natural mortality (and, for grazers, additionally predation), and forage (or, for grazers, avoid predation). For the latter, consumers switch between local, non-directional dispersal and directional advection, considered here through the local dispersal probability, *γ*. Consumers track resources within their perception area Ω, with grazers also avoiding predators. The stronger the cues to leave via advection (e.g. low resource or, for the grazer, high predator density), the lower the value of their local dispersal probability *γ*. **b** *(Left)* Illustration representing the approximate relationship between the behavioural visual resolution *θ* (in), the length of the perceived animal (*L*_*A*_, Eq. (A.2)), and the maximum perception radius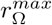. *(Right)* Allometry for diurnal consumer behavioural visual acuity (*θ*^*′*^ in *cpd*) as a function of the axial eye diameter (AD, in *mm*) (Eq. (A.3b)). Diurnal grazers are denoted by red squares, diurnal predators by blue circles. Data derived from (Veilleux and Kirk 2014).

**Figure S.2:**
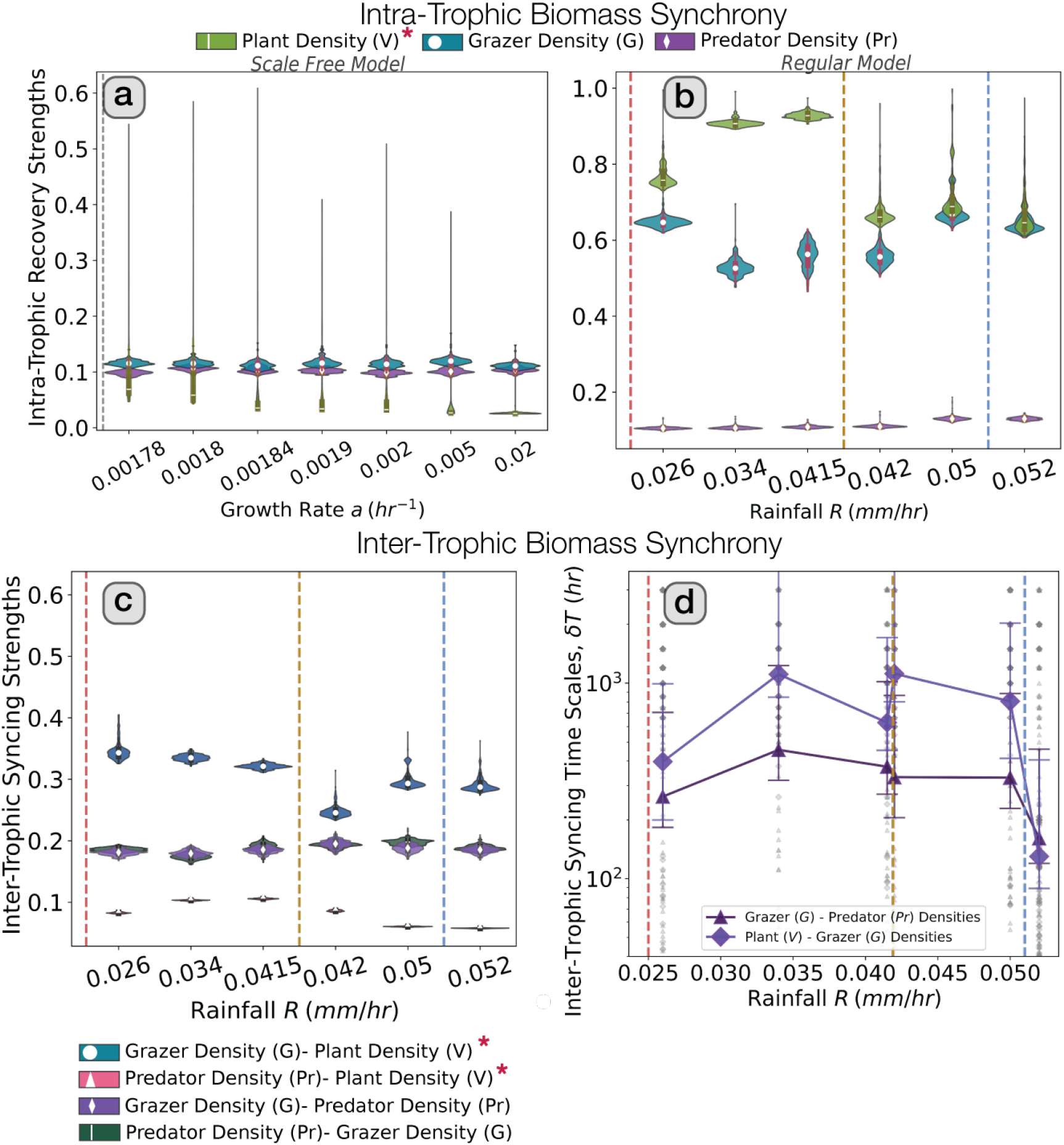
Changes in intra-trophic (recovery) and inter-trophic (interaction/tracking) synchrony of biomass densities with aridity (i.e. with proximity to desertification). For all panels, synchrony is measured using the maximum spatial correlation 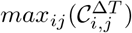; red asterisks (*) in the legend indicate synchrony strengths that rise significantly with rising aridity (Mann-Whitney U rank test, *p <* 0.05). The grey vertical dashed line in panel a indicates the desertification threshold (≈ 0.001728 *hr*^−1^); for panels b-d, dashed vertical lines indicate the desertification threshold (*red*, ≈ 0.024 *mm/hr*), the end of bistability (*yellow*, ≈ 0.0419 *mm/hr*) and the transition to homogenous (i.e. non-patterned) vegetation (*light blue*, ≈ 0.051 *mm/hr*) in the vegetation-only model. **a**-**b** Intra-trophic recovery strengths in population biomasses in the scale-free case (panel a) and regular case (panel b); in both cases, there is a significant rise in vegetation maximum auto-correlation strengths as the desertification thresholds are approached. **c** Inter-trophic interaction strengths between population densities in the regular case; the synchrony between grazer and vegetation densities, and between predator and vegetation densities, rise significantly when rainfall levels are close to the desertification threshold. **d** Interaction time between population densities, corresponding to the time period of the synchrony oscillations in inter-trophic population biomass for the regular vegetation model.

**Figure S.3:**
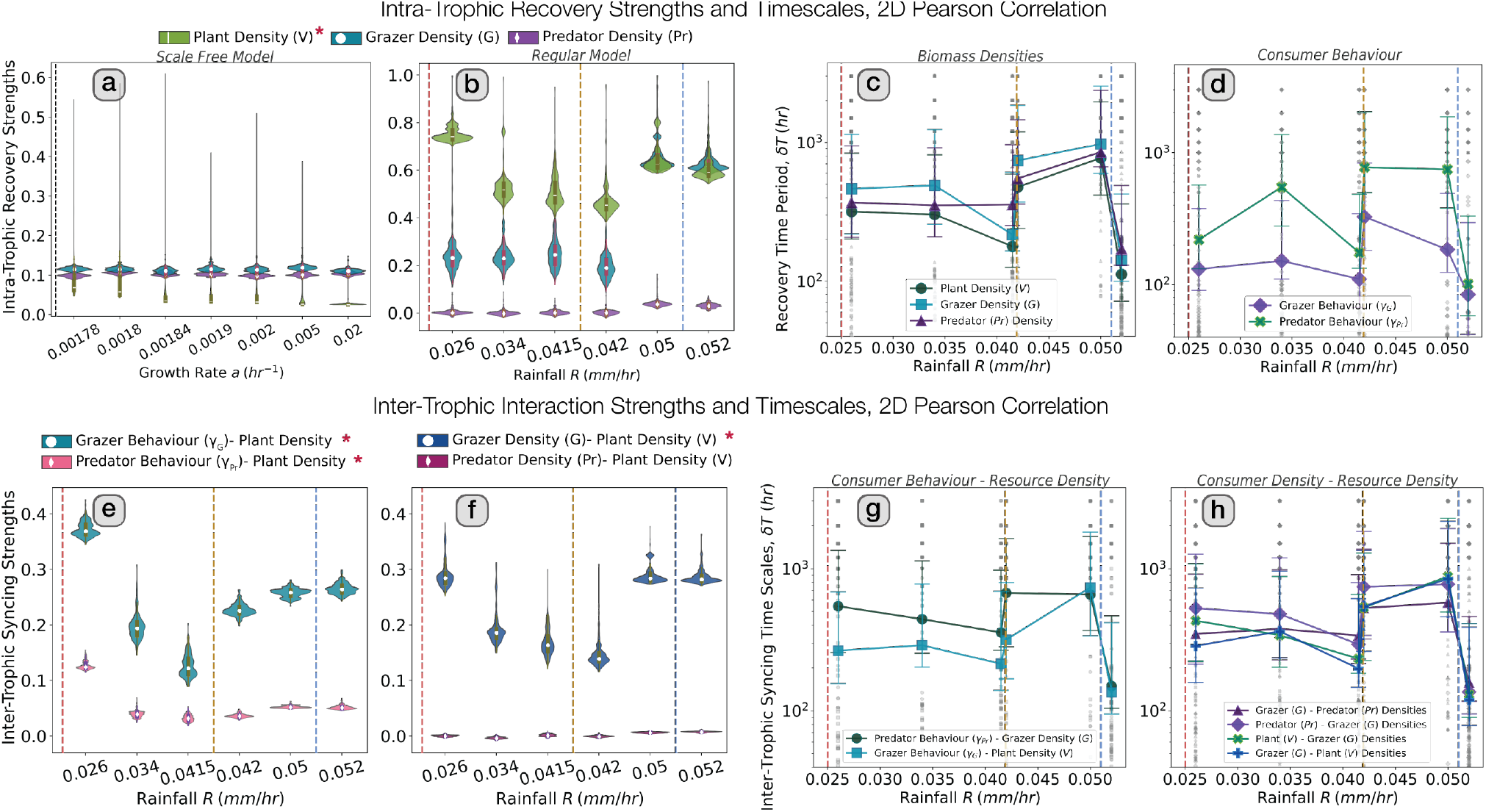
Changes in intra-trophic (recovery) and inter-trophic (interaction/tracking) synchrony of biomass densities and consumer behaviour with aridity. For all panels, synchrony is measured using the 2*D* Pearson Correlation Coefficient, 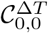; red asterisks (*) in the legend indicate synchrony strengths that rise significantly with rising aridity (Mann-Whitney U rank test, *p <* 0.05). The grey vertical dashed line in panel a indicates the desertification threshold (≈ 0.001728 *hr*^−1^); for panels b-h, dashed vertical lines indicate the desertification threshold (*red*, ≈ 0.024 *mm/hr*), the end of bistability (*yellow*, ≈ 0.0419 *mm/hr*) and the transition to homogenous (i.e. non-patterned) vegetation (*light blue*, ≈ 0.051 *mm/hr*) in the vegetation-only model. **a**-**b** Intra-trophic recovery strengths in population biomasses in the scale-free case (panel a) and regular case (panel b); in both cases, there is a significant rise in vegetation maximum auto-correlation strengths as the desertification thresholds are approached. **c**-**d** Recovery time (i.e. period of oscillations) in the synchrony of intra-trophic population biomass (panel c) and consumer behaviour (panel d) for the regular case. **e**-**f** Inter-trophic interaction strengths between consumer behaviour and population densities (panel e) and between population densities (panel F) in the regular case; the synchrony between predator or grazer behaviour and vegetation density, between grazer and vegetation densities, and between predator and vegetation densities, rise significantly when rainfall levels are close to the desertification threshold. **g**-**h** Interaction time between consumer behaviour and resource density (panel g) and between consumer densities and resource densities (panel h), corresponding to the time period of the oscillations in the respective consumer-resource synchrony plot for the regular vegetation model.

**Figure S.4:**
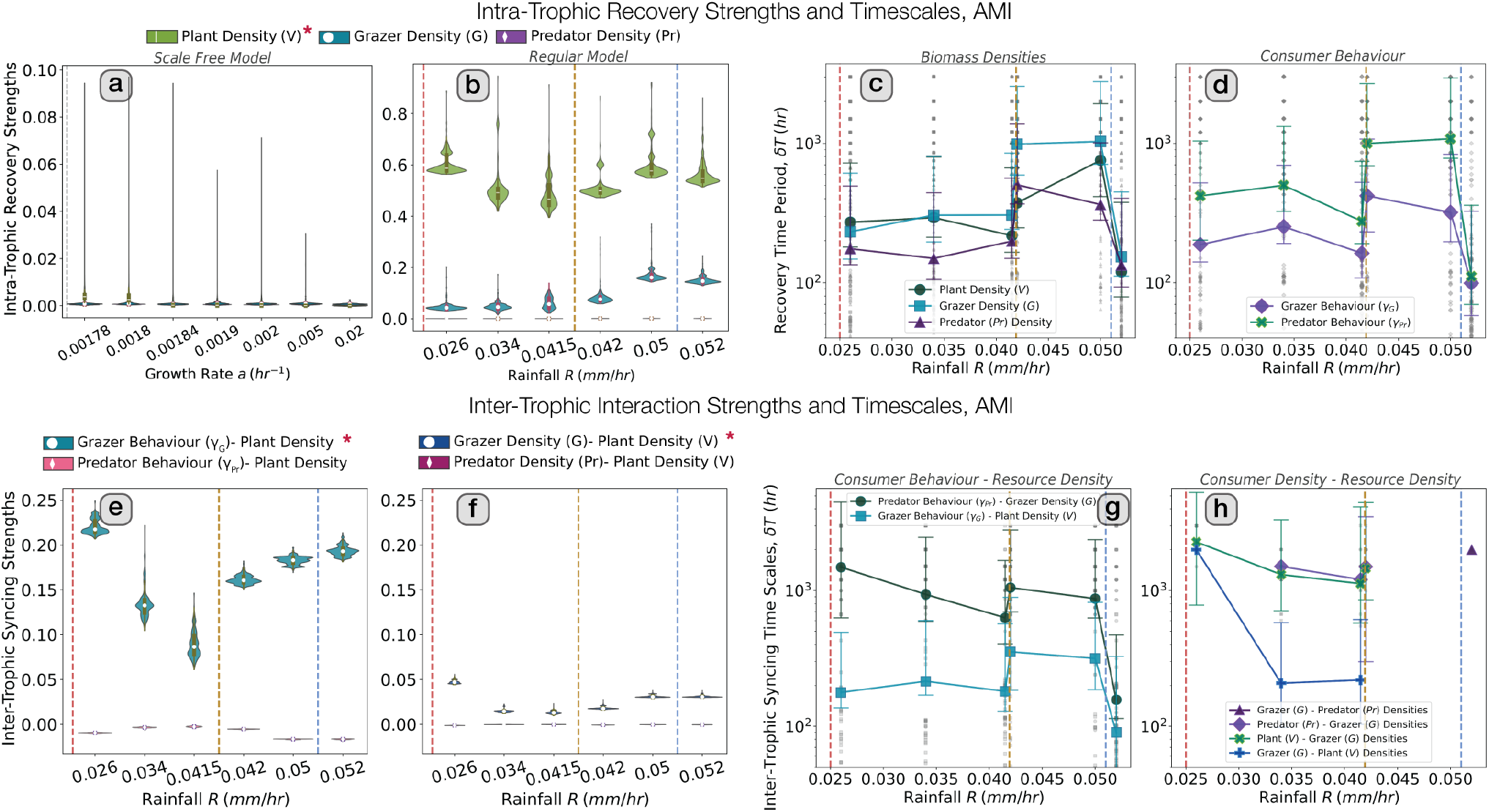
Changes in intra-trophic (recovery) and inter-trophic (interaction/tracking) synchrony of biomass densities and consumer behaviour with aridity. For all panels, synchrony is measured using the Adjusted Mutual Information (AMI) index, *a*^Δ*T*^; red asterisks (*) in the legend indicate synchrony strengths that rise significantly with rising aridity (Mann-Whitney U rank test, *p <* 0.05). The grey vertical dashed line in panel a indicates the desertification threshold (≈ 0.001728 *hr*^−1^); for panels b-h, dashed vertical lines indicate the desertification threshold (*red*, ≈ 0.024 *mm/hr*), the end of bistability (*yellow*, ≈ 0.0419 *mm/hr*) and the transition to homogenous (i.e. non-patterned) vegetation (*light blue*, ≈ 0.051 *mm/hr*) in the vegetation-only model. **a**-**b** Intra-trophic recovery strengths in population biomasses in the scale-free case (panel a) and regular case (panel b); in both cases, there is a significant rise in vegetation maximum auto-correlation strengths as the desertification thresholds are approached. **c**-**d** Recovery time (i.e. period of oscillations) in the synchrony of intra-trophic population biomass (panel c) and consumer behaviour (panel d) for the regular case. **e**-**f** Inter-trophic interaction strengths between consumer behaviour and population densities (panel e) and between population densities (panel f) in the regular case; the synchrony between grazer behaviour and vegetation density rise significantly when rainfall levels are close to the desertification threshold. **g**-**h** Interaction time between consumer behaviour and resource densities (panel g) and between consumer and resource densities (panel h), corresponding to the time period of the oscillations in consumer-resource behavioural and density synchrony for the regular vegetation model.

**Figure S.5:**
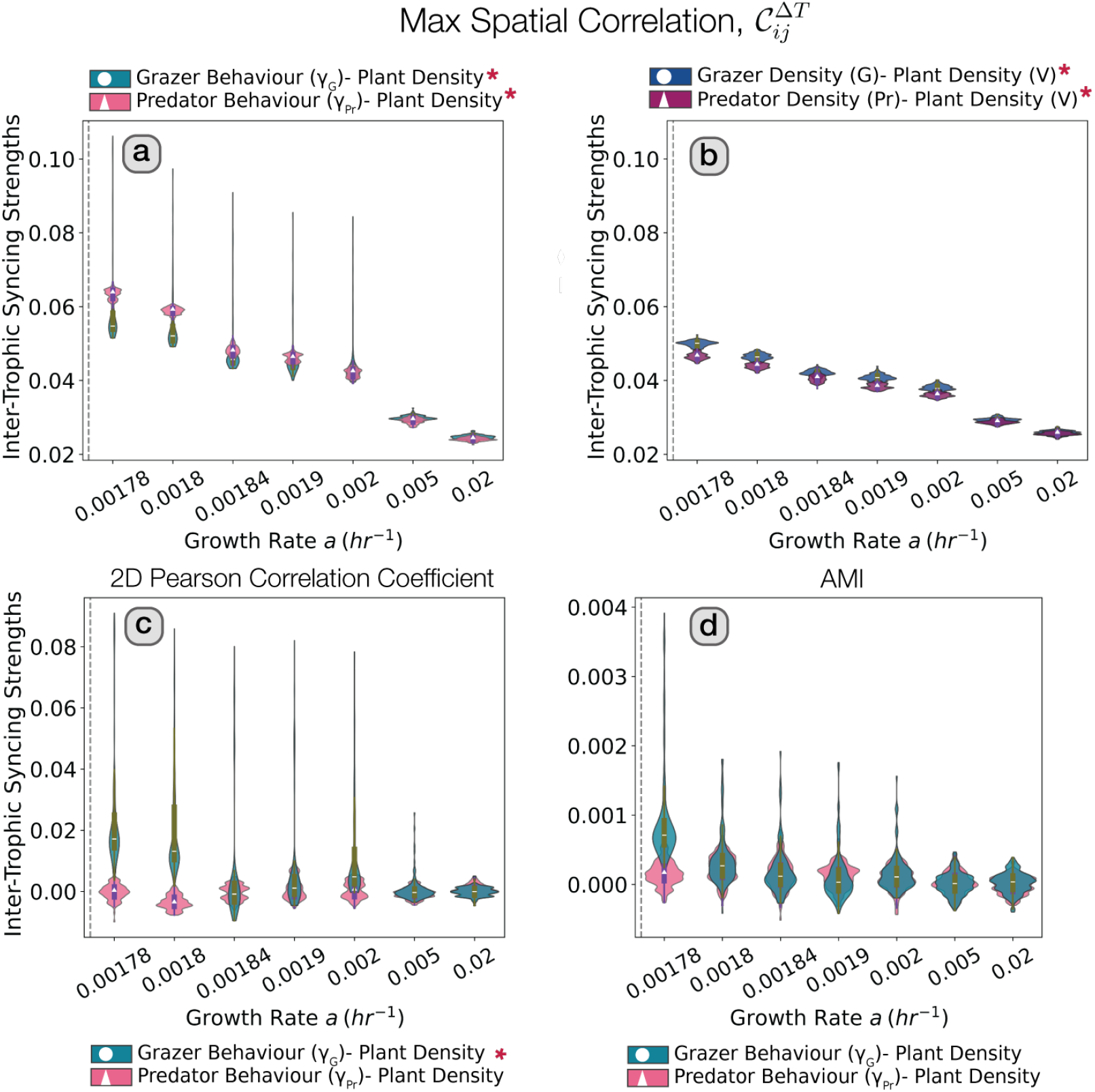
Changes in inter-trophic (interaction/tracking) synchrony of consumer behaviour and biomass densities with aridity in the scale-free model, across the different spatial similarity metrics. Red asterisks (*) in the legend indicate synchrony strengths that rise significantly with rising aridity (Mann-Whitney U rank test, *p <* 0.05); dashed vertical line represents the desertification threshold (≈ 0.001728 *hr*^−1^). **a**-**b** Inter-trophic interaction strengths between consumer behaviour and population densities (panel a) and between population densities (panel b), measured using the maximum spatial correlation,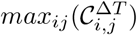. **c**-**d** Inter-trophic interaction strengths between consumer behaviour and population densities, measured using the 2D Pearson Correlation Coefficient, 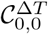 (panel c) and the Adjusted Mutual Information (AMI) index, *a*^Δ*T*^ (panel d); for either choice of metric, inter-trophic synchrony strengths between population densities were negligible (i.e. population densities were asimilar).

**Figure S.6:**
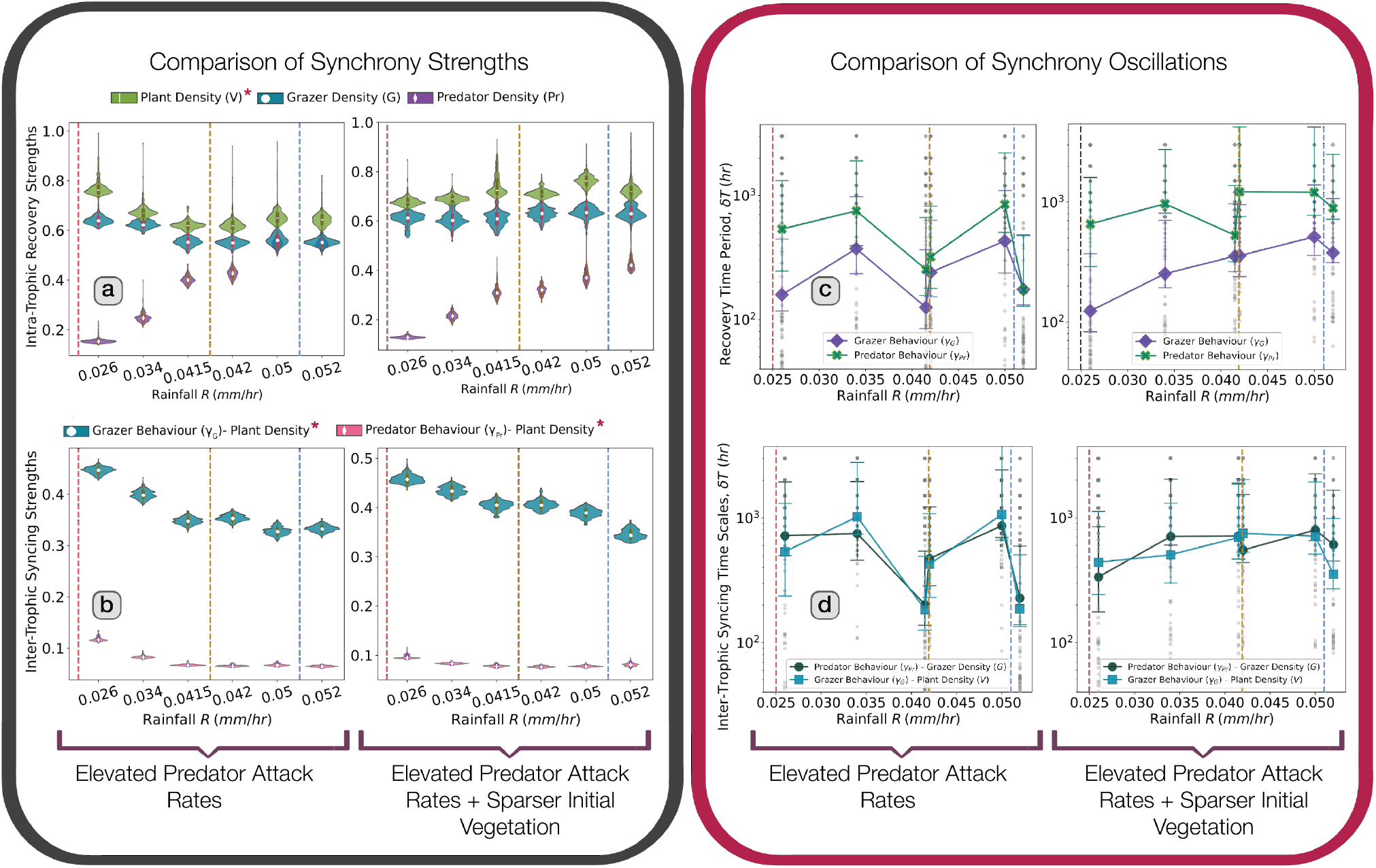
Sensitivity analysis for the effect of attack rate and initial condition on synchrony characteristics for the regular vegetation (RV) tri-trophic model. For all panels, synchrony strengths and time periods are measured using the maximum spatial correlation metric,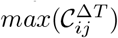. For each box, all panels result from setting attack rates (*a*_*V G*_ and *a*_*GP r*_) to 25x the allometric scaling relationship given in *Table S*.*2*; left panels used initial conditions in which clumps of size 400 *m* separated by 2 *km* are regularly distributed in space and 10% of initial clumps are randomly removed, whereas right panels used sparser initial vegetation clumps (50% of initial vegetation clumps randomly removed). Dashed vertical lines indicate the desertification threshold (*red*, ≈ 0.024 *mm/hr*), the end of bistability (*yellow*, ≈ 0.0419 *mm/hr*) and the transition to homogenous (i.e. non-patterned) vegetation (*light blue*, ≈ 0.051 *mm/hr*) in the vegetation-only model. **a**-**b** Intra-trophic recovery strengths in population biomasses (panels a) and inter-trophic interaction strengths between consumer behaviour and population densities (panels b). Auto-correlation/recovery of vegetation density, and synchrony of grazer behaviour with vegetation density, and predator behaviour with vegetation density rise significantly with increasing aridity across all analyses. The rise in inter-trophic synchrony are particularly robust, holding also for unit predator attack rate scalings (Fig. 4 c), and for most alternative metrics of spatial similarity (Figs. S.3 and S.4), and are all in agreement with the match-mismatch hypothesis (see main text and (Visser and Gienapp 2019)). **c**-**d** Recovery time of synchrony oscillations in intra-trophic consumer behaviour (panels c), and interaction time period between consumer behaviour and resource densities (panels d); the sparser initial vegetation layout disrupts the trend observed for the denser initial condition (see main text), particularly in the case of inter-trophic interaction time periods (d, right panel).

**Figure S.7:**
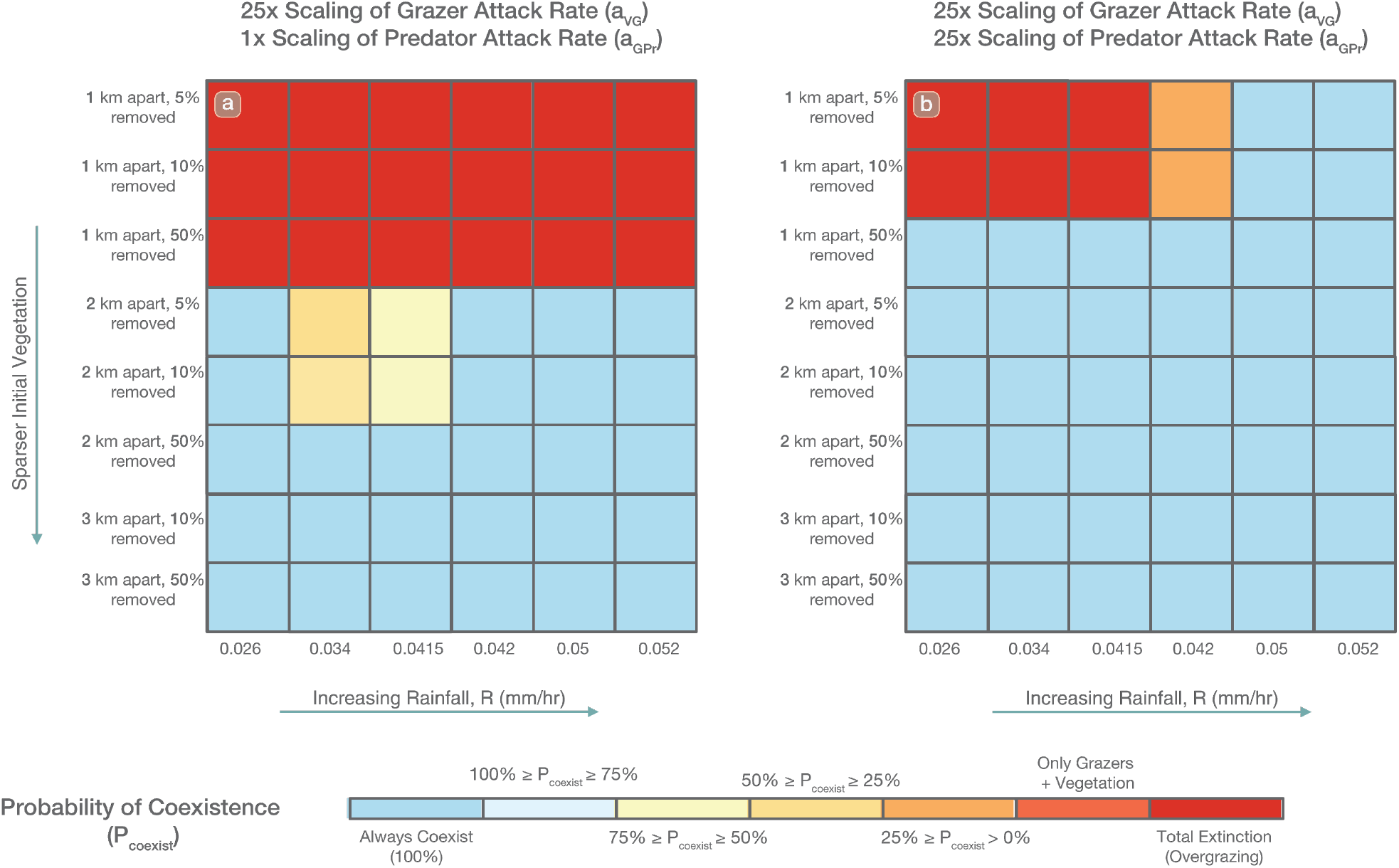
Sensitivity analysis for the effect of attack rate and initial condition on boom-bust coexistence for the regular vegetation (RV) tri-trophic model. All panels result from setting grazer attack rates (*a*_*V G*_ to 25x the allometric scaling relationship given in *Table S*.*2*. Predator attack rates (*a*_*GP r*_) were set to 1x (Panel **a**) and 25x (Panel **b**) of the allometric scaing relationship respectively, representing elevated predator attack rates in the second panel. In each scenario, rainfall levels ranged from 0.026*mm/hr* to 0.052*mm/hr* (x-axis), whereas initial vegetation layouts were arranged from denser to sparser layouts, comprising of vegetation clumps of size 400 *m* separated by 1, 2 or 3 *km* are regularly distributed in space and 5, 10% or 50% of initial clumps are randomly removed. Coexistence in all cases emerged from boom-bust dynamics. Broadly, sparser initial vegetation layouts, which correspond to increasing initial difficulty of successful grazer movement led to increased coexistence, as did increased predator attack rates (Panel **b**), which indicate predators ensure coexistence initially by restricting grazer tiling to specific vegetation clumps, allowing the rest to recover, and thus leading to the emergence of boom-bust dynamics. Increasing rainfall levels successfully mitigated the effects of initial ease of grazer movement (denser vegetation layouts, topmost rows) when coupled with high predator attack rates, thereby boosting coexistence (Panel **b**). For unit scaling of predator attack rates (Panel **a**), at vegetation layouts of intermediate density (initial separation of 2 *km* with 5, 10% randomly removed), the lowest rainfall levels close to the end of spatial bistability of the vegetation only model (Fig. 1D) helped stabilise boom-bust dynamics led coexistence, indicating potential slowing down effects.

**Figure S.8:**
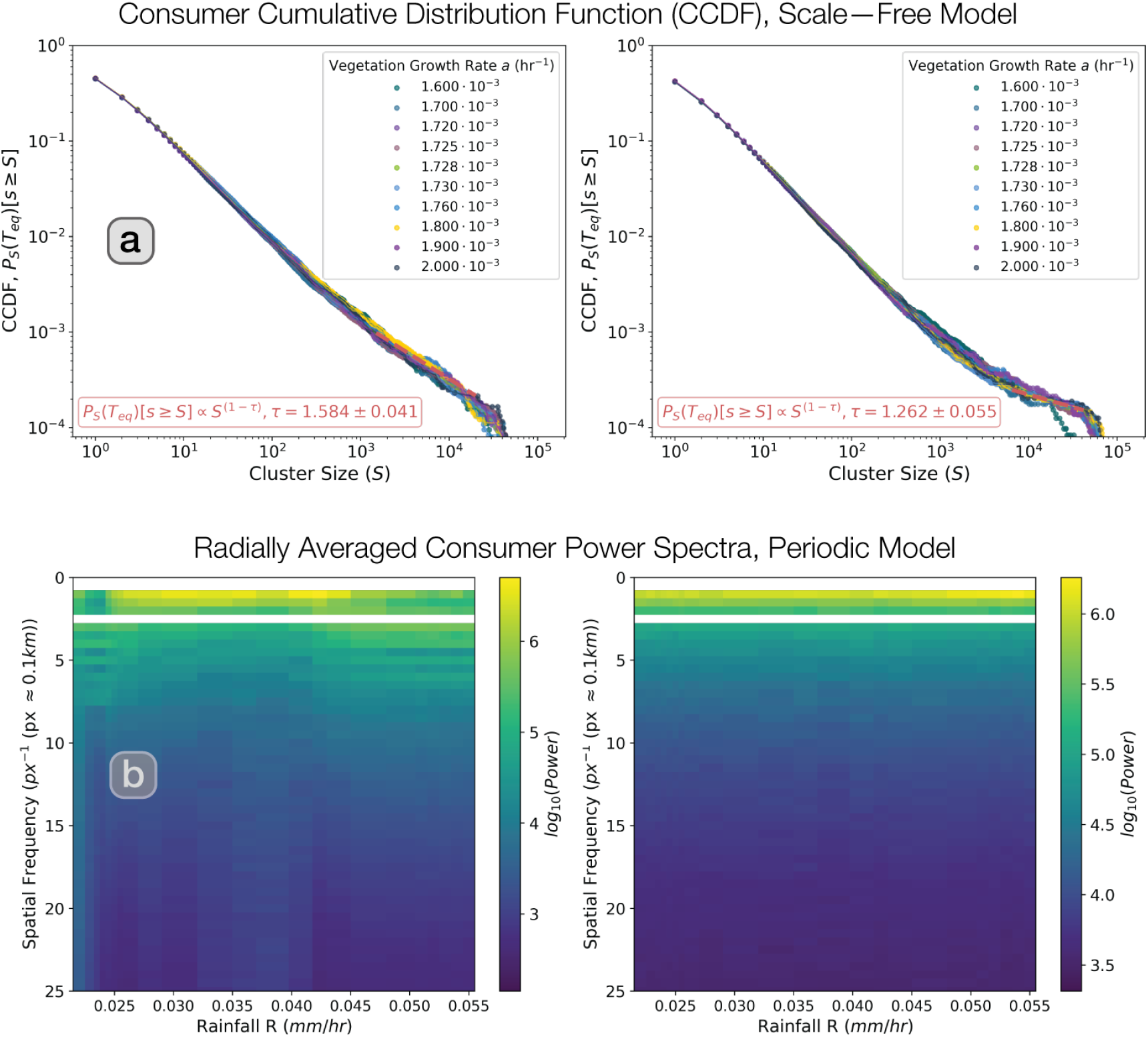
Spatial distribution of consumers in the tri-trophic models. **a** Complementary cumulative probability distributions (CCDF,, *P*[*s* ≥*S*]) for cluster size for grazers (left) and predators (right) in the tri-trophic scale-free model (*L* = 512*×* 512). The distributions show a power-law scaling (i.e. *P*[*s* ≥*S*] ∝ *S*^1−*τ*^) for a wide range of growth rates below and above the desertification threshold (*a*_*c*_ ≈ 1.728 *·*10^−3^ *hr*^−1^), **b** Heatmaps representing the radially-averaged power spectrum across different rainfall levels for grazers (left) and predators (right). The presence of peaks for all rainfalls (yellow bands in the heatmaps) indicates regularity in the corresponding spatial distributions.

**Figure S.9:**
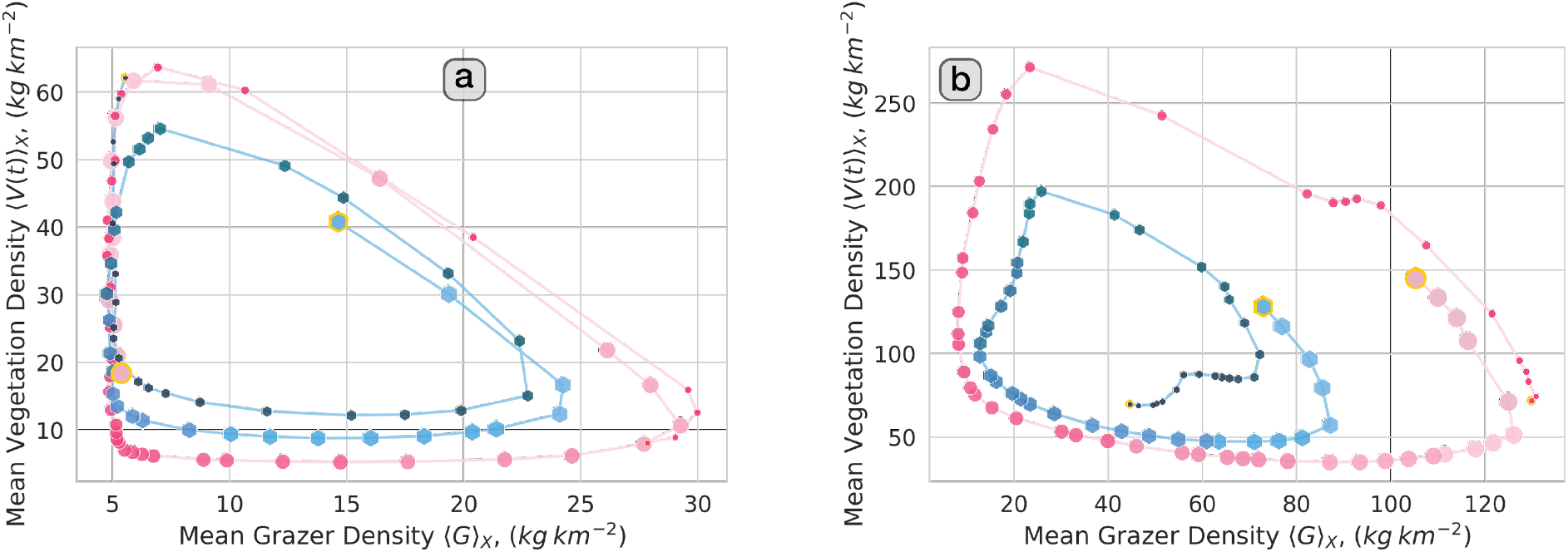
Apparent critical-slowing down in the regular vegetation model. **a**-**b** Phase portraits illustrating oscillatory trajectories for mean grazer and vegetation densities at two random replicates (blue and pink lines) before (*R* = 0.0415 *mmhr*^−1^, panel a) and after (*R* = 0.042 *mmhr*^−1^, panel b) the end of the bistability regime in the vegetation only-model. For clarity, only times between *t* ∈ [80000, 82000] *hr*) are shown, with lighter and larger points reflecting earlier times and smaller, darker points reflecting later times. Starting and final densities are highlighted in yellow. Oscillations are visibly faster at *R* = 0.0415 *mm hr*^−1^ (i.e. densities show more ‘loops’), indicating potential critical-slowing down effects (i.e. the phenomenon where a dynamical system’s recovery time from perturbations diverges as it approaches a bifurcation, manifesting as a weakening of the restoring force to perturbations due to loss of linear stability. This occurs because the eigenvalue governing the rate of return of the dynamical variables (in this case ⟨*G*⟩ _*X*_ and ⟨*V*⟩ _*X*_) to equilibrium approaches zero, causing phase-space trajectories to linger longer near the attractor – in other words a slowdown in phase-portrait oscillations as shown here (Strogatz 2015; Kéfi et al. 2014)).

**Figure S.10:**
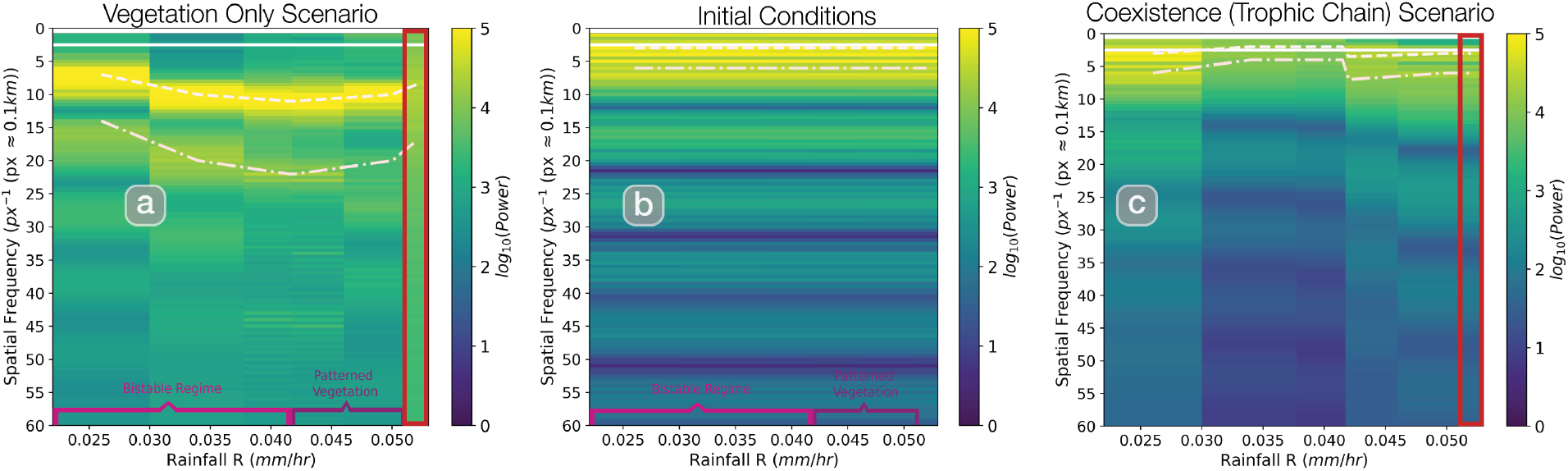
Same initial conditions lead to different regular vegetation patterns when consumers are considered. Panels depict r-spectra heatmaps across different rainfall levels for snapshots of vegetation at stationarity corresponding to the initial condition (middle panel), the vegetation-only case (left panel), and the complete model (right panel). Peaks in the radially averaged power spectra indicate periodicity, with the fundamental peak and the first harmonic denoted in dashed and dash-dotted off-white lines; the fundamental peak is a standard spatial Early-Warning Signal (EWS) used to predict desertification in regular landscapes (Kéfi et al. 2014; Chen, Halder, and Bonachela 2024). Although they start from the same initial condition (given by the middle panel), the regularity resulting from the complete model (right panel) is distinct from that of the vegetation-only model (left panel), more remarkably for high rainfall levels where patterns are lost in the vegetation-only model (red box).

**Figure S.11:**
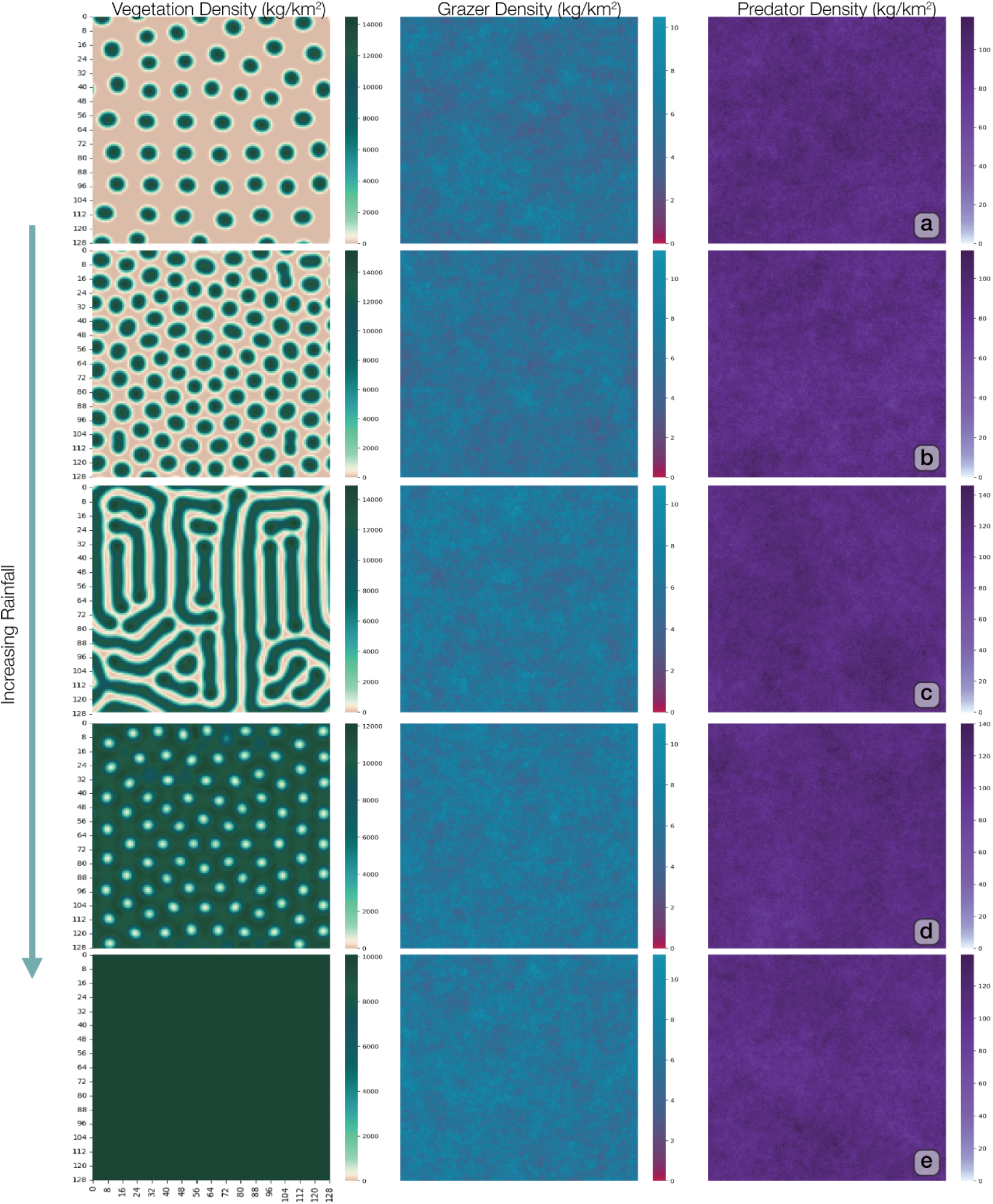
Representative snapshot frames of the tri-trophic model. (at *t* = 2 *·* 10^5^ *hr*) with regular vegetation and unit scaling of the allometric grazer attack rate (*a*_*GP r*_) as provided in Table S.2. As rainfall levels increase (from top to bottom, 0.026*mm/hr* (panel **a**), 0.034*mm/hr* (panel **b**), 0.042*mm/hr* (panel **c**), 0.05 *mm/hr* (panel **d**), and 0.052 *mm/hr* (panel **e**)), the resulting vegetation patterns (left column) showed the classic ‘clump–labyrinth-gap’ progression, culminating in the disappearance of patterns (i.e. only homogeneous vegetation observed) at the highest rainfall level (panel **e**). These vegetation patterns were similar to those obtained from the regular model/parameterisation in the absence of consumers (Fig. 6 c, top row). Grazers and predators (middle and right columns) did not show any boom-bust cycles or regularly organised travelling waves (contrast with Fig. 3 b). Grazer populations in particular always remained very low.

**Figure S.12:**
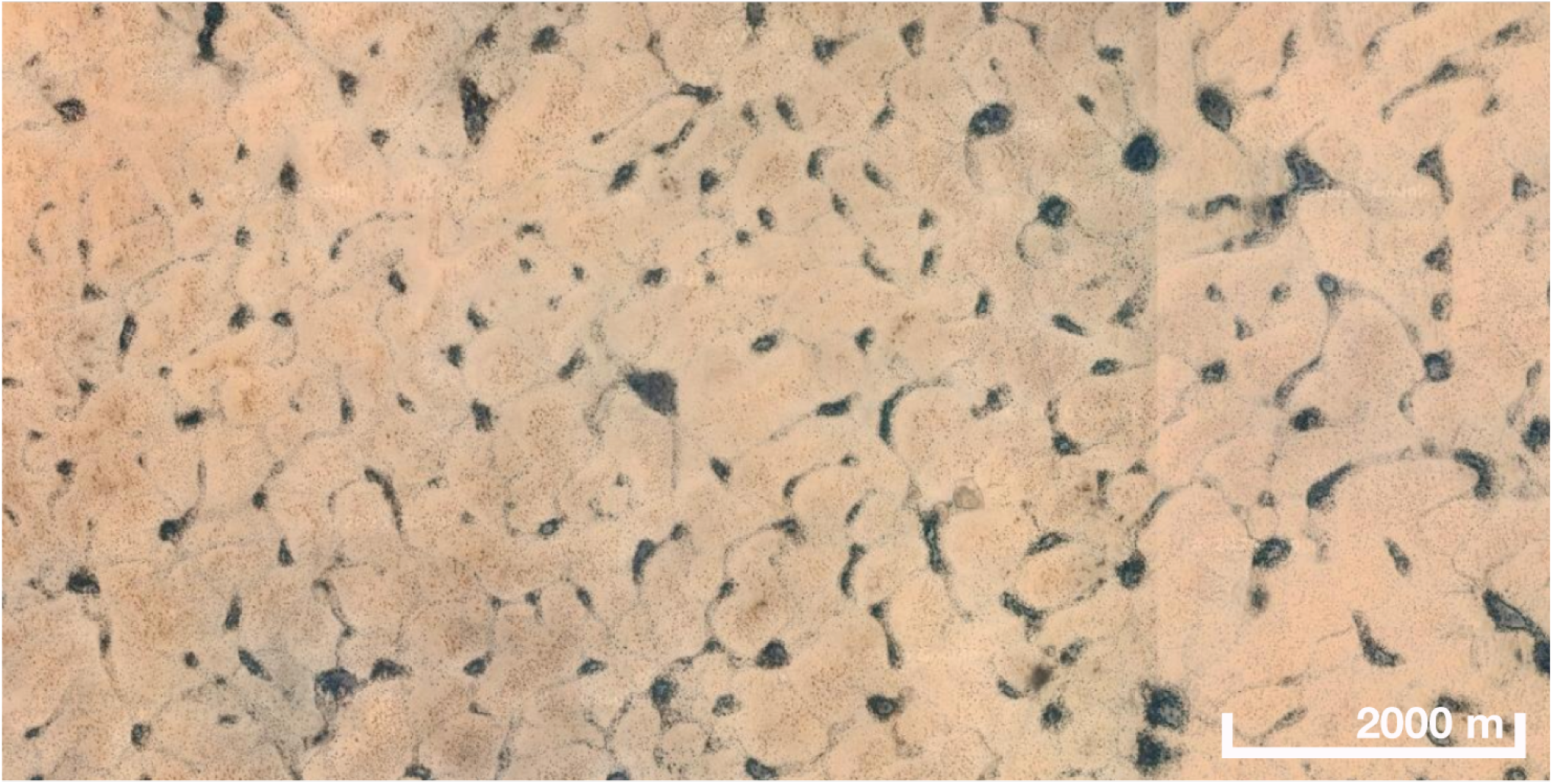
Satellite image of very large-scale dryland vegetation islands (clumps) near Tchintabaraden, Tenere, Niger (16.0325N, 5.6139E) (image credits: Google Earth) of a scale similar to that in our tri-trophic model (*dx* = 100 *m*). Vegetation-only ‘scale-dependent feedback’ (SDF) models parametrised for semi-arid ecosystems produce vegetation patterns in much smaller scales.

**Figure S.13:**
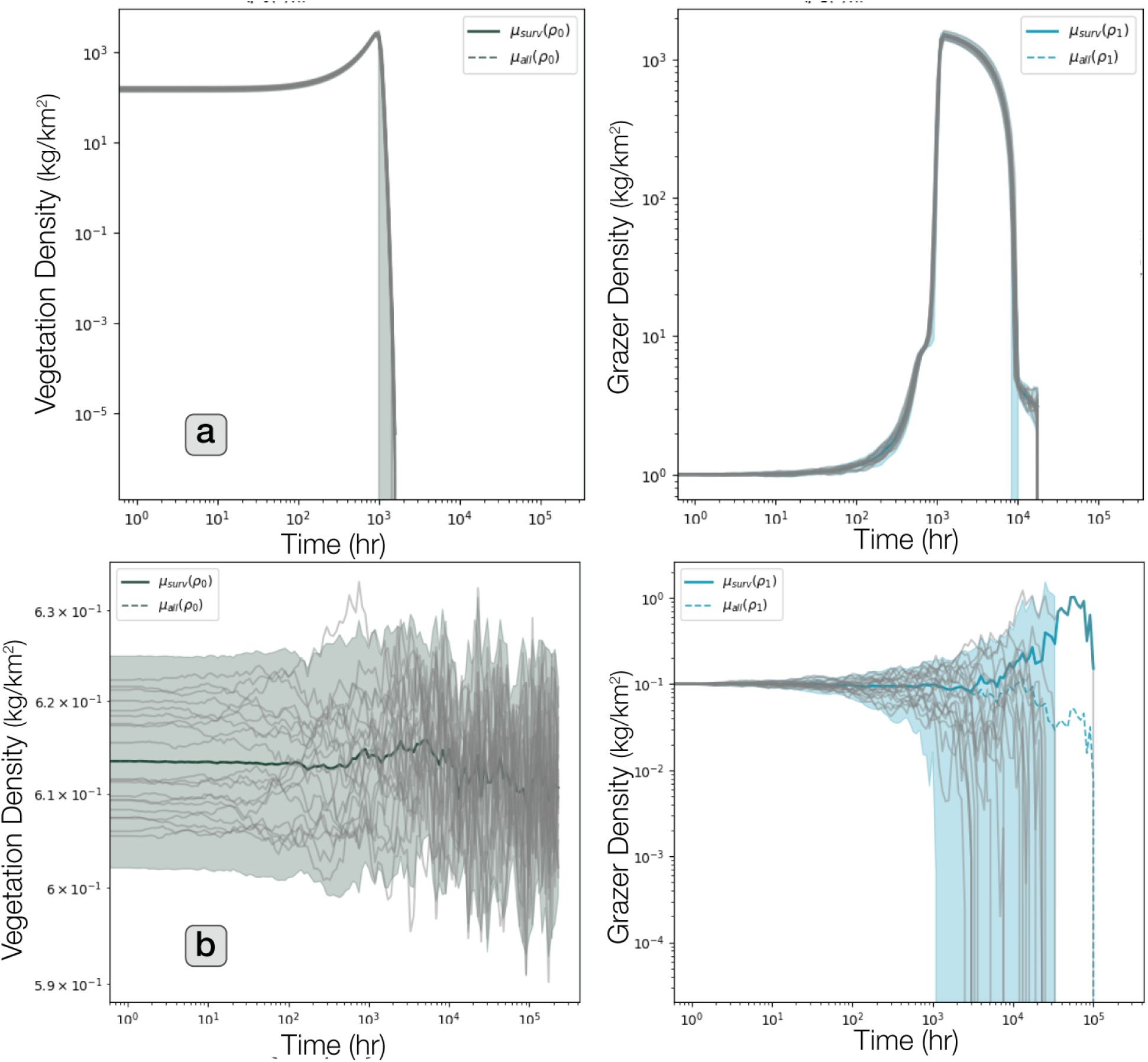
Simulations of the regular vegetation model (panel **a**) and the scale-free vegetation model (panel **b**) **without predators** showing how, without top-down regulation, the grazer population (right panels) grows faster than what the vegetation population (left panels) can sustain, leading to the collapse of the system.

**Figure S.14:**
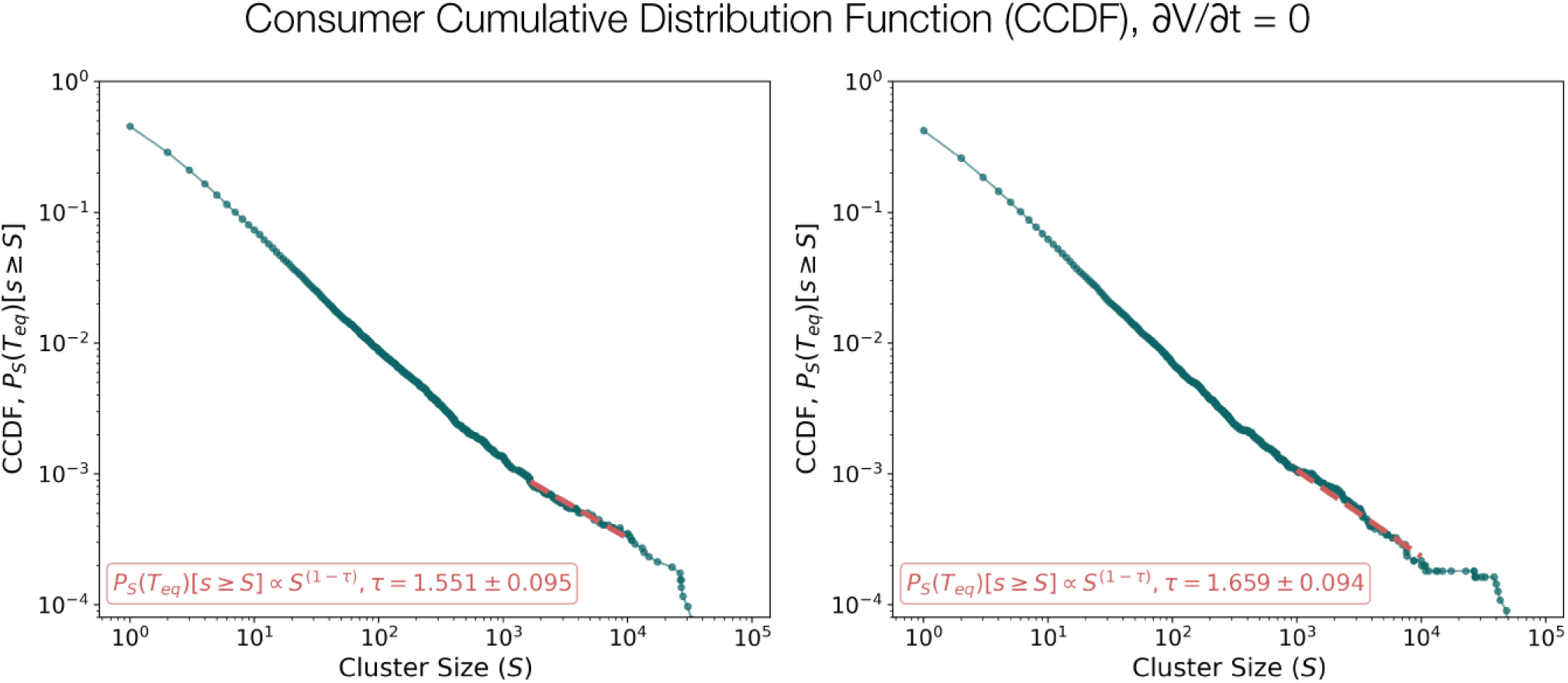
Distribution of patch size for grazers (left) and predators (right) measured using the complementary cumulative distribution function (CCDF) in a version of the model for which vegetation is constant (∂*V/*∂*t* = 0) and fixed to *V* ^⋆^ = 1 *kg/km*^2^.

**Figure S.15:**
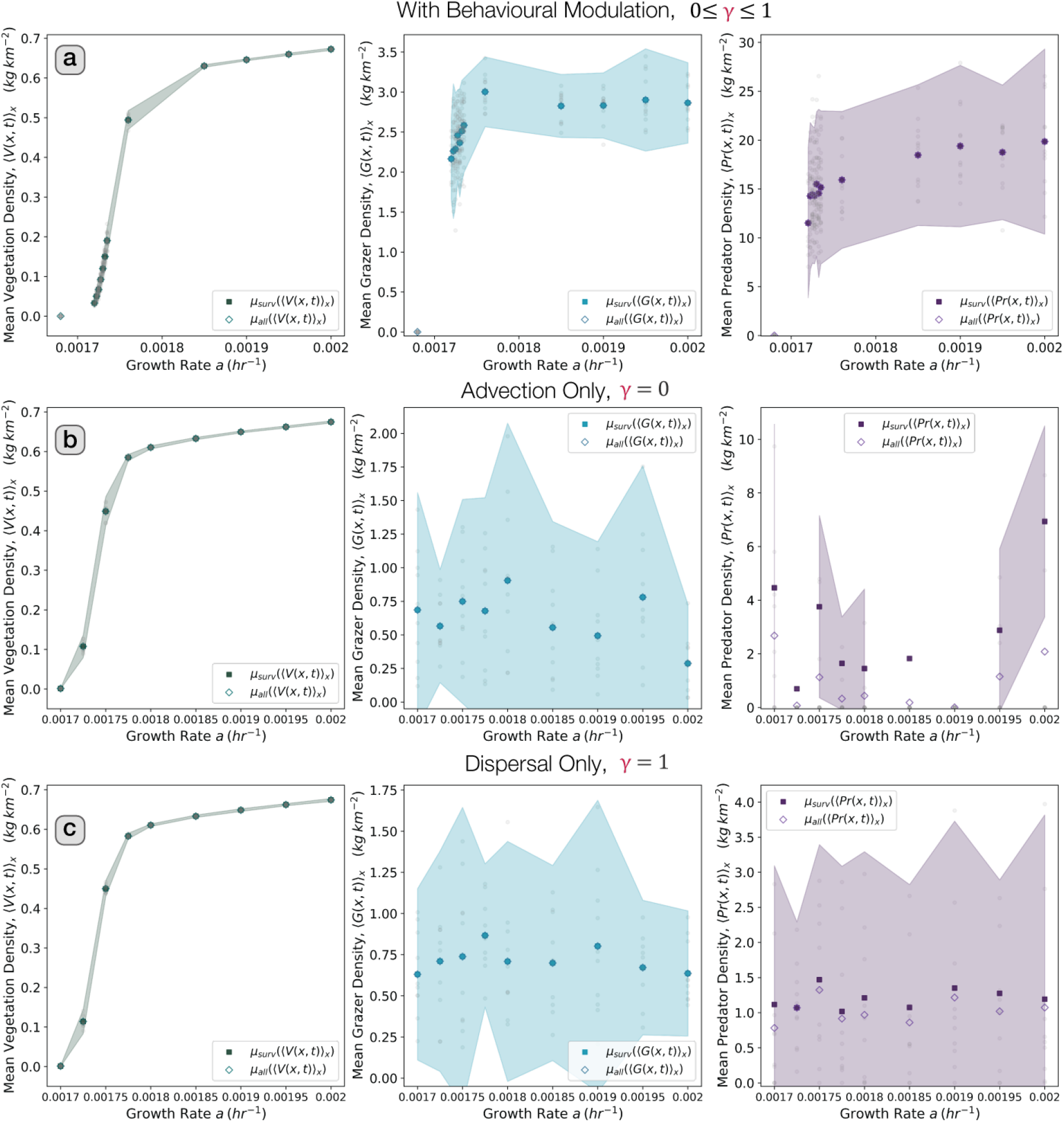
Phase diagram for the tri–trophic scale–free vegetation (SFV) model comprising of average densities for vegetation (left column), grazers (middle column), and predators (right column), showing the emergence of behaviourally modulated trophic cascades (Schmitz, Beckerman, and O’Brien 1997) in a *L* = 512 *×* 512 system. Solid points represent mean values across surviving replicates, hollow points represent mean values across all replicates (surviving or non-surviving), with shaded regions representing the 95% confidence interval. When behavioural modulation occurs (i.e. *γ* is determined as described in Eqs. 5a-5b), trophic cascades are observed as increases in resources correlate to increases in consumers (panel **a**); these trophic cascades disappear when behavioural modulation is eliminated by restricting movement to always advection (i.e. *γ* = 0, panel **b**) or always locally dispersing (i.e. *γ* = 1, panel **c**), as changes in vegetation do not materialize in changes in consumer densities, which remain constant, albeit at significantly lower densities across all vegetation growth rates.

**Table S.1:**
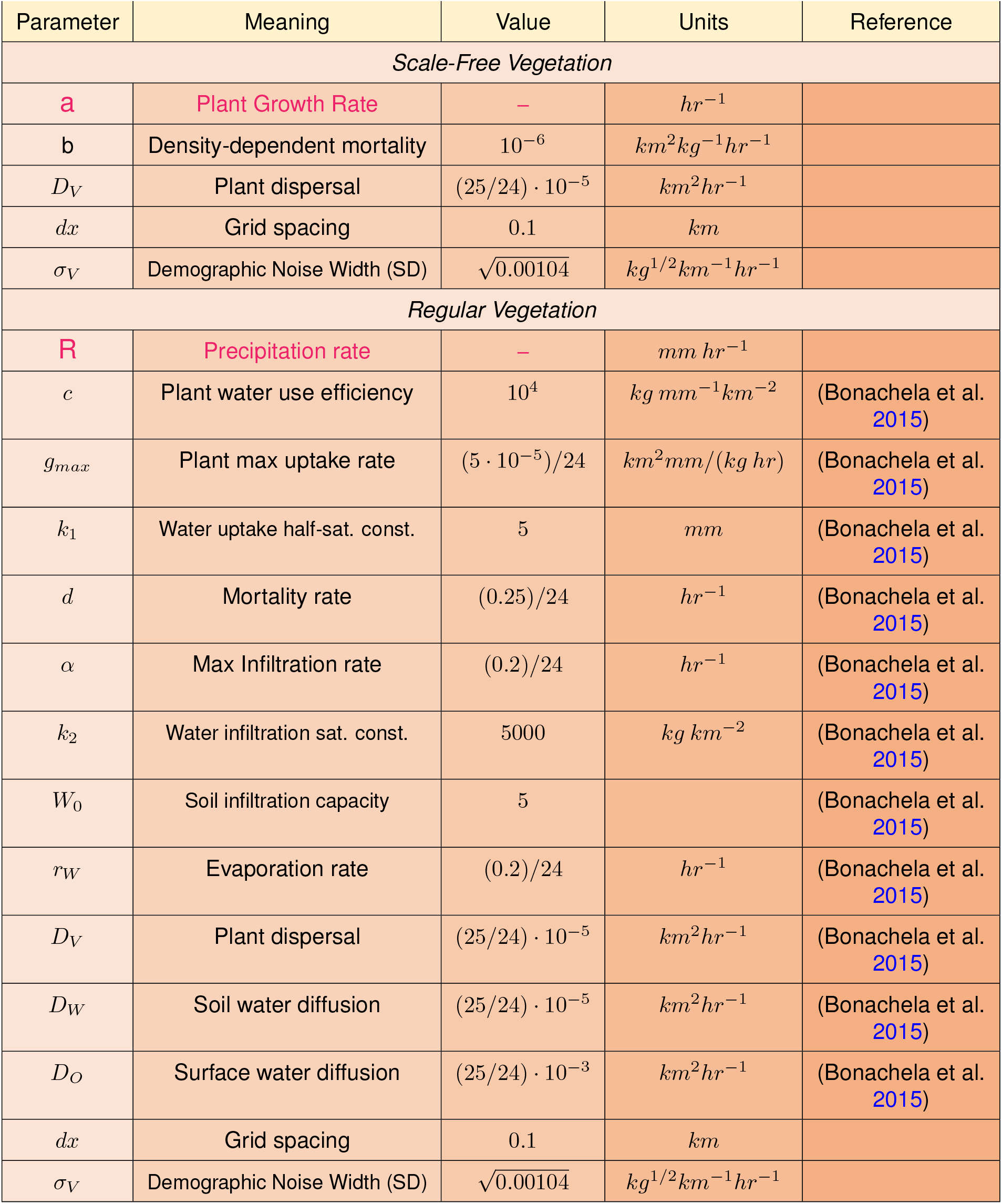
Parameterisation for vegetation parameters in the scale-free and regular vegetation models. These correspond to the variable *V* (*kg/km*^2^), representing vegetation biomass density (Eqs. (1a) and (1d)). Parameters in red reflect control parameters, i.e. the tuneable parameters that reflect the aridity of the system and that can be used to trigger desertification. Parameters for the vegetation-only SFV model were obtained from (Villa Martín et al. 2015), whereas those for the vegetation-only RV model were obtained from (Rietkerk et al. 2002) except for the diffusivity parameters, which were scaled up to ensure that patterns were resolved for our given grid spacing *dx*. This choice of vegetation diffusivity in the RV model was also used for the SFV model, for which the parameterσ_*V*_ defining the width (i.e. standard deviation) of vegetation demographic stochasticity, was set as 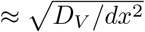.

| Parameter | Meaning | Value | Units | Reference |
|---|---|---|---|---|
| *Scale-Free Vegetation* |  |  |  |  |
| a | Plant Growth Rate | – | hr<sup>−1</sup> |  |
| b | Density-dependent mortality | 10<sup>−6</sup> | km<sup>2</sup>kg<sup>−1</sup>hr<sup>−1</sup> |  |
| D<sub>V</sub> | Plant dispersal | (25/24) · 10<sup>−5</sup> | km<sup>2</sup>hr<sup>−1</sup> |  |
| dx | Grid spacing | 0.1 | km |  |
| σ<sub>V</sub> | Demographic Noise Width (SD) | √0.00104 | kg<sup>1/2</sup>km<sup>−1</sup>hr<sup>−1</sup> |  |
| *Regular Vegetation* |  |  |  |  |
| R | Precipitation rate | – | mm hr<sup>−1</sup> |  |
| c | Plant water use efficiency | 10<sup>4</sup> | kg mm<sup>−1</sup>km<sup>−2</sup> | (Bonachela et al. 2015) |
| g<sub>max</sub> | Plant max uptake rate | (5 · 10<sup>−5</sup>)/24 | km<sup>2</sup>mm/(kg hr) | (Bonachela et al. 2015) |
| k<sub>1</sub> | Water uptake half-sat. const. | 5 | mm | (Bonachela et al. 2015) |
| d | Mortality rate | (0.25)/24 | hr<sup>−1</sup> | (Bonachela et al. 2015) |
| α | Max Infiltration rate | (0.2)/24 | hr<sup>−1</sup> | (Bonachela et al. 2015) |
| k<sub>2</sub> | Water infiltration sat. const. | 5000 | kg km<sup>−2</sup> | (Bonachela et al. 2015) |
| W<sub>0</sub> | Soil infiltration capacity | 5 |  | (Bonachela et al. 2015) |
| r<sub>W</sub> | Evaporation rate | (0.2)/24 | hr<sup>−1</sup> | (Bonachela et al. 2015) |
| D<sub>V</sub> | Plant dispersal | (25/24) · 10<sup>−5</sup> | km<sup>2</sup>hr<sup>−1</sup> | (Bonachela et al. 2015) |
| D<sub>W</sub> | Soil water diffusion | (25/24) · 10<sup>−5</sup> | km<sup>2</sup>hr<sup>−1</sup> | (Bonachela et al. 2015) |
| D<sub>O</sub> | Surface water diffusion | (25/24) · 10<sup>−3</sup> | km<sup>2</sup>hr<sup>−1</sup> | (Bonachela et al. 2015) |
| dx | Grid spacing | 0.1 | km |  |
| σ<sub>V</sub> | Demographic Noise Width (SD) | √0.00104 | kg<sup>1/2</sup>km<sup>−1</sup>hr<sup>−1</sup> |  |

**Table S.2:**
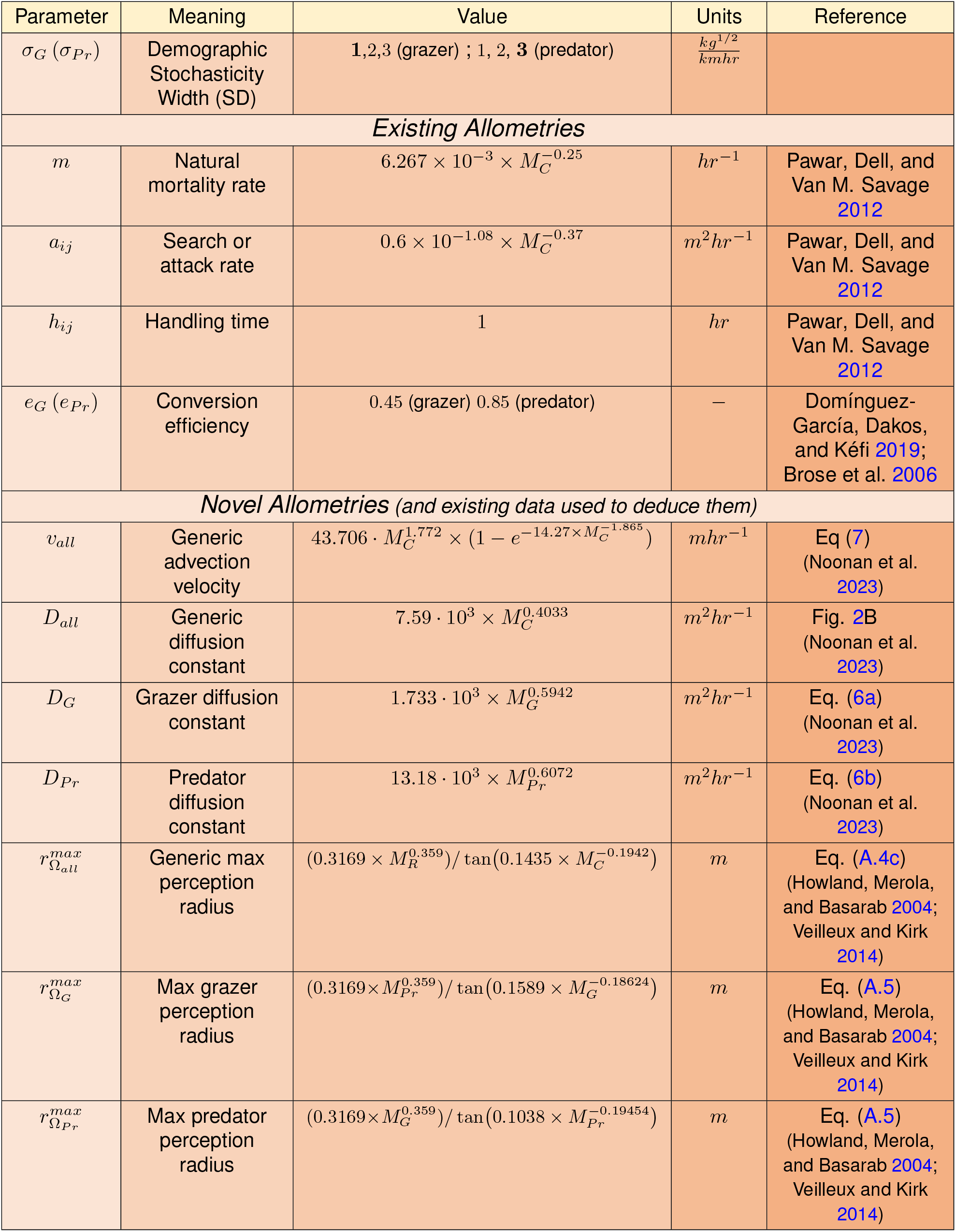
Complete allometric parameterisation for consumers. These correspond to the consumer variables, i.e. grazer biomass density (*G* in *kg/km*^2^, Eq (2)) and predator biomass density (*Pr* in *kg/km*^2^, Eq (3))). *M*_*C*_ refers to the bodymass of a generic consumer (in kg), *M*_*R*_ refers to body mass of a generic resource (in kg), while *M*_*G*_ (= 20 *kg* in simulations) and *M*_*Pr*_ (= 100 *kg* in simulations) refer to bodymasses of generic grazers and predators respectively (in kg). The parameter for the width of consumer demographic noise (**bold** font in figures) correspond to 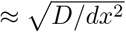

| Parameter | Meaning | Value | Units | Reference |
|---|---|---|---|---|
| σ<sub>G</sub> (σ<sub>Pr</sub>) | Demographic Stochasticity Width (SD) | **1**,2,3 (grazer) ; 1, 2, **3** (predator) | $\frac{kg^{1/2}}{kmhr}$ |  |
| *Existing Allometries* |  |  |  |  |
| *m* | Natural mortality rate | 6.267 × 10<sup>−3</sup> × *M*<sub>*C*</sub><sup>−0.25</sup> | *hr*<sup>−1</sup> | Pawar, Dell, and Van M. Savage 2012 |
| *a*<sub>*ij*</sub> | Search or attack rate | 0.6 × 10<sup>−1.08</sup> × *M*<sub>*C*</sub><sup>−0.37</sup> | *m*<sup>2</sup>*hr*<sup>−1</sup> | Pawar, Dell, and Van M. Savage 2012 |
| *h*<sub>*ij*</sub> | Handling time | 1 | *hr* | Pawar, Dell, and Van M. Savage 2012 |
| *e*<sub>*G*</sub> (*e*<sub>*Pr*</sub>) | Conversion efficiency | 0.45 (grazer) 0.85 (predator) | − | Domínguez-García, Dakos, and Kéfi 2019; Brose et al. 2006 |
| *Novel Allometries* *(and existing data used to deduce them)* |  |  |  |  |
| *v*<sub>*all*</sub> | Generic advection velocity | 43.706 · *M*<sub>*C*</sub><sup>1.772</sup> × (1 − *e*<sup>−14.27×*M*<sub>*C*</sub><sup>−1.865</sup></sup>) | *mhr*<sup>−1</sup> | Eq (7) (Noonan et al. 2023) |
| *D*<sub>*all*</sub> | Generic diffusion constant | 7.59 · 10<sup>3</sup> × *M*<sub>*C*</sub><sup>0.4033</sup> | *m*<sup>2</sup>*hr*<sup>−1</sup> | Fig. 2B (Noonan et al. 2023) |
| *D*<sub>*G*</sub> | Grazer diffusion constant | 1.733 · 10<sup>3</sup> × *M*<sub>*G*</sub><sup>0.5942</sup> | *m*<sup>2</sup>*hr*<sup>−1</sup> | Eq. (6a) (Noonan et al. 2023) |
| *D*<sub>*Pr*</sub> | Predator diffusion constant | 13.18 · 10<sup>3</sup> × *M*<sub>*Pr*</sub><sup>0.6072</sup> | *m*<sup>2</sup>*hr*<sup>−1</sup> | Eq. (6b) (Noonan et al. 2023) |
| *r*<sup>*max*</sup><sub>Ω<sub>*all*</sub></sub> | Generic max perception radius | (0.3169 × *M*<sub>*R*</sub><sup>0.359</sup>)/ tan(0.1435 × *M*<sub>*C*</sub><sup>−0.1942</sup>) | *m* | Eq. (A.4c) (Howland, Merola, and Basarab 2004; Veilleux and Kirk 2014) |
| *r*<sup>*max*</sup><sub>Ω<sub>*G*</sub></sub> | Max grazer perception radius | (0.3169×*M*<sub>*Pr*</sub><sup>0.359</sup>)/ tan(0.1589 × *M*<sub>*G*</sub><sup>−0.18624</sup>) | *m* | Eq. (A.5) (Howland, Merola, and Basarab 2004; Veilleux and Kirk 2014) |
| *r*<sup>*max*</sup><sub>Ω<sub>*Pr*</sub></sub> | Max predator perception radius | (0.3169×*M*<sub>*G*</sub><sup>0.359</sup>)/ tan(0.1038 × *M*<sub>*Pr*</sub><sup>−0.19454</sup>) | *m* | Eq. (A.5) (Howland, Merola, and Basarab 2004; Veilleux and Kirk 2014) |

## References

Biernacki, C., G. Celeux, and G. Govaert (July 2000). “Assessing a mixture model for clustering with the integrated completed likelihood”. In: IEEE Transactions on Pattern Analysis and Machine Intelligence 22.7, pp. 719–725. ISSN: 1939-3539.

Bjørnstad, Ottar N. and Jordi Bascompte (2001). “Synchrony and Second-Order Spatial Correlation in Host-Parasitoid Systems”. In: Journal of Animal Ecology 70.6, pp. 924–933. ISSN: 0021-8790.

Bonachela, Juan A. et al. (Feb. 6, 2015). “Termite mounds can increase the robustness of dryland ecosystems to climatic change”. In: Science 347.6222. Publisher: American Association for the Advancement of Science, pp. 651–655.

Born, Max and Emil Wolf (2019). Principles of Optics: 60th Anniversary Edition. 7th ed. Cambridge University Press.

Brose, U. et al. (Sept. 2008). “Foraging theory predicts predator-prey energy fluxes”. In: Journal of Animal Ecology 77.5, pp. 1072–1078. ISSN: 00218790, 13652656.

Caretta, Martina Angela et al. (2022). “Water”. In: Climate Change 2022: Impacts, Adaptation and Vulnerability. Ed. by Hans-Otto Pörtner et al. Intergovernmental Panel on Climate Change (IPCC), pp. 551–712.

Chen, Zhuoxue, Koustav Halder, and Juan A. Bonachela (2024). “Vegetation clump size and number as indicators for alternative stable states in semi-arid ecosystems”. In: under review.

Dai, Lei, Kirill S. Korolev, and Jeff Gore (Apr. 2013). “Slower recovery in space before collapse of connected populations”. In: Nature 496.7445, pp. 355–358. ISSN: 0028-0836, 1476-4687.

Farías, Ariel A. et al. (2021). “Species interactions across trophic levels mediate rainfall effects on dryland vegetation dynamics”. en. In: Ecological Monographs 91.2, e01441. ISSN: 1557-7015.

Gaynor, Kaitlyn M. et al. (Apr. 1, 2019). “Landscapes of Fear: Spatial Patterns of Risk Perception and Response”. In: Trends in Ecology & Evolution 34.4, pp. 355–368. ISSN: 0169-5347.

Ge, Zhenpeng and Quan-Xing Liu (2022). “Foraging behaviours lead to spatiotemporal self-similar dynamics in grazing ecosystems”. In: Ecology Letters 25.2, pp. 378–390. ISSN: 1461-0248. eprint: https://onlinelibrary.wiley.com/doi/pdf/10.1111/ele.13928.

Grünzweig, José M. et al. (Aug. 2022). “Dryland mechanisms could widely control ecosystem functioning in a drier and warmer world”. en. In: Nature Ecology & Evolution 6.8, pp. 1064–1076. ISSN: 2397-334X.

Guill, Christian, Janne Hülsemann, and Toni Klauschies (2021). “Self-organised pattern formation increases local diversity in metacommunities”. en. In: Ecology Letters 24.12, pp. 2624–2634. ISSN: 1461-0248.

Guttal, Vishwesha and Ciriyam Jayaprakash (May 2008). “Changing skewness: an early warning signal of regime shifts in ecosystems”. In: Ecology Letters 11.5, pp. 450–460. ISSN: 1461-023X, 1461-0248.

Hinrichsen, Haye (Nov. 2000). “Non-equilibrium critical phenomena and phase transitions into absorbing states”. In: Advances in Physics 49.7, pp. 815–958. ISSN: 0001-8732, 1460-6976.

Hirt, Myriam R. et al. (July 17, 2017). “A general scaling law reveals why the largest animals are not the fastest”. In: Nature Ecology & Evolution 1.8, pp. 1116–1122. ISSN: 2397-334X.

Hirt, Myriam R. et al. (Sept. 2018). “Bridging Scales: Allometric Random Walks Link Movement and Biodiversity Research”. In: Trends in Ecology & Evolution 33.9, pp. 701–712. ISSN: 01695347.

Holling, C. S. (July 1959). “Some Characteristics of Simple Types of Predation and Parasitism”. In: The Canadian Entomologist 91.7, pp. 385–398. ISSN: 0008-347X, 1918-3240.

Holling, C. S. (1973). “Resilience and Stability of Ecological Systems”. In: Annual Review of Ecology and Systematics 4. Publisher: Annual Reviews, pp. 1–23. ISSN: 0066-4162.

Hoshen, J. and R. Kopelman (Oct. 1976). “Percolation and cluster distribution. I. Cluster multiple labeling technique and critical concentration algorithm”. In: Physical Review B 14.8. Publisher: American Physical Society, pp. 3438–3445.

Howland, Howard C, Stacey Merola, and Jennifer R Basarab (Aug. 2004). “The allometry and scaling of the size of vertebrate eyes”. In: Vision Research 44.17, pp. 2043–2065. ISSN: 00426989.

“Desertification” (2022a). In: Climate Change and Land: IPCC Special Report on Climate Change, Desertification, Land Degradation, Sustainable Land Management, Food Security, and Greenhouse Gas Fluxes in Terrestrial Ecosystems. Ed. by Intergovernmental Panel on Climate Change. Cambridge: Cambridge University Press, pp. 249–344. ISBN: 978-1-00-915798-8.

“Land degradation” (2022b). In: Climate Change and Land: IPCC Special Report on Climate Change, Desertification, Land Degradation, Sustainable Land Management, Food Security, and Greenhouse Gas Fluxes in Terrestrial Ecosystems. Ed. by Intergovernmental Panel on Climate Change. Cambridge: Cambridge University Press, pp. 345–436. ISBN: 978-1-00-915798-8.

Johnson, Devin S. et al. (2008). “Continuous-Time Correlated Random Walk Model for Animal TELemetry Data”. In: Ecology 89.5, pp. 1208–1215. ISSN: 1939-9170. eprint: https://onlinelibrary.wiley.com/doi/pdf/10.1890/07-1032.1.

Kéfi, Sonia et al. (Sept. 13, 2007). “Spatial vegetation patterns and imminent desertification in Mediterranean arid ecosystems”. In: Nature 449, pp. 213–217.

Kéfi, Sonia et al. (2012). “More than a meal… integrating non-feeding interactions into food webs”. In: Ecology Letters 15.4, pp. 291–300. ISSN: 1461-0248. eprint: https://onlinelibrary.wiley.com/doi/pdf/10.1111/j.1461-0248.2011.01732.x.

Kéfi, Sonia et al. (Mar. 21, 2014). “Early Warning Signals of Ecological Transitions: Methods for Spatial Patterns”. In: PLOS ONE 9.3. Publisher: Public Library of Science, e92097. ISSN: 1932-6203.

Lejeune, O., M. Tlidi, and P. Couteron (July 2002). “Localized vegetation patches: A self-organized response to resource scarcity”. In: Physical Review E 66.1, p. 010901.

Liebhold, Andrew, Walter D. Koenig, and Ottar N. Bjørnstad (Dec. 2004). “Spatial Synchrony in Population Dynamics*”. en. In: Annual Review of Ecology, Evolution, and Systematics 35.Volume 35, 2004, pp. 467–490. ISSN: 1543-592X, 1545-2069.

Maestre, Fernando T. et al. (Nov. 25, 2022). “Grazing and ecosystem service delivery in global drylands”. In: Science 378.6622, pp. 915–920.

Maimaiti, Yimamu, Wenbin Yang, and Jianhua Wu (Apr. 2022). “Turing instability and coexistence in an extended Klausmeier model with nonlocal grazing”. In: Nonlinear Analysis: Real World Applications 64, p. 103443. ISSN: 1468-1218.

Manor, Alon and Nadav M. Shnerb (Aug. 2008). “Facilitation, competition, and vegetation patchiness: From scale free distribution to patterns”. In: Journal of Theoretical Biology 253.4, pp. 838–842. ISSN: 0022-5193.

Marescot, Lucile et al. (Jan. 2025). “A forecasting model for desert locust presence during recession period, using real-time satellite imagery”. In: Remote Sensing Applications: Society and Environment 37, p. 101497. ISSN: 2352-9385.

Morris, Timothy and Mike Letnic (May 2017). “Removal of an apex predator initiates a trophic cascade that extends from herbivores to vegetation and the soil nutrient pool”. In: Proceedings of the Royal Society B: Biological Sciences 284.1854, p. 20170111. ISSN: 0962-8452.

Muñoz, M. A. et al. (May 1999). “Avalanche and spreading exponents in systems with absorbing states”. In: Physical Review. E, Statistical Physics, Plasmas, Fluids, and Related Interdisciplinary Topics 59.5, pp. 6175–6179. ISSN: 1063-651X.

Noonan, Michael J. et al. (Jan. 3, 2023). The search behavior of terrestrial mammals.

Paradis, E. et al. (1999). “Dispersal and spatial scale affect synchrony in spatial population dynamics”. en. In: Ecology Letters 2.2, pp. 114–120. ISSN: 1461-0248.

Pavey, Chris R. et al. (May 2017). “The role of refuges in the persistence of Australian dryland mammals”. In: Biological Reviews 92.2, pp. 647–664. ISSN: 1464-7931, 1469-185X.

Pawar, Samraat, Anthony I. Dell, and Van M. Savage (June 28, 2012). “Dimensionality of consumer search space drives trophic interaction strengths”. In: Nature 486.7404, pp. 485–489. ISSN: 0028-0836, 1476-4687.

Pichon, Benoît et al. (2025). “Grazing Modulates the Multiscale Spatial Structure of Dryland Vegetation”. en. In: Global Change Biology 31.7, e70345. ISSN: 1365-2486.

Ranta, Esa et al. (1997). “The Moran Effect and Synchrony in Population Dynamics”. In: Oikos 78.1, pp. 136–142. ISSN: 0030-1299.

Rao, Xiao et al. (2024). “Population asynchrony within and between trophic levels have contrasting effects on consumer community stability in a subtropical lake”. en. In: Journal of Animal Ecology 93.10, pp. 1593–1605. ISSN: 1365-2656.

Reynolds, James F. et al. (May 11, 2007a). “Global Desertification: Building a Science for Dryland Development”. In: Science 316.5826, pp. 847–851. ISSN: 0036-8075, 1095-9203.

Reynolds, James F. et al. (May 11, 2007b). “Global Desertification: Building a Science for Dryland Development”. In: Science 316.5826, pp. 847–851. ISSN: 0036-8075, 1095-9203.

Rietkerk, M. and J. van de Koppel (1997). “Alternate Stable States and Threshold Effects in Semi-Arid Grazing Systems”. In: Oikos 79.1, pp. 69–76. ISSN: 0030-1299.

Rietkerk, Max and Johan van de Koppel (Mar. 1, 2008). “Regular pattern formation in real ecosystems”. In: Trends in Ecology & Evolution 23.3, pp. 169–175. ISSN: 0169-5347.

Rietkerk, Max et al. (Oct. 2002). “Self-organization of vegetation in arid ecosystems”. In: The American Naturalist 160.4, pp. 524–530. ISSN: 1537-5323.

Rietkerk, Max et al. (Sept. 24, 2004). “Self-Organized Patchiness and Catastrophic Shifts in Ecosystems”. In: Science 305.5692, pp. 1926–1929. ISSN: 0036-8075, 1095-9203.

Riginos, Corinna (2015). “Climate and the landscape of fear in an African savanna”. en. In: Journal of Animal Ecology 84.1, pp. 124–133. ISSN: 1365-2656.

Rosenzweig, Cynthia et al. (Mar. 4, 2014). “Assessing agricultural risks of climate change in the 21st century in a global gridded crop model intercomparison”. In: Proceedings of the National Academy of Sciences 111.9. Publisher: Proceedings of the National Academy of Sciences, pp. 3268–3273.

Scanlon, Todd M. et al. (Sept. 2007). “Positive feedbacks promote power-law clustering of Kalahari vegetation”. In: Nature 449.7159. Number: 7159 Publisher: Nature Publishing Group, pp. 209–212. ISSN: 1476-4687.

Schmitz, Oswald J., Andrew P. Beckerman, and Kathleen M. O’Brien (1997). “Behaviorally Mediated Trophic Cascades: Effects of Predation Risk on Food Web Interactions”. en. In: Ecology 78.5, pp. 1388–1399. ISSN: 1939-9170.

Schneider, Florian D. and Sonia Kéfi (June 2016). “Spatially heterogeneous pressure raises risk of catastrophic shifts”. In: Theoretical Ecology 9.2, pp. 207–217. ISSN: 1874-1738, 1874-1746.

Siero, Eric et al. (Mar. 2019). “Grazing Away the Resilience of Patterned Ecosystems”. In: The American Naturalist 193.3, pp. 472–480. ISSN: 0003-0147, 1537-5323.

Silva, M. (Feb. 20, 1998). “Allometric Scaling of Body Length: Elastic or Geometric Similarity in Mammalian Design”. In: Journal of Mammalogy 79.1, pp. 20–32. ISSN: 1545-1542, 0022-2372.

Singha, Joydeep, Hannes Uecker, and Ehud Meron (Dec. 2025). “Traveling vegetation–herbivore waves may sustain ecosystems threatened by droughts and population growth”. In: Physica D: Nonlinear Phenomena 483, p. 134914. ISSN: 0167-2789.

Skalski, Garrick T. and James F. Gilliam (Mar. 2003). “A Diffusion-Based Theory of Organism Dispersal in Heterogeneous Populations.” In: The American Naturalist 161.3. Publisher: The University of Chicago Press, pp. 441–458. ISSN: 0003-0147.

Tarnita, Corina E. et al. (Jan. 2017). “A theoretical foundation for multi-scale regular vegetation patterns”. In: Nature 541.7637. Number: 7637 Publisher: Nature Publishing Group, pp. 398–401. ISSN: 1476-4687.

The State of the World’s Forests (2020). FAO and UNEP. ISBN: 978-92-5-132419-6.

Turchin, Peter (1998). Quantitative Analysis of Movement: Measuring and Modeling Population Redistribution in Animals and Plants. Google-Books-ID: ZbdmQgAACAAJ. Sinauer. 396 pp. ISBN: 978-0-87893-847-6.

Tyson, R.C., J.B. Wilson, and W.D. Lane (May 2011). “Beyond diffusion: Modelling local and long-distance dispersal for organisms exhibiting intensive and extensive search modes”. In: Theoretical Population Biology 79.3, pp. 70–81. ISSN: 00405809.

Vardaro, Jessica A. Castillo et al. (2021). “Resource availability and heterogeneity shape the self-organisation of regular spatial patterning”. In: Ecology Letters 24, pp. 1880–1891.

Veilleux, Carrie C. and E. Christopher Kirk (2014). “Visual Acuity in Mammals: Effects of Eye Size and Ecology”. In: Brain, Behavior and Evolution 83.1, pp. 43–53. ISSN: 0006-8977, 1421-9743.

Villa Martín, Paula et al. (Apr. 14, 2015). “Eluding catastrophic shifts”. In: Proceedings of the National Academy of Sciences 112.15, E1828–E1836. ISSN: 0027-8424, 1091-6490.

Vinh, Nguyen Xuan, Julien Epps, and James Bailey (Dec. 2010). “Information Theoretic Measures for Clusterings Comparison: Variants, Properties, Normalization and Correction for Chance”. In: J. Mach. Learn. Res. 11, pp. 2837–2854. ISSN: 1532-4435.

Visser, Marcel E. and Phillip Gienapp (June 2019). “Evolutionary and demographic consequences of phenological mismatches”. en. In: Nature Ecology & Evolution 3.6, pp. 879–885. ISSN: 2397-334X.

Walter, Jonathan A. et al. (July 2017). “The geography of spatial synchrony”. en. In: Ecology Letters 20.7. Ed. by Bernd Blasius, pp. 801–814. ISSN: 1461-023X, 1461-0248.

West, Geoffrey B., James H. Brown, and Brian J. Enquist (Aug. 1999). “A general model for the structure and allometry of plant vascular systems”. In: Nature 400.6745, pp. 664–667. ISSN: 0028-0836, 1476-4687.

Yodzis, Peter and S. Innes (1992). “Body Size and Consumer-Resource Dynamics”. In: The American Naturalist 139.6, pp. 1151–1175.

Zhao, Jianlong, Yong Sang, and Fuhai Duan (2019). “The state of the art of two-dimensional digital image correlation computational method”. In: Engineering Reports 1.2. e12038 ENG-2019-04-0110.R2, e12038. eprint: https://onlinelibrary.wiley.com/doi/pdf/10.1002/eng2.12038.

## Supplementary References

Clauset, Aaron, Cosma Rohilla Shalizi, and M. E. J. Newman (Nov. 4, 2009). “Power-Law Distributions in Empirical Data”. In: SIAM Review 51.4, pp. 661–703. ISSN: 0036-1445, 1095-7200.

Dornic, Ivan, Hugues Chaté, and Miguel A Muñoz (2005). “Integration of Langevin equations with multiplicative noise and the viability of field theories for absorbing phase transitions”. In: Physical review letters 94.10, p. 100601.

E. Ewing Richard and Hong Wang (Mar. 2001). “A summary of numerical methods for time-dependent advection-dominated partial differential equations”. In: Journal of Computational and Applied Mathematics. Numerical Analysis 2000. Vol. VII: Partial Differential Equations 128.1, pp. 423–445. ISSN: 0377-0427.

Fried, D. L. (Oct. 1966). “Optical Resolution Through a Randomly Inhomogeneous Medium for Very Long and Very Short Exposures”. EN. In: JOSA 56.10, pp. 1372–1379.

Frigo, M. and S.G. Johnson (Feb. 2005). “The Design and Implementation of FFTW3”. In: Proceedings of the IEEE 93.2, pp. 216–231. ISSN: 0018-9219.

Hardy, John W. (1998). Adaptive optics for astronomical telescopes. eng. Open Library ID: OL660197M. New York: Oxford University Press. ISBN: 9780195090192.

LeVeque, Randall J. (2002). Finite Volume Methods for Hyperbolic Problems. Cambridge Texts in Applied Mathematics. Cambridge: Cambridge University Press. ISBN: 978-0-521-00924-9.

Mann, H. B. and D. R. Whitney (1947). “On a Test of Whether one of Two Random Variables is Stochastically Larger than the Other”. In: The Annals of Mathematical Statistics 18.1, pp. 50–60. ISSN: 0003-4851.

Scott, David W. (Dec. 1, 1979). “On optimal and data-based histograms”. In: Biometrika 66.3, pp. 605–610. ISSN: 0006-3444.

Vinh, Nguyen Xuan, Julien Epps, and James Bailey (June 14, 2009). “Information theoretic measures for clusterings comparison: is a correction for chance necessary?” In: Proceedings of the 26th Annual International Conference on Machine Learning. ICML ‘09. New York, NY, USA: Association for Computing Machinery, pp. 1073–1080. ISBN: 9781605585161.

Yoder, Paul R. and Daniel Vukobratovich (2011). Field Guide to Binoculars and Scopes. Google-Books-ID: uR2YZwEACAAJ. SPIE. 140 pp. ISBN: 978-0-8194-8649-3.

## Supplementary References

Brose, Ulrich et al. (Oct. 2006). “Consumer–resource body-size relationships in natural food webs”. In: Ecology 87.10, pp. 2411–2417. ISSN: 0012-9658.

Domínguez-García, Virginia, Vasilis Dakos, and Sonia Kéfi (Dec. 17, 2019). “Unveiling dimensions of stability in complex ecological networks”. In: Proceedings of the National Academy of Sciences 116.51, pp. 25714–25720. ISSN: 0027-8424, 1091-6490.

Noonan, Michael J. et al. (Jan. 3, 2023). The search behavior of terrestrial mammals.

Strogatz, Steven H. (2015). Nonlinear dynamics and chaos: with applications to physics, biology, chemistry, and engineering. Second edition. OCLC: ocn842877119. Boulder, CO: Westview Press, a member of the Perseus Books Group. 513 pp. ISBN: 978-0-8133-4910-7.

